# Analyzing the interaction of human ACE2 and RBD of spike protein of SARS-CoV-2 in perspective of Omicron variant

**DOI:** 10.1101/2021.12.23.473991

**Authors:** Arijit Samanta, Syed Sahajada Mahafujul Alam, Safdar Ali, Mehboob Hoque

**Author notes:** Correspondence Dr Mehboob Hoque, Department of Biological Sciences, Aliah University, Kolkata 700160, India.

## Abstract

The newly identified Omicron (B.1.1.529) variant of Severe Acute Respiratory Syndrome Voronavirus 2 (SARS-CoV-2) has steered concerns across the world due to the possession of large number of mutations leading to high infectivity and vaccine escape potential. The Omicron variant houses 32 mutations in S protein alone. The viral infectivity is determined mainly by the ability of spike (S) protein receptor binding domain (RBD) to bind to the human Angiotensin I Converting Enzyme 2 (hACE2) receptor. In this paper, the interaction of the RBDs of SARS-CoV-2 variants with hACE2 was analyzed by using protein-protein docking and compared with the novel Omicron variant. Our findings reveal that the Omicron RBD interacts strongly with hACE2 receptor via unique amino acid residues as compared to the Wuhan and many other variants. However, the interacting residues of RBD are found to be the same in Lamda (C.37) variant. These unique binding of Omicron RBD with hACE2 suggests an increased potential of infectivity and vaccine evasion potential of the new variant. The evolutionary drive of the SARS-CoV-2 may not be exclusively driven by RBD variants but surely provides for the platform for emergence of new variants.

## 1. Introduction

For the last two years the World has been witnessing an unprecedented pandemic of the coronavirus disease 2019 (COVID-19) caused by Severe Acute Respiratory Syndrome Coronavirus 2 (SARS-CoV-2). The global concerted efforts and significantly fast scientific advancements resulted in reducing disease fatality and infection rates with record number of vaccination. Despite the achievement in such a short span of time, there remains ever rising challenges and uncertainties posed by the emergence of new mutated variants of SARS-CoV-2 The World Health Organization (WHO) has been regularly monitoring the evolution of SARS-CoV-2 after declaring it a pandemic in February 2020. It has classified the emerging variants of SARS-CoV-2 into variants of concern (VOCs), variants of interest (VOIs) and variant under monitoring (VUMs). According to the latest update on December 13, 2021 by WHO, a total of 5 variants have been listed under VOCs, 2 variants under VOIs, 7 variants under VUM and 12 formerly monitored variants (https://www.who.int/en/activities/tracking-SARS-CoV-2-variants/). The VOCs show high transmissibility, highly virulence, decrease in the efficacy and effectiveness of public health and social measures or available diagnostics, vaccines and therapeutics. Most recently, on November 26, 2021, a highly mutated new variant of SARS-CoV-2, named Omicron (B.1.1.529), first detected in South Africa, has been classified as one of the VOCs by the WHO (https://www.who.int/). Occurrence of Omicron variant has been detected in 77 countries till December 20, 2021 (https://www.gisaid.org/). There have been four variants already designated as VOCs including Alpha (B.1.1.7), Beta (B.1.351), Gamma (P.1), and Delta (B.1.617.2).

The currently available data suggest that the Omicron has more than 50 mutations, including 32 on the Spike (S) protein alone, including amino acid substitution, deletion, and insertion (Poudel et al., 2022). Interestingly some of these mutations and substitutions are present in the Receptor Binding Domain (RBD). These mutations are attributed for the increased rate of infection and immune evasion capacity (Quarleri et al., 2021)(Poudel et al., 2022)(Nasrin and Ali, 2021). The SARS-RBD of S protein interacts to the extracellular peptidase domain of human Angiotensin I Converting Enzyme 2 (hACE2) receptor to mediate cell entrance. Therefore, the mutations at the RBD are expected to alter the binding efficiency with hACE2, influencing the infectivity. The current study has been designed to study the binding affinity and interaction pattern of RBD of omicron and other variants of SARS-CoV-2 with hACE2 receptor.

## 2. Materials and methods

### 2.1. Retrieval of the hACE2 and SARS-CoV-2 S protein RBD complex structure

The RDB region of B chain of S protein mainly interacts with A chain of hACE2 during infection. Therefore, only the RDB region of B chain of S protein was selected for analysis. The crystal structure of SARS-CoV-2 S protein RBD bound with hACE2 receptor was retrieved from Protein DataBank (http://www.rcsb.org/) as a reference (PDB ID: 6M0J). The hACE2 structure from the complex was extracted using BIOVIA discovery studio and assigned as chain A.

### 2.2. Retrieval of gene sequences

Full genome sequences of 26 variants of SARS-CoV-2 described by WHO were retrieved from GISAID (https://www.gisaid.org/) (**Supplementary Table S1**) on 4^th^ December 2021. For the prediction of coding sequence of RDB of S protein from all variants, Multiple Sequence Alignment (MSA) were performed by using MAFFT webtool (version 7) (Katoh and Standley, 2013). For MSA RBD of Wuhan variant (PDB ID: 6M0J) was used as a reference and RDB sequence of S protein of other 25 variants were retrieved by MAFFT webtool (version 7). The S protein sequences of all 25 variants were saved in FASTA format for further *in silico* studies.

### 2.3. In silico homology modelling

Due to the absence of the three-dimensional (3D) structure of RDB of S protein of all variants of SARS-CoV-2 in protein data bank (PDB), their structural models were generated using online homology modelling server, SWISS-MODEL (https://swissmodel.expasy.org/) (Waterhouse et al., 2018).

### 2.4. Model validations

The quality of each 3D model of SARS-CoV-2 RBD generated in SWISS-MODEL server were validated by two different web servers. Structure Analysis and Verification Server (SAVES) version 5.0 (https://servicesn.mbi.ucla.edu/SAVES/) uses Ramachandran plot, ERRAT score, verify 3D score to evaluate the models. Protein Structure Analysis (ProSA)-web server (https://prosa.services.came.sbg.ac.at/prosa.php) uses Z-score (indicates the overall model quality) to evaluate the models.

### 2.5. Protein-protein Docking

Docking between hACE2 and RBD of S protein of different variants of SARS-CoV-2 was performed using ClusPro server (https://cluspro.org) (Kozakov et al., 2017). Crystal PDB structure of hACE2 was uploaded in the ClusPro server as a receptor and validated PDB structure of RBD of S protein of all variants were uploaded one by one as a ligand. Using default setting of ClusPro server docking was done between A chain (of hACE2) and B chain (of S proteins). After each docking between hACE2 and S protein of each variant, thirty cluster models had been created, of which the model with lowest energy score were selected. The selected cluster models were downloaded in PDB format for further analysis.

### 2.6. Analysis of Direct Contact Residues of hACE2 : S Protein RBD

Direct contact residues between hACE2 : RBD of S protein of 25 all variants were analysed using PDBSum (https://www.ebi.ac.uk/thorntonsrv/databases/pdbsum/Generate.html) (Laskowski, 2001). After selecting the best cluster model of docking between hACE2 and RBD of S protein of each variant, the cluster model was uploaded in the PDBSum to analyse the interactions between them and analysed further.

### 2.7. Phylogenetic tree constructions

MGEAX was used to align the sequences of spike proteins from different variations. The phylogenetic analysis was carried out using the MGEAX neighbour-joining approach, which involved numerous comparisons. ClustalW multiple sequence alignment was used for multiple comparisons, neighbour-joining phylogenies were estimated, and 1000 bootstraps were used (Liu et al., 2020; Laskar and Ali, 2021a). Three different phylogenetic trees were made using sequence of RBD of all variants of SARS-CoV-2, S protein of all variants of SARS-CoV-2, whole genome sequence of all variants of SARS-CoV-2.

## 3. Results and Discussion

### 3.1. Model validation

To understand the interaction between the RBD of S protein of SARS-CoV-2 variants and hACE2 receptor, the 3D structures of the variant RBD were obtained by *in silico* modelling. For this purpose, the retrieved RBD sequences were used to model the 3D structures in SWISS Model and validated using SAVES ver 5.0 and ProSA web server. The sequence coverage, resolution, identity, and similarity with the query sequence obtained in SWISS model are tabulated in **Supplementary Table S2**. The Qualitative Model Energy Analysis (QMEAN) and global quality estimation score (GMQE) values of each model were checked to evaluate the biologically relevant models. The QMEAN value, also known as the “degree of nativeness,” reflects how well the model matches the experimental structures. A decent model should have a QMEAN value of not less than −4.0 and close to zero (Benkert et al., 2011). The QMEAN scores obtained for the predicted protein model fall within the recommended range validating the high quality of the modelled structures (**Supplementary Table S2**). Moreover, the GMQE results further validate the structures as the obtained values fall withing the standard range of 0 to 1 (Biasini et al., 2014).

Further, the RBD models of all the variants showed acceptable parameters as validated in SAVES ver 5.0 and ProSA web server (**Supplementary Table S3**). The models of all variants contain greater than 78% of its residues in the allowed region, according to the Ramachandran plot (Supplementary Figure S1). This result further verified the protein model’s high quality. These 3D structures of Spike RBD were considered for the interaction analysis with hACE2 receptor.

### 3.2. Omicron RBD shows strong affinity for hACE2

The hACE2:RBD complex retrieved from PDB (ID: 6M0J) was analysed using PDBSum web server. For the prediction of molecular interactions between the hACE2 receptor and RBD of S protein of SARS-CoV-2, protein-protein docking was performed using ClusPro web server. *In silico* molecular docking is a widely employed tool to predict the protein-protein interactions (Kozakov et al., 2017). The binding affinities of RBD across variants of SARS-CoV-2 to hACE2 were found to be different because of the sequence variations due to mutation, substitution or insertion in RBD. The binding efficiencies, represented in terms of lowest energy, between the hACE2 and variant RBDs are tabulated in **Table 1**. The lowest energy score of Omicron variant RBD with hACE2 was found to be the maximum among all the VOCs and VUMs (−1216.2). This indicates that the Omicron variant RBD binds to hACE2 with stronger affinity than other VOCs and VUMs, implying a larger transmission potential. However, following these dockings, an interesting observation was made when compared the docking score of the Omicron variant to the VOIs. The C.37 (Lambda) variant showed much greater lowest energy score (−1352.6) than the Omicron variant (Table 1). We may deduce from the docking score that the Lambda variant (C.37) should be closely monitored for its emergence as a lethal variant at a later stage.

**Table 1.** Lowest energy and interactions of RBD of SARS-CoV-2 variants with hACE2.

| SARS CoV-2 Variants | Lowest Energy (kcal/mol) | No of Bonds/Interactions |  |  |
| --- | --- | --- | --- | --- |
|  |  | H-bonds | Salt bridge | Non-bonded Contacts |
| WIV04 (Wuhan Variant) | - | 10 | 1 | 101 |
| B.1.1.7 GR/501Y.V1 (Alpha) | -1138.1 | 33 | 8 | 331 |
| B.1.351 GH/501Y.V2 (Beta) | -1186.3 | 27 | 9 | 399 |
| B.1.1.28.1, alias P.1 GR/501Y.V3 (Gamma) | -1183 | 19 | 8 | 227 |
| B.1.617 G/452R.V3 (Delta) | -1191 | 35 | 10 | 381 |
| B.1.1.529 (Omicron) | **-1216.2** | 28 | 6 | 258 |
| C.37, GR/452Q.V1 (Lambda) | -1352.6 | 26 | 6 | 238 |
| B.1.621, GH (Mu) | -861.1 | 16 | 4 | 232 |
| B.1.427<br>B.1.429 (Epsilon) | -1175 | 36 | 10 | 377 |
| R.1 | -823.8 | 14 | 5 | 238 |
| B.1.466.2 | -869.4 | 20 | 4 | 243 |
| B.1.1.318 | -823.8 | 14 | 5 | 238 |
| B.1.1.519 | -834.7 | 18 | 5 | 227 |
| C.36.3 | -1138.1 | 33 | 8 | 331 |
| B.1.214.2 | -897.9 | 20 | 5 | 206 |
| B.1.1.523 | -824.2 | 14 | 5 | 245 |
| B.1.619 | -823.8 | 14 | 5 | 238 |
| B.1.620 | -818.3 | 19 | 3 | 159 |
| C.1.2 | -1135.7 | 29 | 7 | 337 |
| B.1.617.1 (Kappa) | -1191 | 35 | 10 | 381 |
| B.1.526 (Iota) | -823.8 | 14 | 5 | 238 |
| B.1.525 (Eta) | -823.8 | 14 | 5 | 238 |
| B.1.630 | -1079.9 | 26 | 5 | 291 |
| B.1.640 | -1126.4 | 30 | 8 | 336 |
| AV.1 | -832.6 | 11 | 5 | 249 |
| AT.1 | -823.8 | 14 | 5 | 238 |

### 3.3. Omicron RBD shows unique interaction with hACE2

This study further envisaged to investigate the pattern of interactions of RBD variants with hACE2 as this interaction is a crucial point for the viral infectivity. To analyse the amino acid residues involved in direct interaction between RBD and hACE2, PDBSum (https://www.ebi.ac.uk/thornton-srv/databases/pdbsum/Generate.html) web server was used. The probable interacting residues of both the interacting partners were obtained in 2D format. All these interactions are presented in **Supplementary Figure S2**. From these results the interacting amino acids of RDB of all variants were extracted. Similarly, the interacting amino acids of hACE2 to the RDB were also extracted.

To understand the extent and diversity of these interactions, the interacting residues were closely analysed. The interacting amino acid residues of RBD of Omicron is compared with other 25 variants (including Wuhan Variant). The RBD of Omicron variant interacts using 30 amino acid residues with 41 residues on hACE2. These residues of Omicron RBD are completely different from those of Wuhan variant (**Figure 1A**). Similarly, the omicron variant uses unique binding residues than thirteen more variants. However, some variants show similarity binding residues with the Omicron. For instance, the Lamda variant (C.37) RBD interacts with the hACE2 with 100% identical residues with that of Omicron variant (Figure 1A). Other variants like C.1.2 and Beta (B.1.351) used 92.86% and 90.33% identical residues to that of Omicron RBD to interact with hACE2. Furthermore, we examined the interacting residues of hACE2 involved in binding the RDBs of different variations. The Beta variant (B.1.351) was found to interact with the hACE2 on 58.93% identical residues that are recognized by the Omicron RBD (**Figure 1B**). More than half (51.1%) of the amino acid residues of hACE2 to which the Omicron interacts are identical to that of the C.1.2 variant. The resemblance between the Omicron and the rest of the variants is less than 50% in the recognizing residues on hACE2. The interaction of hACE2 with Omicron variant RBD was compared with all other variant RBDs and the relative number of identical residues are presented in **Supplementary Table S4**. This is the first report encompassing comparative analysis of interacting residues of RBD of variants and hACE2. The present dataset demonstrates that the novel variant, Omicron, interacts with unique residues to new amino acid residues in the hACE2 receptor, raising the question of infectivity, vaccination potential and immune evasion. A recent study based on hACE: RBD interaction, suggested that the Omicron variant show over ten times higher infectivity than the Wuhan variant, are capable of escape currently available vaccines and efficacy of neutralizing monoclonal antibodies (Chen et al., 2021) supporting our prediction. The interaction of hACE with S protein RBD of Omicron variant and three variants with over ninety percent identical residues as Omicron namely Beta (B.1.351), Lamda (C.37) and C.1.2, are shown in Figure 2. The interaction of Wuhan variant (WIV04) was represented as reference. Moreover, the 3D illustrations of the corresponding interactions are shown in **Figure 3**.

**Figure 1.**
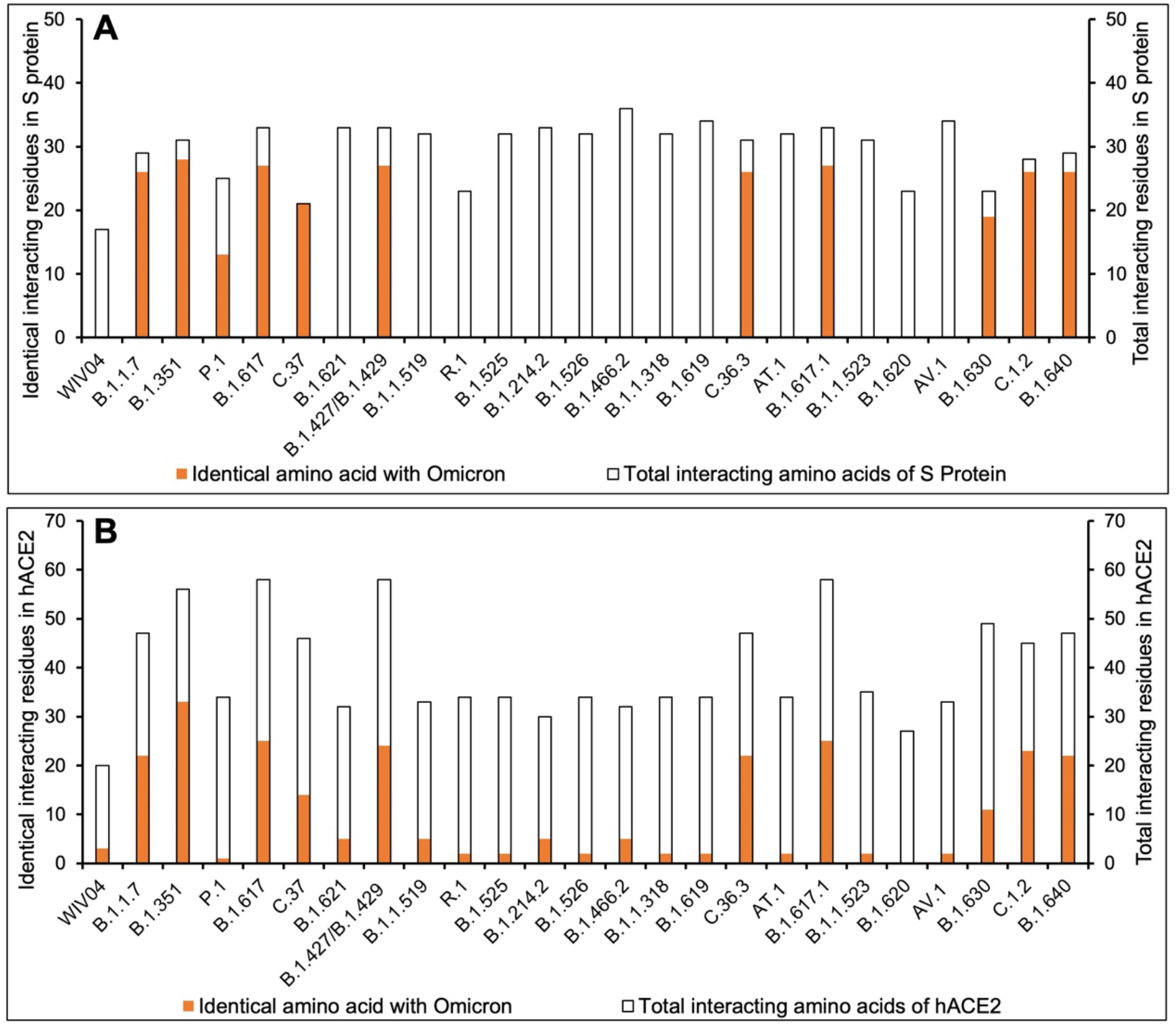
Comparative interacting residues of (A) RBD of SARS-CoV-2 variants and (B) hACE2 with respect of Omicron variant. The protein-protein interaction between hACE2 and SARS-CoV-2 RBD was studied by performing molecular using ClusPro server (https://cluspro.org) and analysed in PDBSum. The number of interacting residues were calculated and identical residues to that of Omicron variant were calculated.

**Figure 2.**
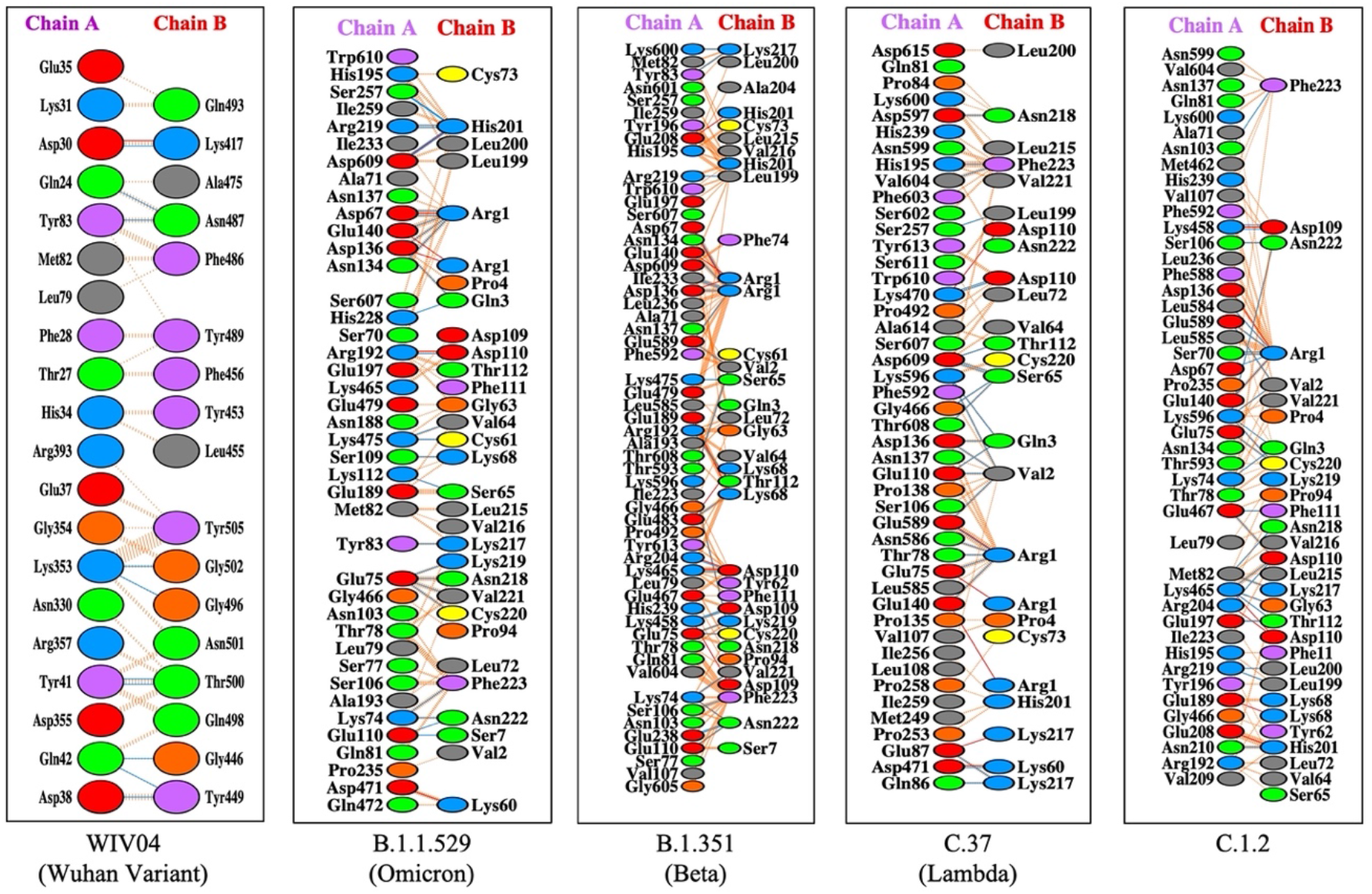
Protein–protein docking representation of hACE2 and RBD of SARS CoV-2. The interaction representation, which includes hydrogen (blue line), salt bridges (red line), and non-bonded (orange dash) interactions. Chain A represents hACE2 and chain B represents RBD. Residue colours: positive (blue): His, Lys, Arg; Negative (red): Asp, Glu; Neutral (green): Ser, Thr, Asn, Gln; Aliphatic (grey): Ala, Val, Leu, Ile, Met; Aromatic (purple): Phe, Tyr, Trp; and Pro & Gly (orange).

**Figure 3.**
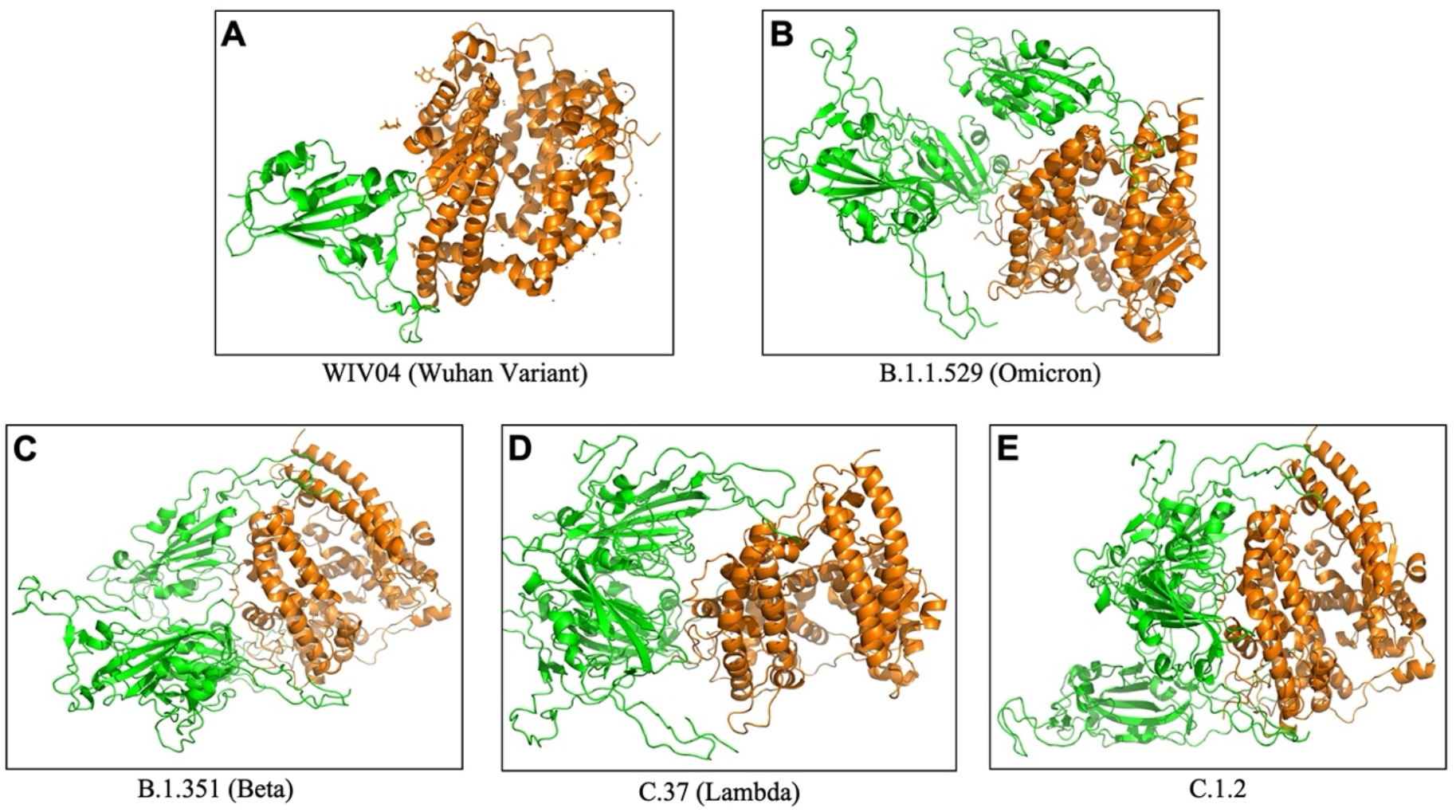
The 3D binding interface of S protein RBD (Green) and hACE2 (Orange) complex obtained from ClusPro.

The interactions of hACE2 with the RBD of the currently listed variants (26 including reference) by WHO (https://www.who.int/en/activities/tracking-SARS-CoV-2-variants/) were covered in this study and compared with the Omicron variant. However, SARS-CoV-2 is continuously evolving due to the mutations in the genome and there are hundreds of lineages available which are not listed as WHO variants (Rambaut et al., 2020; Laskar and Ali, 2021a, 2021b). The present study along with the available records suggests a probability for the incidence of mutations in the lineages with higher infectivity that will eventually emerge as a novel variant in near future.

### 3.3. Omicron RBD evolution may be independent of other variants

Viruses are continually evolving due to mutations in their genetic coding as they spread and in the process creating a roadmap for evolving and varying pathological manifestations (Laskar and Ali, 2021c). To understand the evolution pattern of RBD of Omicron variant, the phylogenetic tree was prepared along with other variants using MEGA X software (**Figure 4**). The phylogenetic analysis was performed at three different levels leading to some interesting observations when we looked at the positioning of Omicron variant across the trees. First, the phylogenetic tree of complete genome (**Figure 4A**) when compared with the one based on S protein only (**Figure 4B**) reveals that albeit the positioning of variants is different, some sense of similarity persists. This is clearly representative of the mutations incident in genomic regions outside of the S protein. Secondly, the whole genome phylogenetic tree revealed the Lamda (C.37) variant to be in close proximity to Omicron both of which had nearly identical residues in the presently studied interactions. Lastly, the phylogenetic tree prepared on the basis of RBD sequence of different SARS-CoV-2 variants showed that the Omicron variant has independently emerged of the rest of the variants (**Figure 4C**). However, it is most closely related to Beta variant (B.1.351), followed by Gamma variant (B.1.1.28.1), but distantly related to the Wuhan variant. This is reflective of the accrued mutations in the RBD with time. The Omicron variant is similarly distantly localized in the phylogenetic tree based on S protein sequence but has different companions (**Figure 4B**). The above observations signify that even though evolution is being driven by mutations across the genome, the ones affecting the RBD-hACE2 interaction have a greater probability of emerging as new variant.

**Figure 4.**
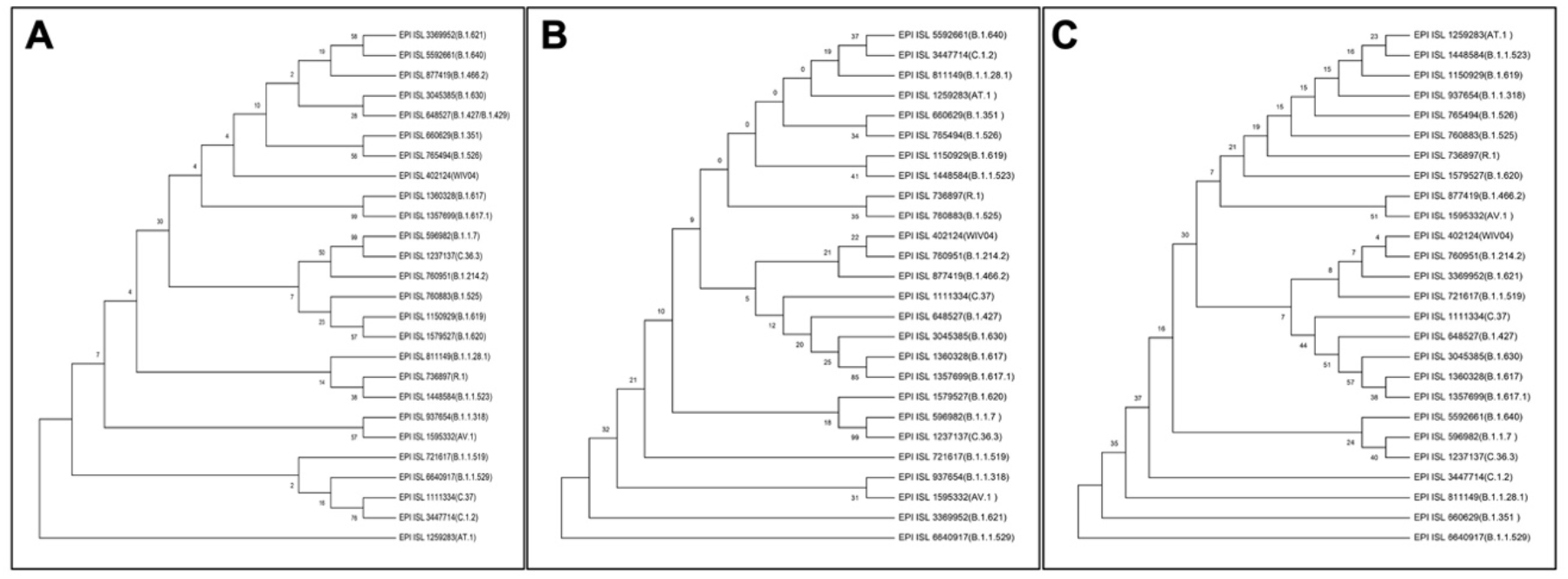
Phylogenetic Tree of SARS-CoV-2 variants constructed using (A) Whole Genome (B) S protein and (C) RBD.

## 4. Conclusion

The Omicron variant interacts with the hACE2 receptor with strong affinity involving unique amino acid residues than most of the SARS-CoV-2 variants suggesting increased infectivity and immune/vaccine evasion potential of the variant. However, the Lamda variant (C.37) interacts with hACE2 with higher affinity using identical RBD residues of the Omicron. This study calls for a closer observation on the evolution of SARS-CoV-2 with the genomic alterations affecting the RBD:hACE2 interaction as the key basis for monitoring.

## Acknowledgement

The authors are thankful to the Department of Biological Sciences, Aliah University, Kolkata, India for providing the necessary facilities for this work. SSMA acknowledges the Council of Scientific & Industrial Research (CSIR) Govt. of India for financial assistance in the form of Junior Research Fellowship (JRF).

## Conflict of interests

The authors declare that there exists no conflict of interests.

## Notes

### Competing Interest Statement

The authors have declared no competing interest.

