## Supplementary for "Analyzing the interaction of human ACE2 and RBD of spike protein of SARS-CoV-2 in perspective of Omicron variant"

### ***SUPPLEMENTARY FILES***

### Interactions of Human ACE2 with Spike Protein RBD of SARS-Cov-2 Variants: A Comparative Analysis with the Omicron Variant


**Arijit Samanta[a], Syed Sahajada Mahafujul Alam[a], Safdar Ali[b], Mehboob Hoque[a]***

*[a]Applied BioChemistry Laboratory, Department of biological Sciences, Aliah University, Kolkata 700160, INDIA*

*[b]Clinical and Applied Genomics (CAG) Laboratory, Department of biological Sciences, Aliah University, Kolkata 700160, INDIA*

*Correspondence

Dr Mehboob Hoque

Department of Biological Sciences, Aliah University, Kolkata 700160, India

#### *SUPPLEMENTARY TABLES*

**Supplementary Table S1. Details of the SARS-CoV-2 variants.** The first submitted complete genome sequences were retrieved from GISAID.

| Sl No | Variants | Virus name | Accession ID | Submission Date | First detected in | Genome size (nucleotide) |
|---|---|---|---|---|---|---|
| **Reference** | | | | | | |
| 1 | (**Wuhan Variant)** | WIV04 | EPI_ISL_402124 | 11.01.2020 | Wuhan | 29891 |
| **Variants of Concerns (VOCs)** | | | | | | |
| 2 | B.1.1.7 GR/501Y.V1 (Alpha) | hCoV-19/England/CAMC-A58BA4/2020 | EPI_ISL_596982 | 2020-10-27 | United Kingdom | 29763 |
| 3 | B.1.351 GH/501Y.V2 (Beta) | hCoV-19/South Africa/KRISP-K004599/2020 | EPI_ISL_660629 | 2020-11-27 | South Africa | 29848 |
| 4 | B.1.1.28.1, alias P.1 GR/501Y.V3 (Gamma) | hCoV-19/Brazil/AM-FIOCRUZ-20143138FN-R2/2020 | EPI_ISL_811149 | 2021-01-13 | Brazil | 29593 |
| 5 | B.1.617 G/452R.V3 (Delta) | hCoV-19/India/MH-NEERI-NGP-29787/2021 | EPI_ISL_1360328 | 2021-03-25 | India | 29839 |
| 6 | B.1.1.529 (Omicron) | hCoV19/Botswana/R40B60_BHP_ 3321001247/2021 | EPI_ISL_6640917 | 2021-11-23 | Botswana/Hong Kong/South Africa | 29684 |
| **Variants of Interest (VOIs)** | | | | | | |
| 7 | C.37, GR/452Q.V1 (Lambda) | hCoV-19/Peru/LIM-INS-436/2021 | EPI_ISL_1111334 | 2021-03-01 | Peru | 29901 |
| 8 | B.1.621, GH (Mu) | hCoV-19/Colombia/RIS-VG-3857/2021 | EPI_ISL_3369952 | 2021-08-12 | Colombia | 29902 |
| **Variants Under Monitoring (VUMs)** | | | | | | |
| 9 | C.1.2 | hCoV-19/South Africa/KRISP-K020275/2021 | EPI_ISL_3447714 | 2021-08-17 | South Africa | 29803 |
| 10 | B.1.617.1 (Kappa) | hCoV-19/India/WB-1931300244219/2021 | EPI_ISL_1357699 | 2021-03-25 | India | 29781 |
| 11 | B.1.630 | hCoV-19/Dominican Republic/33884/2021 | EPI_ISL_3045385 | 2021-07-22 | Dominican Republic | 29799 |
| 12 | B.1.640 | hCoV-19/Congo/FCRM-100-A32.28.09.21/2021 | EPI_ISL_5592661 | 2021-10-27 | Republic of Congo | 29683 |
| **Formerly Under Monitoring** | | | | | | |
| 13 | B.1.427 B.1.429 (Epsilon) | hCoV-19/USA/CA-CZB-12872/2020 | EPI_ISL_648527 | 2020-11-21 | United States of America | 29816 |
| 14 | R.1 | hCoV-19/Japan/IC-0386/2020 | EPI_ISL_736897 | 2020-12-26 | Asia / Japan | 29770 |
| 15 | B.1.466.2 | hCoV-19/Indonesia/JI-ITDua-33281NT/2020 | EPI_ISL_877419 | 2021-01-27 | Indonesia | 29890 |
| 16 | B.1.1.318 | hCoV-19/England/ALDP-115DB1E/2021 | EPI_ISL_937654 | 2021-02-05 | Europe / United Kingdom / England | 29753 |
| 17 | B.1.1.519 | hCoV-19/Japan/IC-0377/2020 | EPI_ISL_721617 | 2020-12-21 | Asia / Japan | 29768 |
| 18 | C.36.3 | hCoV-19/USA/TX-HMH-MCoV-27355/2021 | EPI_ISL_1237137 | 2021-03-13 | North America / USA / Texas / Houston | 29804 |

| | | | | | | |
|---|---|---|---|---|---|---|
| 19 | B.1.214.2 | hCoV-19/England/ALDP-CB0759/2020 | EPI_ISL_760951 | 2021-01-04 | Europe / United Kingdom / England | 29750 |
| 20 | B.1.1.523 | hCoV-19/Switzerland/GE-33615516/2021 | EPI_ISL_1448584 | 2021-04-02 | Europe / Switzerland / Geneva | 29856 |
| 21 | B.1.619 | hCoV-19/Germany/BE-RKI-I-020766/2021 | EPI_ISL_1150929 | 2021-03-04 | Europe / Germany / Berlin | 29809 |
| 22 | B.1.620 | hCoV-19/Lithuania/LTU000_NMVRI55624/2021 | EPI_ISL_1579527 | 2021-04-13 | Europe / Lithuania / Vilniaus apskritis | 29840 |
| 23 | B.1.526 (Lota) | hCoV-19/USA/NY-Wadsworth-290357-01/2020 | EPI_ISL_765494 | 2021-01-04 | United States of America | 29782 |
| 24 | B.1.525 (Eta) | hCoV-19/England/CAMC-C769B3/2020 | EPI_ISL_760883 | 2021-01-04 | United Kingdom | 29709 |
| 25 | AV.1 | hCoV-19/England/QEUH-148F257/2021 | EPI_ISL_1595332 | 2021-04-15 | United Kingdom | 29764 |
| 26 | AT.1 | hCoV-19/Russia/PSK-16/2021 | EPI_ISL_1259283 | 2021-03-16 | Russian Federation | 29803 |

**Supplementary Table S2. SWISS Model parameters of the predicted 3D structures of RBDs of SARS-CoV-2 variants.**

| Variants | Accession ID | Template Swiss model ID | Features of template used in generating S Protein model | | | | | Quality assessment of the S Protein model | |
|---|---|---|---|---|---|---|---|---|---|
| | | | Sequence Identity | Method | Resolution | Sequence similarity | Coverage | QMEAN | GMQE |
| B.1.1.7 GR/501Y.V1 (Alpha) | EPI_ISL_596982 | 7lwv.1.A | 100.00% | EM | - | 0.63 | 1.00 | 0.78 | 0.84 |
| B.1.351 GH/501Y.V2 (Beta) | EPI_ISL_660629 | 7lww.1.A | 100.00% | EM | - | 0.63 | 1.00 | 0.77 | 0.84 |
| B.1.1.28.1, alias P.1 GR/501Y.V3 (Gamma) | EPI_ISL_811149 | 7v78.1.A | 100.00% | EM | - | 0.63 | 1.00 | 0.74 | 0.81 |
| B.1.617 G/452R.V3 (Delta) | EPI_ISL_1360328 | 7cn4.1.A | 89.69% | EM | - | 0.59 | 1.00 | 0.70 | 0.77 |
| B.1.1.529 (Omicron) | EPI_ISL_6640917 | 7lyq.1.A | 94.17% | EM | - | 0.61 | 1.00 | 0.74 | 0.81 |
| C.37, GR/452Q.V1 (Lambda) | EPI_ISL_1111334 | 6zgf.1.A | 89.69% | EM | - | 0.59 | 1.00 | 0.72 | 0.78 |
| B.1.621, GH (Mu) | EPI_ISL_3369952 | 6xc4.1.A | 99.55% | X-ray | 2.34 Å | 0.63 | 1.00 | 0.80 | 0.80 |
| B.1.427 B.1.429 (Epsilon) | EPI_ISL_648527 | 7cn4.1.A | 89.69% | EM | - | 0.59 | 1.00 | 0.70 | 0.77 |
| R.1 | EPI_ISL_736897 | 6xc4.1.A | 99.55% | X-ray | 2.34 Å | 0.63 | 1.00 | 0.80 | 0.80 |
| B.1.466.2 | EPI_ISL_877419 | 6xc4.1.A | 99.55% | X-ray | 2.34 Å | 0.63 | 1.00 | 0.80 | 0.80 |
| B.1.1.318 | EPI_ISL_937654 | 6xc4.1.A | 99.55% | X-ray | 2.34 Å | 0.63 | 1.00 | 0.80 | 0.80 |
| B.1.1.519 | EPI_ISL_721617 | 6xc4.1.A | 99.55% | X-ray | 2.34 Å | 0.63 | 1.00 | 0.80 | 0.80 |
| C.36.3 | EPI_ISL_1237137 | 7lwv.1.A | 100.00% | EM | - | 0.63 | 1.00 | 0.78 | 0.84 |
| B.1.214.2 | EPI_ISL_760951 | 6xc4.1.A | 99.55% | X-ray | 2.34 Å | 0.63 | 1.00 | 0.80 | 0.80 |
| B.1.1.523 | EPI_ISL_1448584 | 6xc4.1.A | 99.10% | X-ray | 2.34 Å | 0.63 | 1.00 | 0.80 | 0.80 |
| B.1.619 | EPI_ISL_1150929 | 6xc4.1.A | 99.55% | X-ray | 2.34 Å | 0.63 | 1.00 | 0.80 | 0.80 |
| B.1.620 | EPI_ISL_1579527 | 6xc4.1.A | 99.10% | X-ray | 2.34 Å | 0.63 | 1.00 | 0.80 | 0.80 |
| C.1.2 | EPI_ISL_3447714 | 7lwv.1.A | 99.10% | EM | - | 0.63 | 1.00 | 0.77 | 0.84 |
| B.1.617.1 (Kappa) | EPI_ISL_1357699 | 7cn4.1.A | 89.69% | EM | - | 0.59 | 1.00 | 0.70 | 0.77 |
| B.1.526 (Lota) | EPI_ISL_765494 | 6xc4.1.A | 99.55% | X-ray | 2.34 Å | 0.63 | 1.00 | 0.80 | 0.80 |
| B.1.525 (Eta) | EPI_ISL_760883 | 6xc4.1.A | 99.55% | X-ray | 2.34 Å | 0.63 | 1.00 | 0.80 | 0.80 |
| B.1.630 | EPI_ISL_3045385 | 7sbl.1.A | 99.10% | EM | 3.44 Å | 0.63 | 1.00 | 0.73 | 0.10 |
| B.1.640 | EPI_ISL_5592661 | 7lwv.1.A | 98.21% | EM | - | 0.63 | 1.00 | 0.78 | 0.84 |
| AV.1 | EPI_ISL_1595332 | 6xc4.1.A | 99.10% | X-ray | 2.34 Å | 0.63 | 1.00 | 0.81 | 0.80 |
| AT.1 | EPI_ISL_1259283 | 6xc4.1.A | 99.55% | X-ray | 2.34 Å | 0.63 | 1.00 | 0.80 | 0.80 |

**Supplementary Table S3. S protein RBD model validation parameters obtained in SAVES and ProSA.**

| Variants | Accession ID | ERRAT | % of the residues have averaged 3D-1D score >= 0.2 | Ramachandran plot | | | G-factors | *Z*-Score |
|---|---|---|---|---|---|---|---|---|
| | | | | core | allow | gener | | |
| B.1.1.7 GR/501Y.V1 (Alpha) | EPI_ISL_596982 | **94.5364** | 93.12% | 88.4% | 11.2% | 0.3% | -0.19 | **-5.93** |
| B.1.351 GH/501Y.V2 (Beta) | EPI_ISL_660629 | **94.0594** | 91.18% | 89.3% | 10.5% | 0.2% | -0.20 | **-5.62** |
| B.1.1.28.1, alias P.1 GR/501Y.V3 (Gamma) | EPI_ISL_811149 | **81.1287** | 87.89% | 87.0% | 12.4% | 0.3% | -0.14 | **-5.48** |
| B.1.617 G/452R.V3 (Delta) | EPI_ISL_1360328 | **93.6594** | 93.72% | 78.6% | 21.2% | 0.2% | -0.21 | **-5.58** |
| B.1.1.529 (Omicron) | EPI_ISL_6640917 | **96.6443** | 95.37% | 87.7% | 11.8% | 0.5% | -0.24 | **-5.56** |
| C.37, GR/452Q.V1 (Lambda) | EPI_ISL_1111334 | **80** | 96.71% | 83.2% | 16.2% | 0.5% | -0.22 | **-5.52** |
| B.1.621, GH (Mu) | EPI_ISL_3369952 | **97.076** | 100.00% | 90.6% | 9.4% | 0.0 | -0.14 | **-5.96** |
| B.1.427 B.1.429 (Epsilon) | EPI_ISL_648527 | **93.8739** | 93.57% | 78.4% | 20.9% | 0.7% | -0.22 | **-5.6** |
| R.1 | EPI_ISL_736897 | **96.5116** | 97.99% | 90.6% | 9.4% | 0.0% | -0.13 | **-5.88** |
| B.1.466.2 | EPI_ISL_877419 | **96.5116** | 96.98% | 90.6% | 9.4% | 0.0% | -0.14 | **-5.87** |
| B.1.1.318 | EPI_ISL_937654 | **96.5116** | 97.99% | 90.6% | 9.4% | 0.0% | -0.13 | **-5.88** |
| B.1.1.519 | EPI_ISL_721617 | **97.0588** | 97.99% | 90.6% | 9.4% | 0.0% | -0.15 | **-6.01** |
| C.36.3 | EPI_ISL_1237137 | **94.5364** | 93.12% | 88.4% | 11.2% | 0.3% | -0.19 | **-5.93** |
| B.1.214.2 | EPI_ISL_760951 | **96.5116** | 97.99% | 90.6% | 9.4% | 0.0% | -0.14 | **-5.99** |
| B.1.1.523 | EPI_ISL_1448584 | **96.5116** | 98.49% | 90.6% | 9.4% | 0.0% | -0.14 | **-5.84** |
| B.1.619 | EPI_ISL_1150929 | **96.5116** | 97.99% | 90.6% | 9.4% | 0.0% | -0.13 | **-5.88** |
| B.1.620 | EPI_ISL_1579527 | **97.076** | 97.99% | 90.6% | 9.4% | 0.0% | -0.13 | **-5.92** |
| C.1.2 | EPI_ISL_3447714 | **94.4816** | 93.12% | 88.4% | 11.2% | 0.3% | -0.18 | **-6** |
| B.1.617.1 (Kappa) | EPI_ISL_1357699 | **93.6594** | 93.72% | 78.6% | 21.2% | 0.2% | -0.21 | **-5.58** |
| B.1.526 (Lota) | EPI_ISL_765494 | **96.5116** | 97.99% | 90.6% | 9.4% | 0.0% | -0.13 | **-5.88** |
| B.1.525 (Eta) | EPI_ISL_760883 | **96.5116** | 97.99% | 90.6% | 9.4% | 0.0% | -0.13 | **-5.88** |
| B.1.630 | EPI_ISL_3045385 | **89.2857** | 93.27% | 86.5% | 13.1% | 0.3% | -0.25 | **-5.28** |
| B.1.640 | EPI_ISL_5592661 | **94.01** | 92.97% | 88.3% | 11.4% | 0.3% | -0.19 | **-6.08** |
| AV.1 | EPI_ISL_1595332 | **96.5116** | 96.98% | 90.6% | 9.4% | 0.0% | -0.13 | **-5.87** |
| AT.1 | EPI_ISL_1259283 | **96.5116** | 97.99% | 90.6% | 9.4% | 0.0% | -0.13 | **-5.88** |

**Supplementary Table S4. Comparative analysis of interacting amino acid residues between hACE and Omicron RBD with all other variant RBD**

| Variants | S Protein RBD | | | hACE2 Receptor | | |
|---|---|---|---|---|---|---|
| | No. of Total Interacting Amino Acid(s) | No. of Amino Acid residues identical to Omicron | % of identical residues with Omicron | No. of Total Interacting Amino Acid(s) | No. of Amino Acid residues identical to Omicron | % of identical residues with Omicron |
| EPI ISL 402124(WIV04) | 17 | 0 | **0** | 20 | 3 | 15 |
| EPI ISL 596982(B.1.1.7 ) | 29 | 26 | 89.66 | 47 | 22 | 46.81 |
| EPI ISL 660629(B.1.351 ) | 32 | 28 | **90.32** | 56 | 33 | 58.93 |
| EPI ISL 811149(B.1.1.28.1) | 25 | 13 | 52 | 34 | 1 | 2.94 |
| EPI ISL 1360328(B.1.617) | 33 | 27 | 81.82 | 58 | 25 | 43.10 |
| EPI ISL 1111334(C.37) | 21 | 21 | **100** | 46 | 14 | 30.43 |
| EPI ISL 3369952(B.1.621) | 33 | 0 | **0** | 32 | 5 | 15.63 |
| EPI ISL 648527(B.1.427) | 33 | 27 | 81.82 | 58 | 24 | 41.38 |
| EPI ISL 736897(R.1) | 23 | 0 | **0** | 34 | 2 | 5.88 |
| EPI ISL 877419(B.1.466.2) | 36 | 0 | **0** | 32 | 5 | 15.63 |
| EPI ISL 937654(B.1.1.318) | 32 | 0 | **0** | 34 | 2 | 5.88 |
| EPI ISL 721617(B.1.1.519) | 32 | 0 | **0** | 33 | 5 | 15.15 |
| EPI ISL 1237137(C.36.3) | 31 | 26 | 83.87 | 47 | 22 | 46.81 |
| EPI ISL 760951(B.1.214.2) | 33 | 0 | **0** | 30 | 5 | 16.67 |
| EPI ISL 1448584 (B.1.1.523) | 31 | 0 | **0** | 35 | 2 | 5.71 |
| EPI ISL 1150929(B.1.619) | 34 | 0 | **0** | 34 | 2 | 5.88 |
| EPI ISL 1579527(B.1.620) | 23 | 0 | **0** | 27 | 0 | **0** |
| EPI ISL 3447714(C.1.2) | 28 | 26 | **92.86** | 45 | 23 | 51.11 |
| EPI ISL 1357699(B.1.617.1) | 33 | 27 | 81.82 | 58 | 25 | 43.10 |
| EPI ISL 765494(B.1.526) | 32 | 0 | **0** | 34 | 2 | 5.88 |
| EPI ISL 760883(B.1.525) | 32 | 0 | **0** | 34 | 2 | 5.88 |
| EPI ISL 3045385(B.1.630) | 23 | 19 | 82.61 | 49 | 11 | 22.45 |
| EPI ISL 5592661(B.1.640) | 29 | 26 | 89.66 | 47 | 22 | 46.81 |
| EPI ISL 1595332(AV.1) | 34 | 0 | **0** | 33 | 2 | 6.06 |
| EPI ISL 1259283(AT.1) | 32 | 0 | **0** | 34 | 2 | 5.89 |

#### ***SUPPLEMENTARY FIGURES***

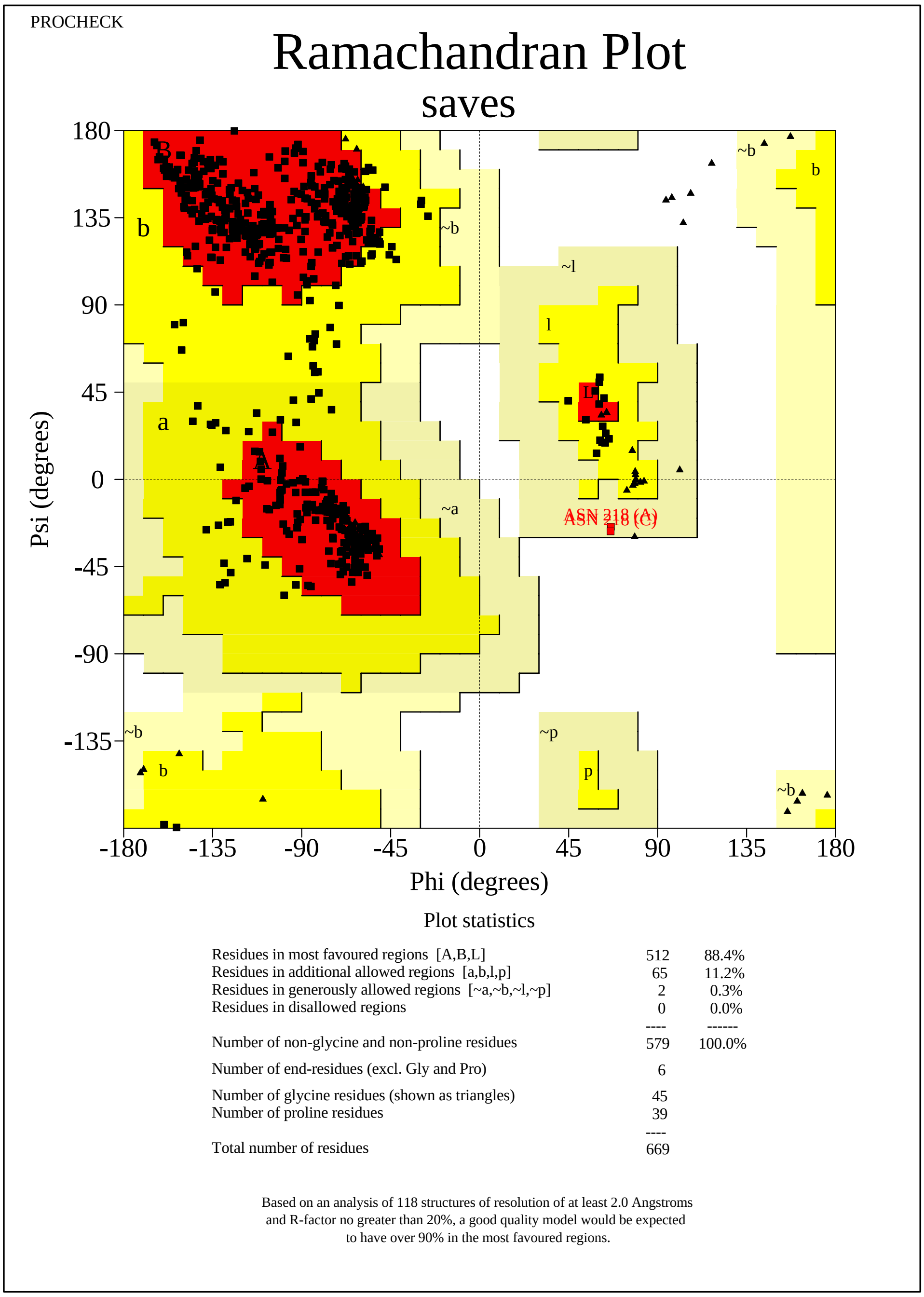

saves_01.ps

Supplementary Figure S1 (A). The PROCHECK Ramachandran plot for the validation of the 3D model of Spike RBD of SARS-CoV-2 B.1.1.7(Alpha) variant (EPI_ISL_596982).

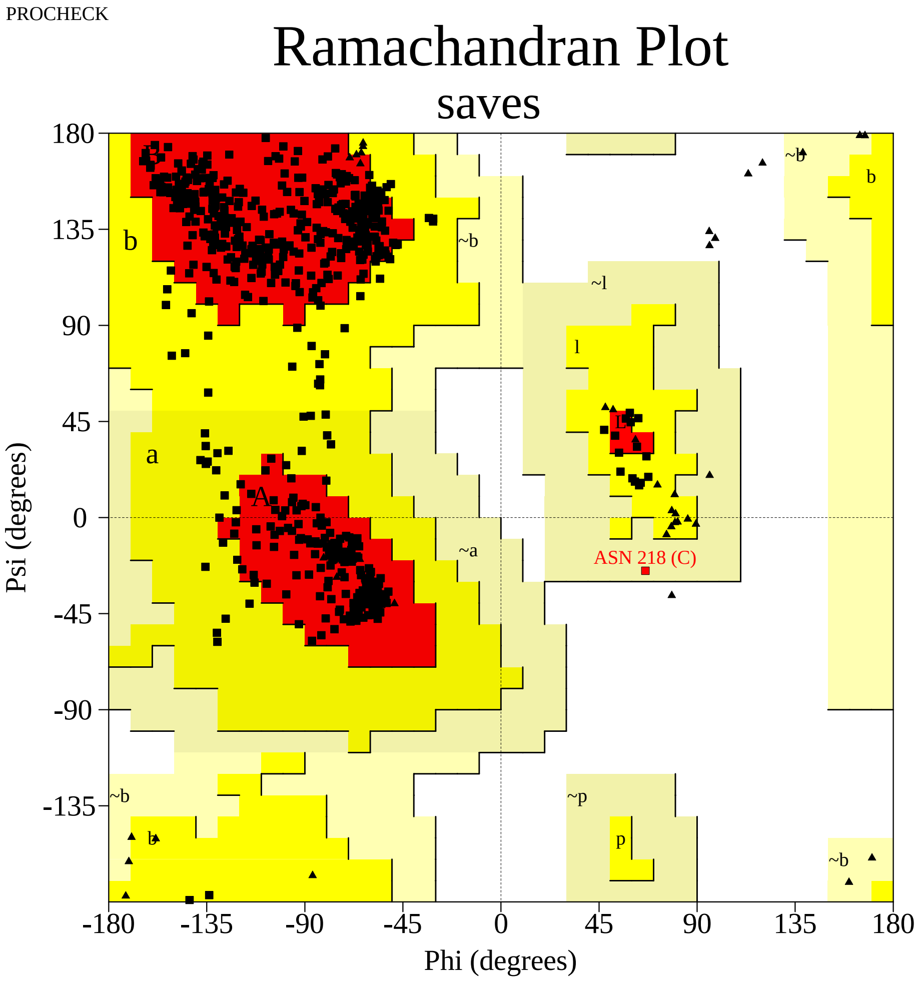


Plot statistics

| | | |
|---|---|---|
| Residues in most favoured regions [A,B,L] | 517 | 89.3% |
| Residues in additional allowed regions [a,b,l,p] | 61 | 10.5% |
| Residues in generously allowed regions [~a,~b,~l,~p] | 1 | 0.2% |
| Residues in disallowed regions | 0 | 0.0% |
|  | ---- | ------ |
| Number of non-glycine and non-proline residues | 579 | 100.0% |
| Number of end-residues (excl. Gly and Pro) | 6 | |
| Number of glycine residues (shown as triangles) | 45 | |
| Number of proline residues | 39 | |
|  | ---- |  |
| Total number of residues | 669 | |

Based on an analysis of 118 structures of resolution of at least 2.0 Angstroms and R-factor no greater than 20%, a good quality model would be expected to have over 90% in the most favoured regions.

saves_01.ps

Supplementary Figure S1 (B). The PROCHECK Ramachandran plot for the validation of the 3D model of Spike RBD of SARS-CoV-2 B.1.351(Beta) variant (EPI_ISL_660629)

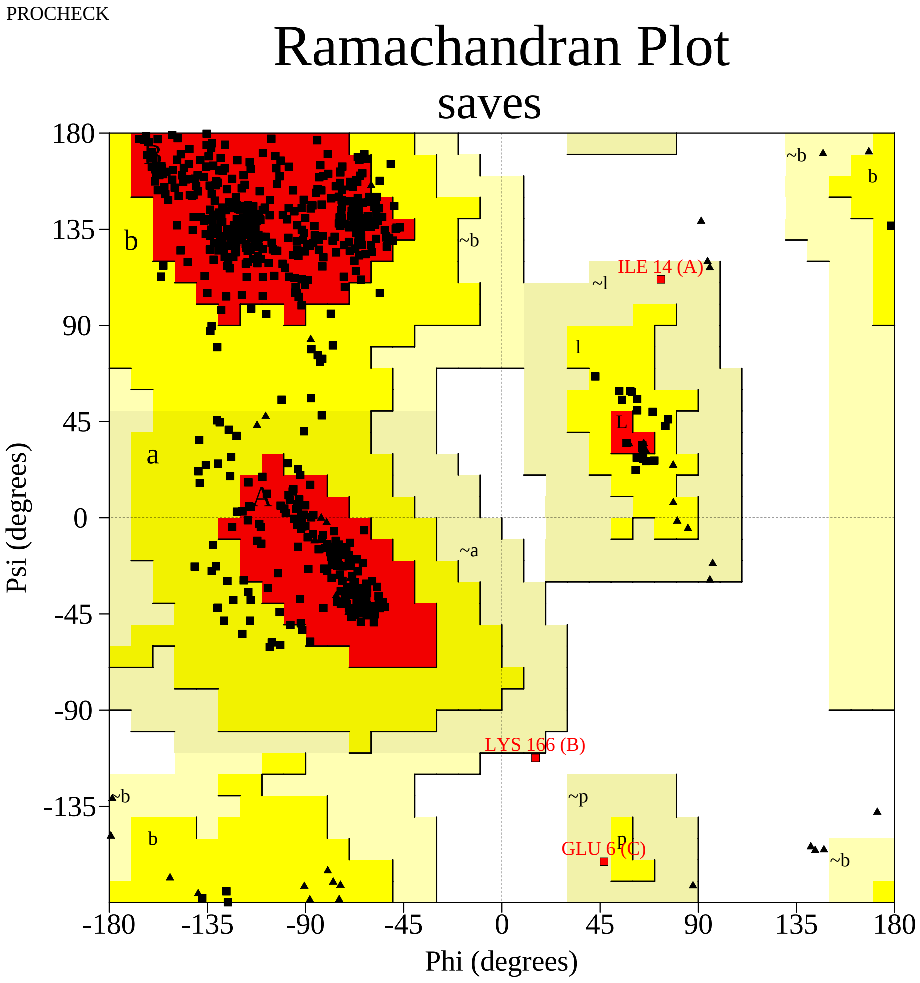


Plot statistics

| | | |
|---|---|---|
| Residues in most favoured regions [A,B,L] | 504 | 87.0% |
| Residues in additional allowed regions [a,b,l,p] | 72 | 12.4% |
| Residues in generously allowed regions [~a,~b,~l,~p] | 2 | 0.3% |
| Residues in disallowed regions | 1 | 0.2% |
|  | ---- | ------ |
| Number of non-glycine and non-proline residues | 579 | 100.0% |
| Number of end-residues (excl. Gly and Pro) | 6 | |
| Number of glycine residues (shown as triangles) | 45 | |
| Number of proline residues | 39 | |
|  | ---- |  |
| Total number of residues | 669 | |

Based on an analysis of 118 structures of resolution of at least 2.0 Angstroms and R-factor no greater than 20%, a good quality model would be expected to have over 90% in the most favoured regions.

saves_01.ps

Supplementary Figure S1 (C). The PROCHECK Ramachandran plot for the validation of the 3D model of Spike RBD of SARS-CoV-2 B.1.1.28.1(Gamma) variant EPI_ISL_811149

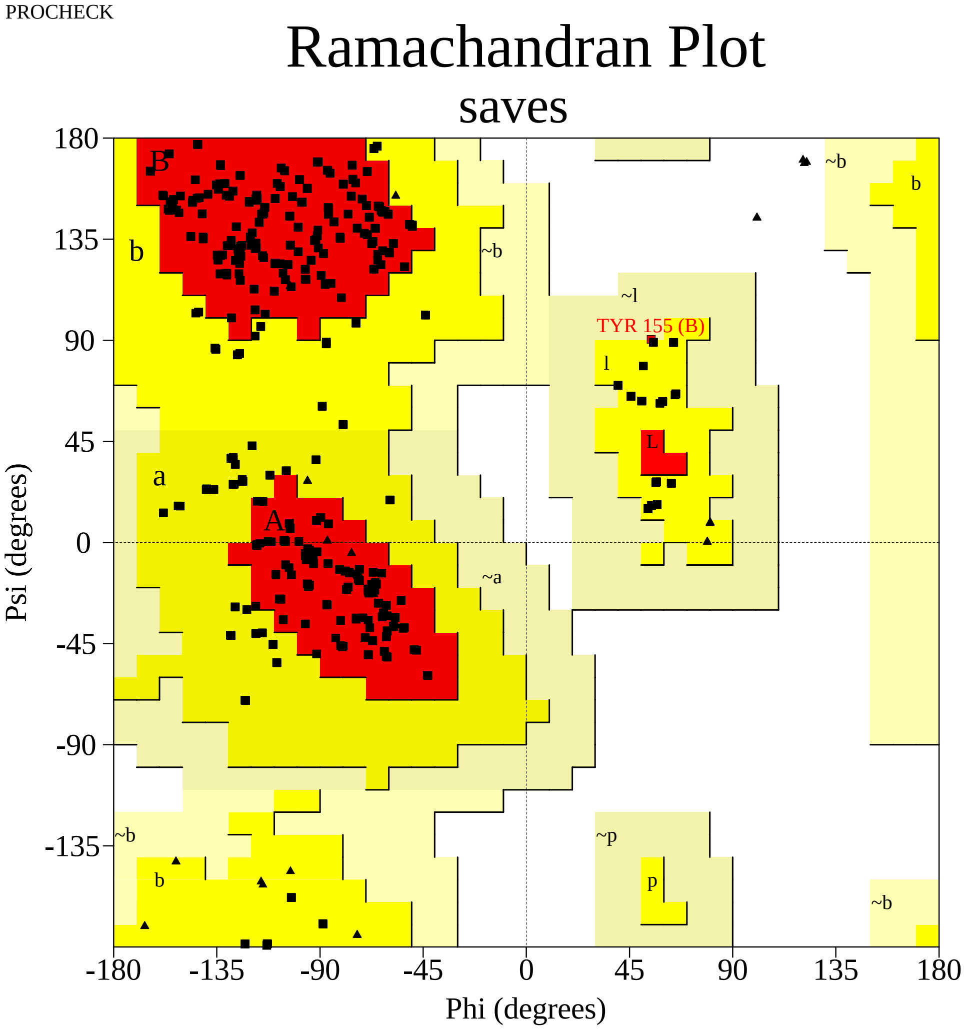


Plot statistics

| | | |
|---|---|---|
| Residues in most favoured regions [A,B,L] | 455 | 78.6% |
| Residues in additional allowed regions [a,b,l,p] | 123 | 21.2% |
| Residues in generously allowed regions [~a,~b,~l,~p] | 1 | 0.2% |
| Residues in disallowed regions | 0 | 0.0% |
|  | ---- | ------ |
| Number of non-glycine and non-proline residues | 579 | 100.0% |
| Number of end-residues (excl. Gly and Pro) | 6 | |
| Number of glycine residues (shown as triangles) | 45 | |
| Number of proline residues | 39 | |
|  | ---- |  |
| Total number of residues | 669 | |

Based on an analysis of 118 structures of resolution of at least 2.0 Angstroms and R-factor no greater than 20%, a good quality model would be expected to have over 90% in the most favoured regions.

saves_01.ps

Supplementary Figure S1 (D). The PROCHECK Ramachandran plot for the validation of the 3D model of Spike RBD of SARS-CoV-2 B.1.617(Delta) variant (EPI_ISL_1360328)

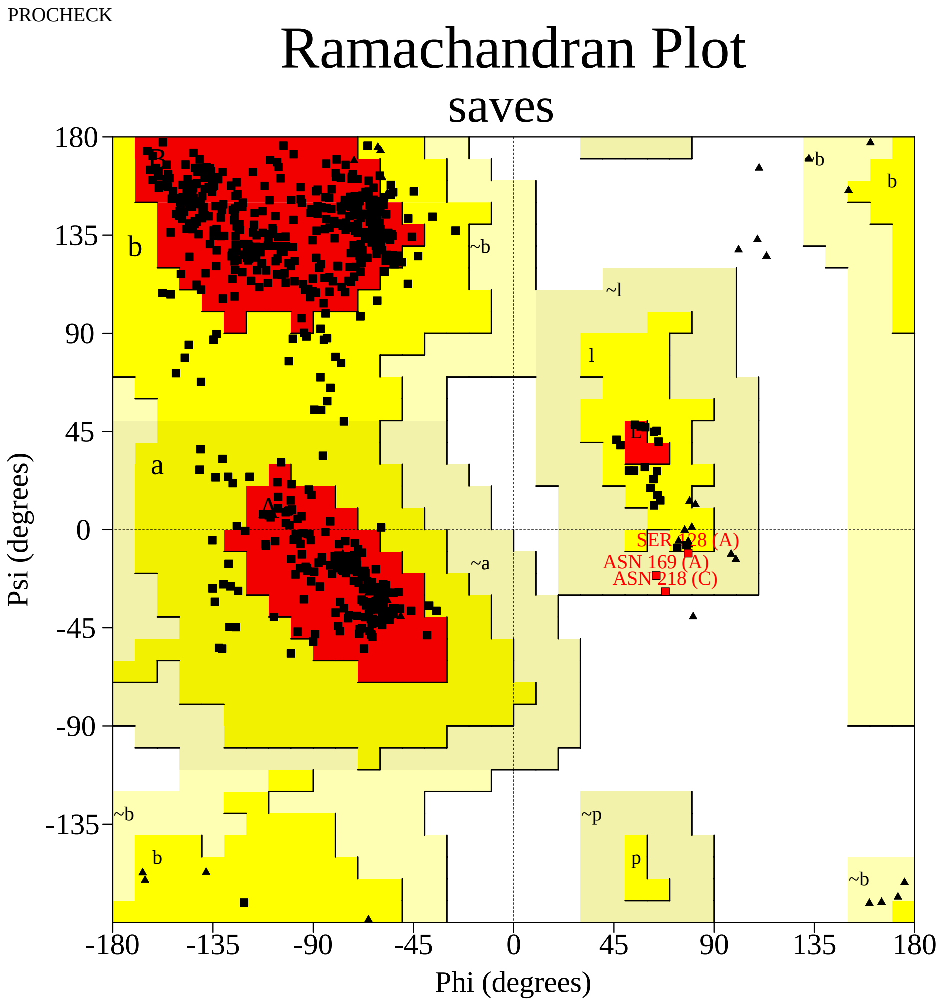


Plot statistics

| | | |
|---|---|---|
| Residues in most favoured regions [A,B,L] | 513 | 87.7% |
| Residues in additional allowed regions [a,b,l,p] | 69 | 11.8% |
| Residues in generously allowed regions [~a,~b,~l,~p] | 3 | 0.5% |
| Residues in disallowed regions | 0 | 0.0% |
|  | ---- | ------ |
| Number of non-glycine and non-proline residues | 585 | 100.0% |
| Number of end-residues (excl. Gly and Pro) | 6 | |
| Number of glycine residues (shown as triangles) | 36 | |
| Number of proline residues | 42 | |
|  | ---- |  |
| Total number of residues | 669 | |

Based on an analysis of 118 structures of resolution of at least 2.0 Angstroms and R-factor no greater than 20%, a good quality model would be expected to have over 90% in the most favoured regions.

saves_01.ps

Supplementary Figure S1 (E). The PROCHECK Ramachandran plot for the validation of the 3D model of Spike RBD of SARS-CoV-2 B.1.1.529(Omicron) variant (EPI_ISL_6640917)

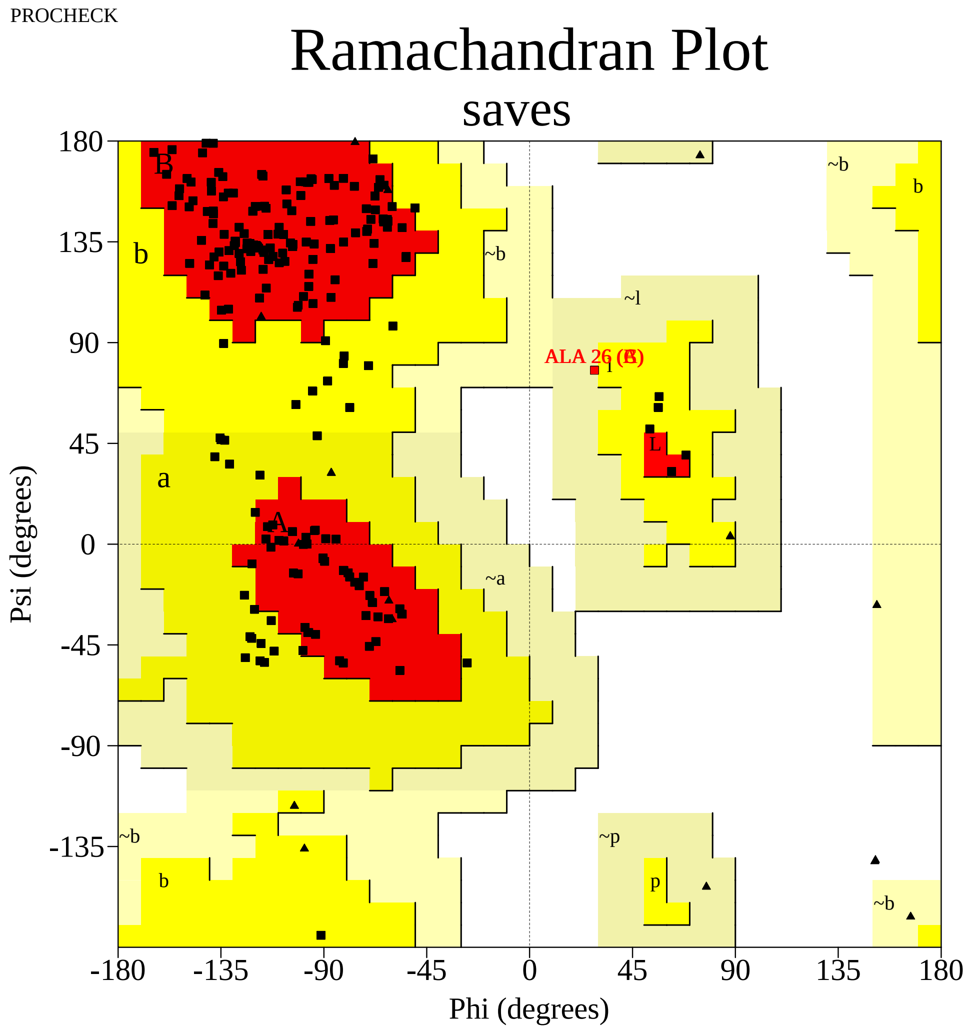


Plot statistics

| | | |
|---|---|---|
| Residues in most favoured regions [A,B,L] | 482 | 83.2% |
| Residues in additional allowed regions [a,b,l,p] | 94 | 16.2% |
| Residues in generously allowed regions [~a,~b,~l,~p] | 3 | 0.5% |
| Residues in disallowed regions | 0 | 0.0% |
|  | ---- | ------ |
| Number of non-glycine and non-proline residues | 579 | 100.0% |
| Number of end-residues (excl. Gly and Pro) | 6 | |
| Number of glycine residues (shown as triangles) | 45 | |
| Number of proline residues | 39 | |
|  | ---- |  |
| Total number of residues | 669 | |

Based on an analysis of 118 structures of resolution of at least 2.0 Angstroms
and R-factor no greater than 20%, a good quality model would be expected
to have over 90% in the most favoured regions.

saves_01.ps

Supplementary Figure S1 (F). The PROCHECK Ramachandran plot for the validation of the 3D model of Spike RBD of SARS-CoV-2 C.37(Lambda) variant (EPI_ISL_1111334)

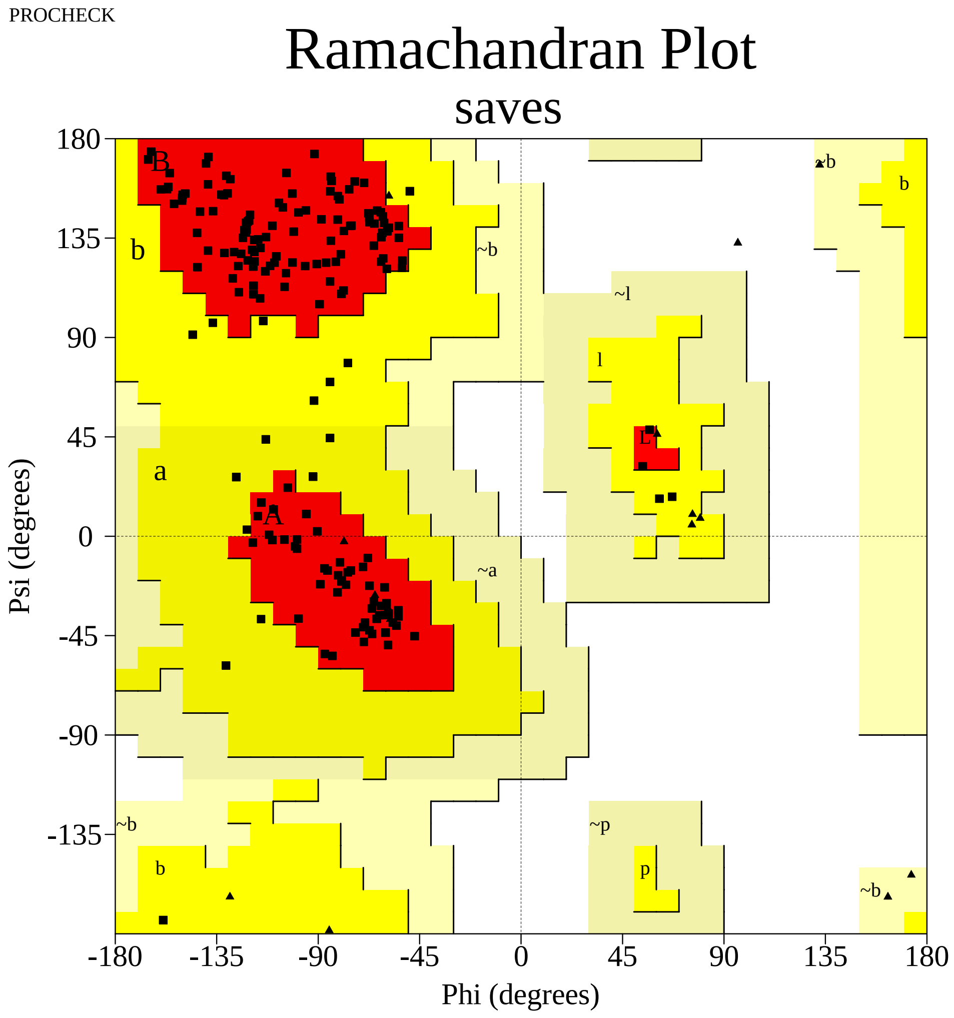


Plot statistics

| | | |
|---|---|---|
| Residues in most favoured regions [A,B,L] | 155 | 90.6% |
| Residues in additional allowed regions [a,b,l,p] | 16 | 9.4% |
| Residues in generously allowed regions [~a,~b,~l,~p] | 0 | 0.0% |
| Residues in disallowed regions | 0 | 0.0% |
|  | ---- | ------ |
| Number of non-glycine and non-proline residues | 171 | 100.0% |
| Number of end-residues (excl. Gly and Pro) | 2 | |
| Number of glycine residues (shown as triangles) | 15 | |
| Number of proline residues | 11 | |
|  | ---- |  |
| Total number of residues | 199 | |

Based on an analysis of 118 structures of resolution of at least 2.0 Angstroms and R-factor no greater than 20%, a good quality model would be expected to have over 90% in the most favoured regions.

saves_01.ps

Supplementary Figure S1 (G). The PROCHECK Ramachandran plot for the validation of the 3D model of Spike RBD of SARS-CoV-2 B.1.621(Mu) variant (EPI_ISL_3369952)

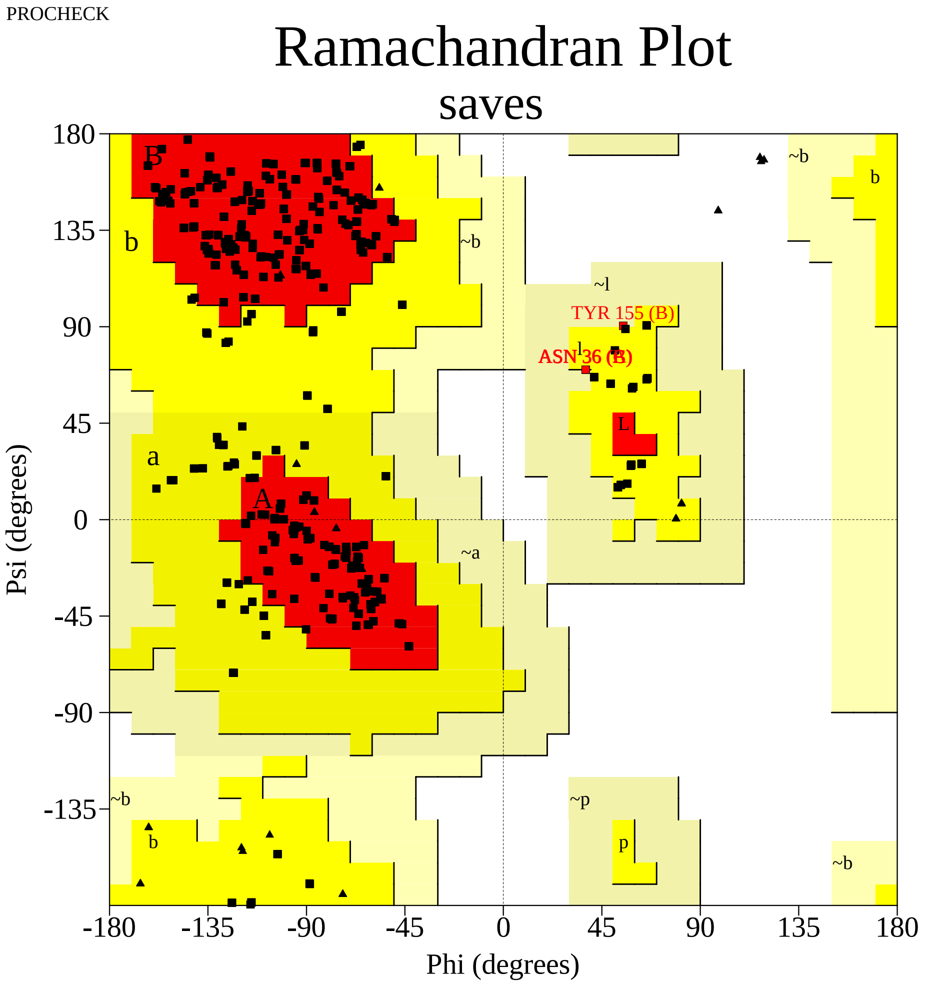


Plot statistics

| | | |
|---|---|---|
| Residues in most favoured regions [A,B,L] | 454 | 78.4% |
| Residues in additional allowed regions [a,b,l,p] | 121 | 20.9% |
| Residues in generously allowed regions [~a,~b,~l,~p] | 4 | 0.7% |
| Residues in disallowed regions | 0 | 0.0% |
|  | ---- | ------ |
| Number of non-glycine and non-proline residues | 579 | 100.0% |
| Number of end-residues (excl. Gly and Pro) | 6 | |
| Number of glycine residues (shown as triangles) | 45 | |
| Number of proline residues | 39 | |
|  | ---- |  |
| Total number of residues | 669 | |

Based on an analysis of 118 structures of resolution of at least 2.0 Angstroms
and R-factor no greater than 20%, a good quality model would be expected
to have over 90% in the most favoured regions.

saves_01.ps

Supplementary Figure S1 (H). The PROCHECK Ramachandran plot for the validation of the 3D model of Spike RBD of SARS-CoV-2 B.1.427/B.1.429(Epsilon) variant (EPI_ISL_648527)

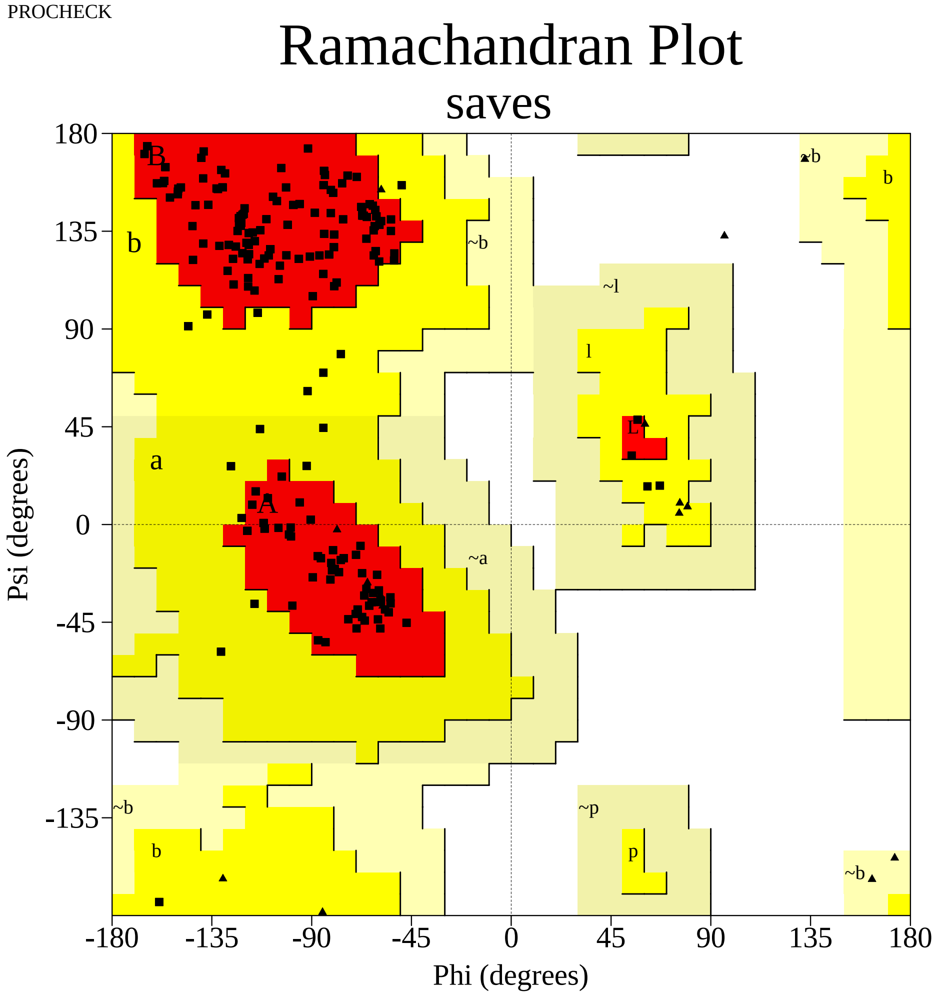


Plot statistics

| | | |
|---|---|---|
| Residues in most favoured regions [A,B,L] | 155 | 90.6% |
| Residues in additional allowed regions [a,b,l,p] | 16 | 9.4% |
| Residues in generously allowed regions [~a,~b,~l,~p] | 0 | 0.0% |
| Residues in disallowed regions | 0 | 0.0% |
|  | ---- | ------ |
| Number of non-glycine and non-proline residues | 171 | 100.0% |
| Number of end-residues (excl. Gly and Pro) | 2 | |
| Number of glycine residues (shown as triangles) | 15 | |
| Number of proline residues | 11 | |
|  | ---- |  |
| Total number of residues | 199 | |

Based on an analysis of 118 structures of resolution of at least 2.0 Angstroms and R-factor no greater than 20%, a good quality model would be expected to have over 90% in the most favoured regions.

saves_01.ps

Supplementary Figure S1 (I). The PROCHECK Ramachandran plot for the validation of the 3D model of Spike RBD of SARS-CoV-2 B.1.1.519 variant (EPI_ISL_721617)

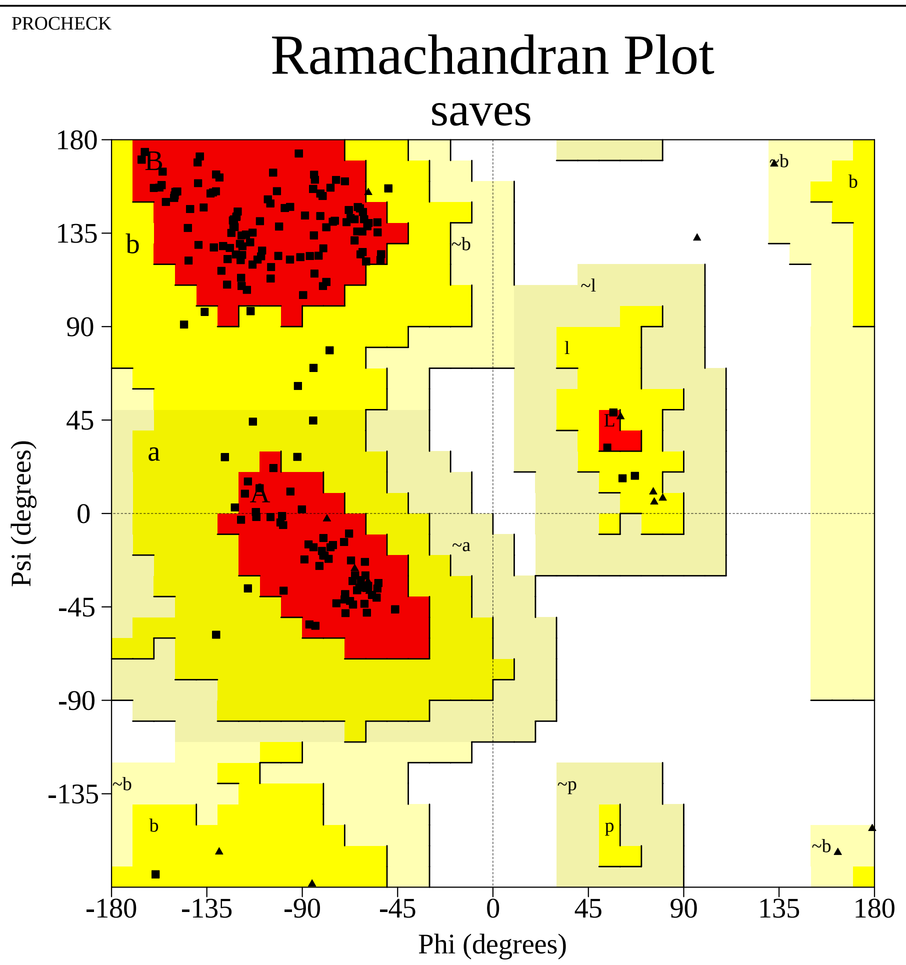


Plot statistics

| | | |
|---|---|---|
| Residues in most favoured regions [A,B,L] | 155 | 90.6% |
| Residues in additional allowed regions [a,b,l,p] | 16 | 9.4% |
| Residues in generously allowed regions [~a,~b,~l,~p] | 0 | 0.0% |
| Residues in disallowed regions | 0 | 0.0% |
|  | ---- | ------ |
| Number of non-glycine and non-proline residues | 171 | 100.0% |
| Number of end-residues (excl. Gly and Pro) | 2 | |
| Number of glycine residues (shown as triangles) | 15 | |
| Number of proline residues | 11 | |
|  | ---- |  |
| Total number of residues | 199 | |

Based on an analysis of 118 structures of resolution of at least 2.0 Angstroms and R-factor no greater than 20%, a good quality model would be expected to have over 90% in the most favoured regions.

saves_01.ps

Supplementary Figure S1 (J). The PROCHECK Ramachandran plot for the validation of the 3D model of Spike RBD of SARS-CoV-2 R.1 variant (EPI_ISL_736897)

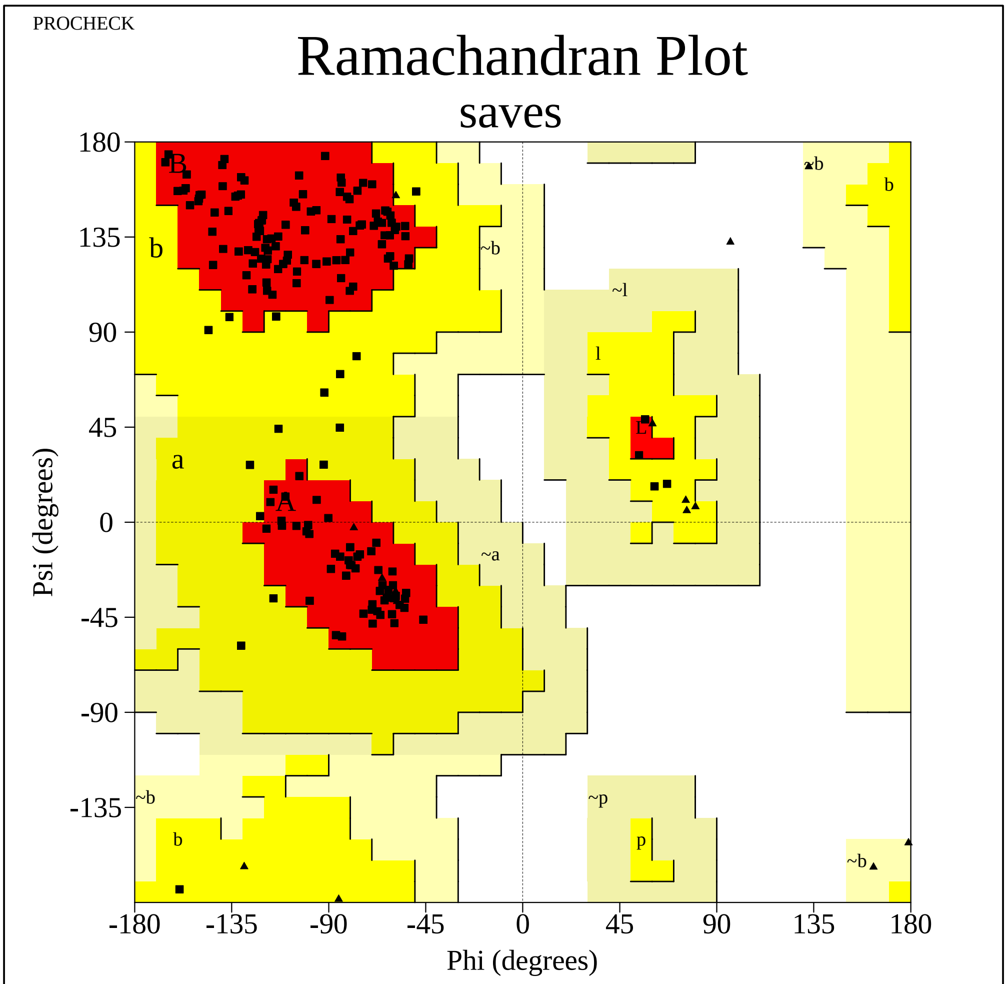


Plot statistics

| | | |
|---|---|---|
| Residues in most favoured regions [A,B,L] | 155 | 90.6% |
| Residues in additional allowed regions [a,b,l,p] | 16 | 9.4% |
| Residues in generously allowed regions [~a,~b,~l,~p] | 0 | 0.0% |
| Residues in disallowed regions | 0 | 0.0% |
|  | ---- | ------ |
| Number of non-glycine and non-proline residues | 171 | 100.0% |
| Number of end-residues (excl. Gly and Pro) | 2 | |
| Number of glycine residues (shown as triangles) | 15 | |
| Number of proline residues | 11 | |
|  | ---- |  |
| Total number of residues | 199 | |

Based on an analysis of 118 structures of resolution of at least 2.0 Angstroms and R-factor no greater than 20%, a good quality model would be expected to have over 90% in the most favoured regions.

saves_01.ps

Supplementary Figure S1 (K). The PROCHECK Ramachandran plot for the validation of the 3D model of Spike RBD of SARS-CoV-2 B.1.525(Eta) variant (EPI_ISL_760883)

PROCHECK

### Ramachandran Plot

#### saves

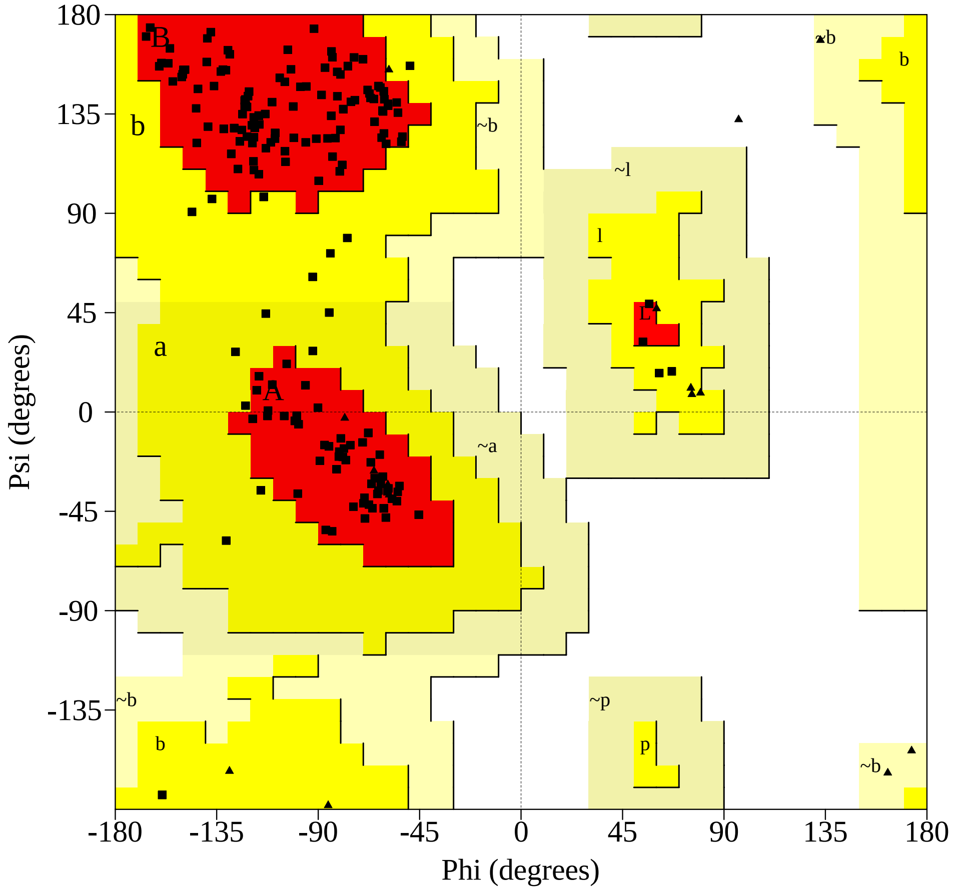


##### Plot statistics

| | | |
|---|---|---|
| Residues in most favoured regions [A,B,L] | 155 | 90.6% |
| Residues in additional allowed regions [a,b,l,p] | 16 | 9.4% |
| Residues in generously allowed regions [~a,~b,~l,~p] | 0 | 0.0% |
| Residues in disallowed regions | 0 | 0.0% |
|  | ---- | ------ |
| Number of non-glycine and non-proline residues | 171 | 100.0% |
| Number of end-residues (excl. Gly and Pro) | 2 | |
| Number of glycine residues (shown as triangles) | 15 | |
| Number of proline residues | 11 | |
|  | ---- |  |
| Total number of residues | 199 | |

Based on an analysis of 118 structures of resolution of at least 2.0 Angstroms
and R-factor no greater than 20%, a good quality model would be expected
to have over 90% in the most favoured regions.

saves_01.ps

Supplementary Figure S1 (L). The PROCHECK Ramachandran plot for the validation of the 3D model of Spike RBD of SARS-CoV-2 B.1.214.2 variant (EPI_ISL_760951)

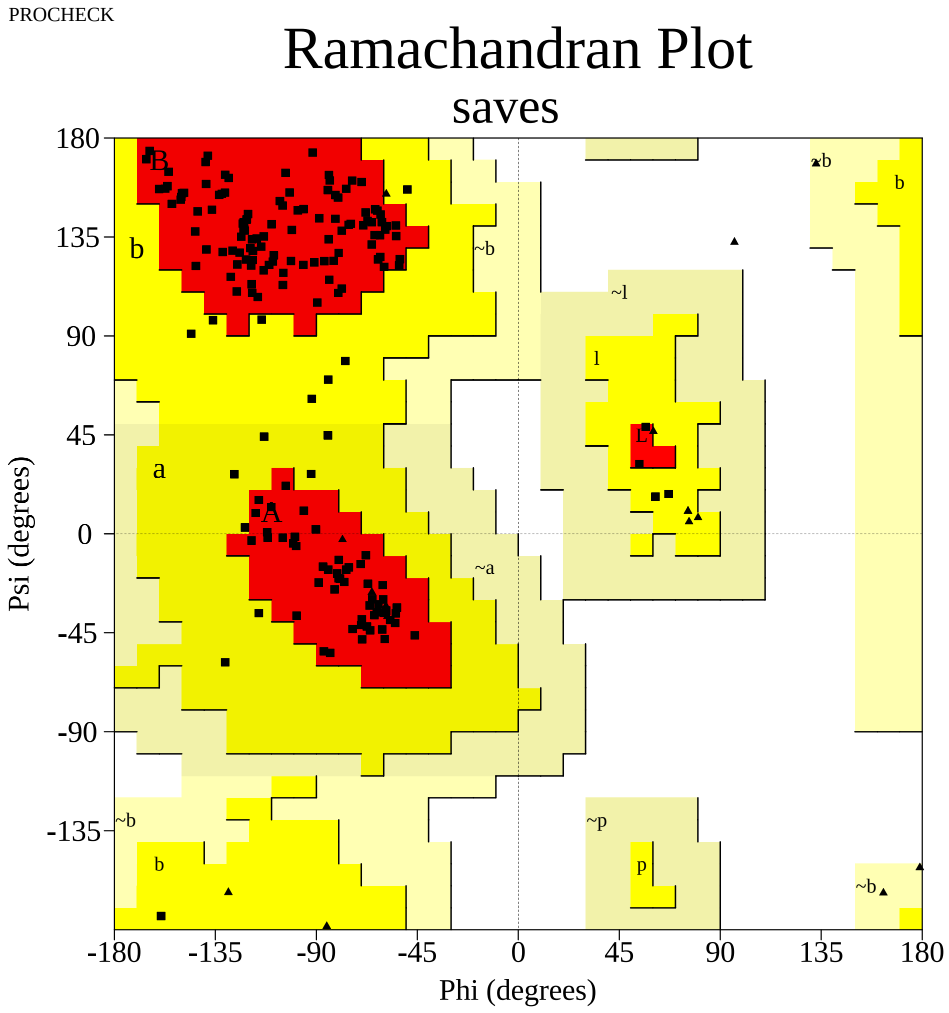


Plot statistics

| | | |
|---|---|---|
| Residues in most favoured regions [A,B,L] | 155 | 90.6% |
| Residues in additional allowed regions [a,b,l,p] | 16 | 9.4% |
| Residues in generously allowed regions [~a,~b,~l,~p] | 0 | 0.0% |
| Residues in disallowed regions | 0 | 0.0% |
|  | ---- | ------ |
| Number of non-glycine and non-proline residues | 171 | 100.0% |
| Number of end-residues (excl. Gly and Pro) | 2 | |
| Number of glycine residues (shown as triangles) | 15 | |
| Number of proline residues | 11 | |
|  | ---- |  |
| Total number of residues | 199 | |

Based on an analysis of 118 structures of resolution of at least 2.0 Angstroms and R-factor no greater than 20%, a good quality model would be expected to have over 90% in the most favoured regions.

saves_01.ps

Supplementary Figure S1 (M). The PROCHECK Ramachandran plot for the validation of the 3D model of Spike RBD of SARS-CoV-2 B.1.526(Lota) variant (EPI_ISL_765494)

PROCHECK

### Ramachandran Plot

#### saves

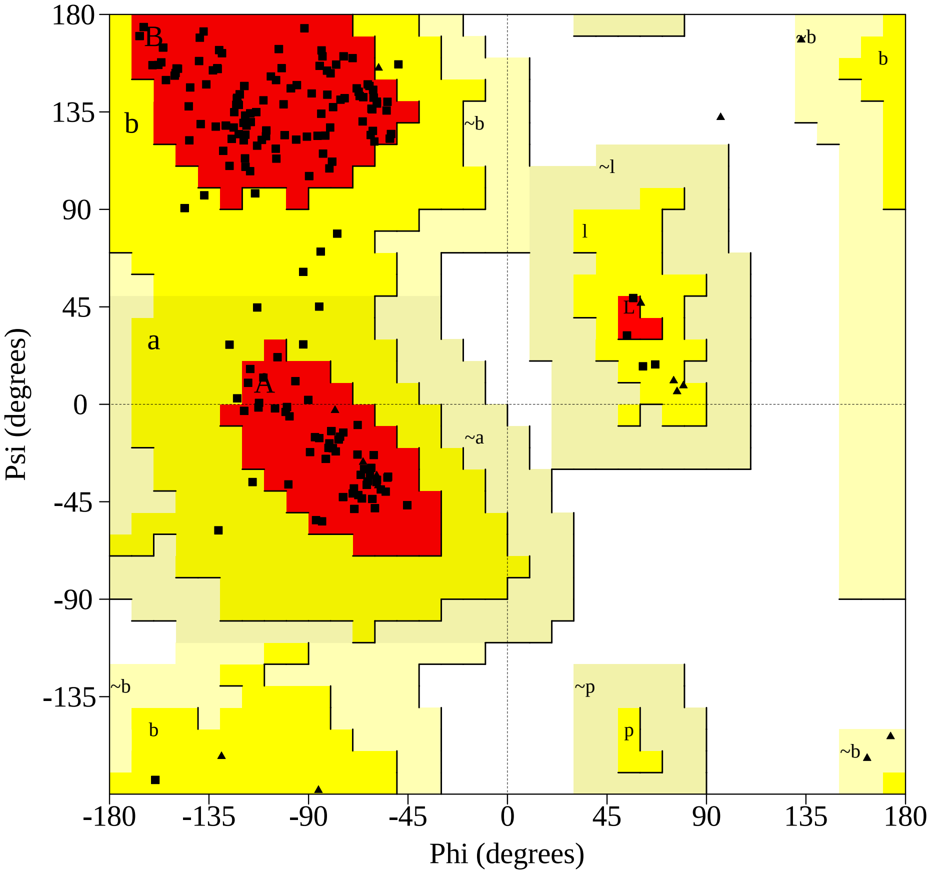


Plot statistics

| | | |
|---|---|---|
| Residues in most favoured regions [A,B,L] | 155 | 90.6% |
| Residues in additional allowed regions [a,b,l,p] | 16 | 9.4% |
| Residues in generously allowed regions [~a,~b,~l,~p] | 0 | 0.0% |
| Residues in disallowed regions | 0 | 0.0% |
|  | ---- | ------ |
| Number of non-glycine and non-proline residues | 171 | 100.0% |
| Number of end-residues (excl. Gly and Pro) | 2 | |
| Number of glycine residues (shown as triangles) | 15 | |
| Number of proline residues | 11 | |
|  | ---- |  |
| Total number of residues | 199 | |

Based on an analysis of 118 structures of resolution of at least 2.0 Angstroms
and R-factor no greater than 20%, a good quality model would be expected
to have over 90% in the most favoured regions.

saves_01.ps

Supplementary Figure S1 (N). The PROCHECK Ramachandran plot for the validation of the 3D model of Spike RBD of SARS-CoV-2 B.1.466.2 variant (EPI_ISL_877419)

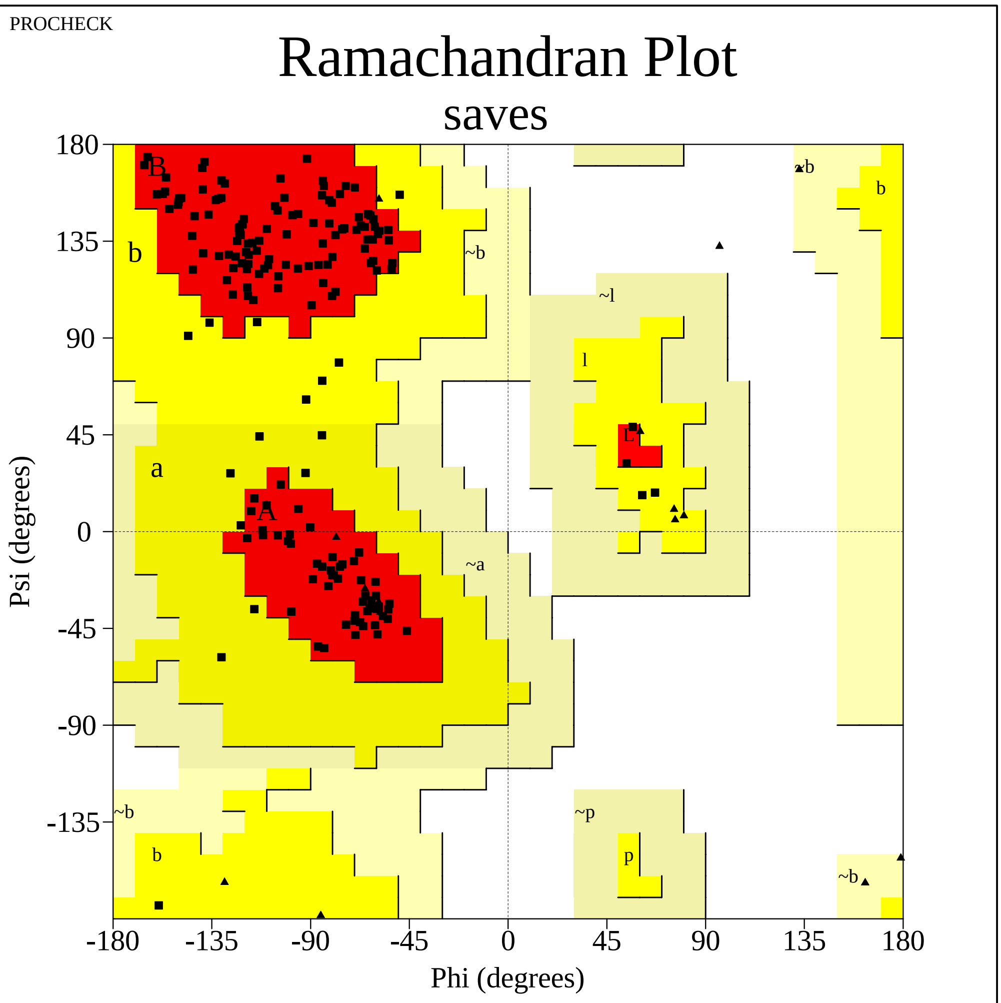


Plot statistics

| | | |
|---|---|---|
| Residues in most favoured regions [A,B,L] | 155 | 90.6% |
| Residues in additional allowed regions [a,b,l,p] | 16 | 9.4% |
| Residues in generously allowed regions [~a,~b,~l,~p] | 0 | 0.0% |
| Residues in disallowed regions | 0 | 0.0% |
|  | ---- | ------ |
| Number of non-glycine and non-proline residues | 171 | 100.0% |
| Number of end-residues (excl. Gly and Pro) | 2 | |
| Number of glycine residues (shown as triangles) | 15 | |
| Number of proline residues | 11 | |
|  | ---- |  |
| Total number of residues | 199 | |

Based on an analysis of 118 structures of resolution of at least 2.0 Angstroms and R-factor no greater than 20%, a good quality model would be expected to have over 90% in the most favoured regions.

saves_01.ps

Supplementary Figure S1 (O). The PROCHECK Ramachandran plot for the validation of the 3D model of Spike RBD of SARS-CoV-2 B.1.1.318 variant (EPI_ISL_937654)

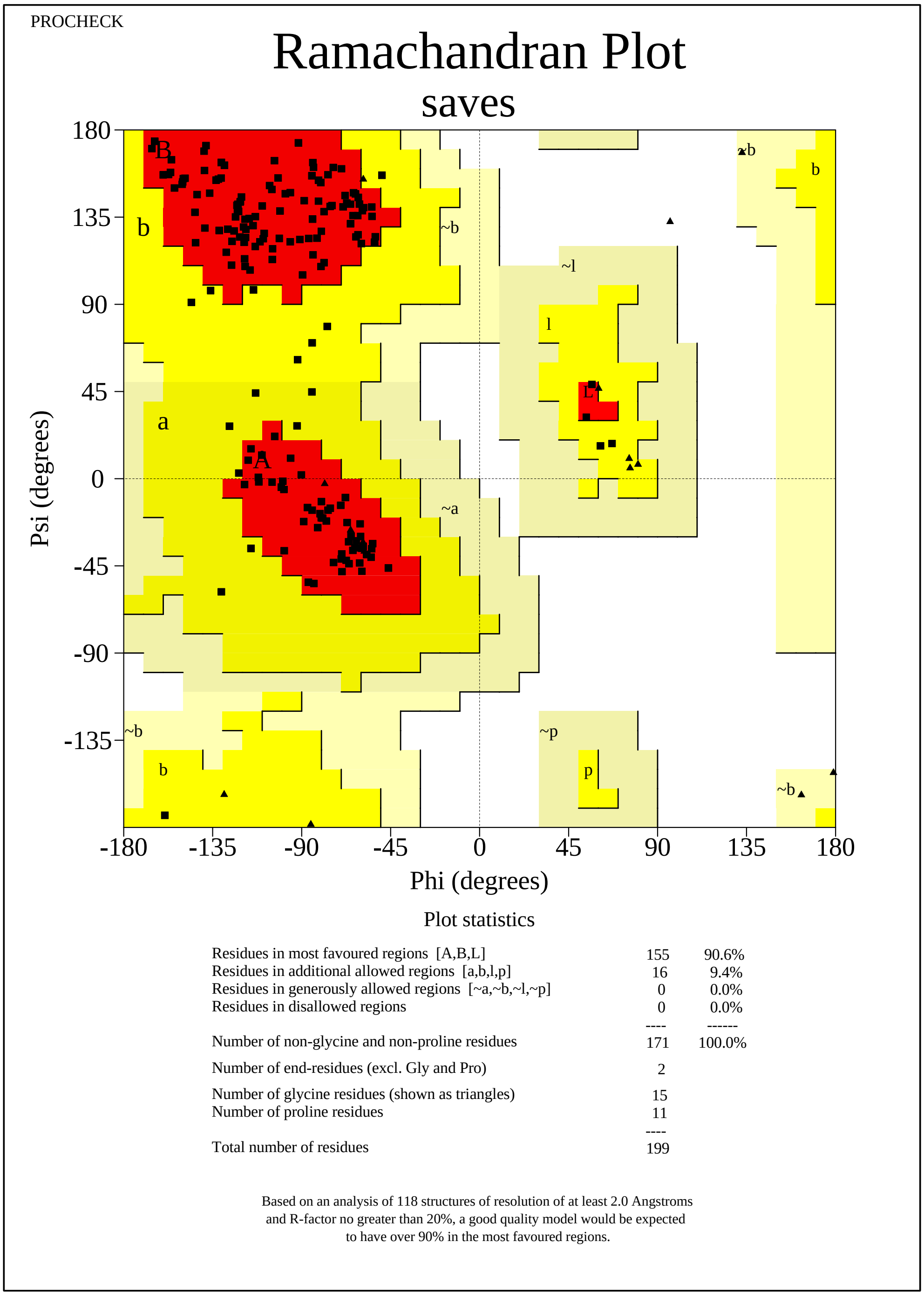


saves_01.ps

Supplementary Figure S1 (P). The PROCHECK Ramachandran plot for the validation of the 3D model of Spike RBD of SARS-CoV-2 B.1.619 variant (EPI_ISL_1150929)

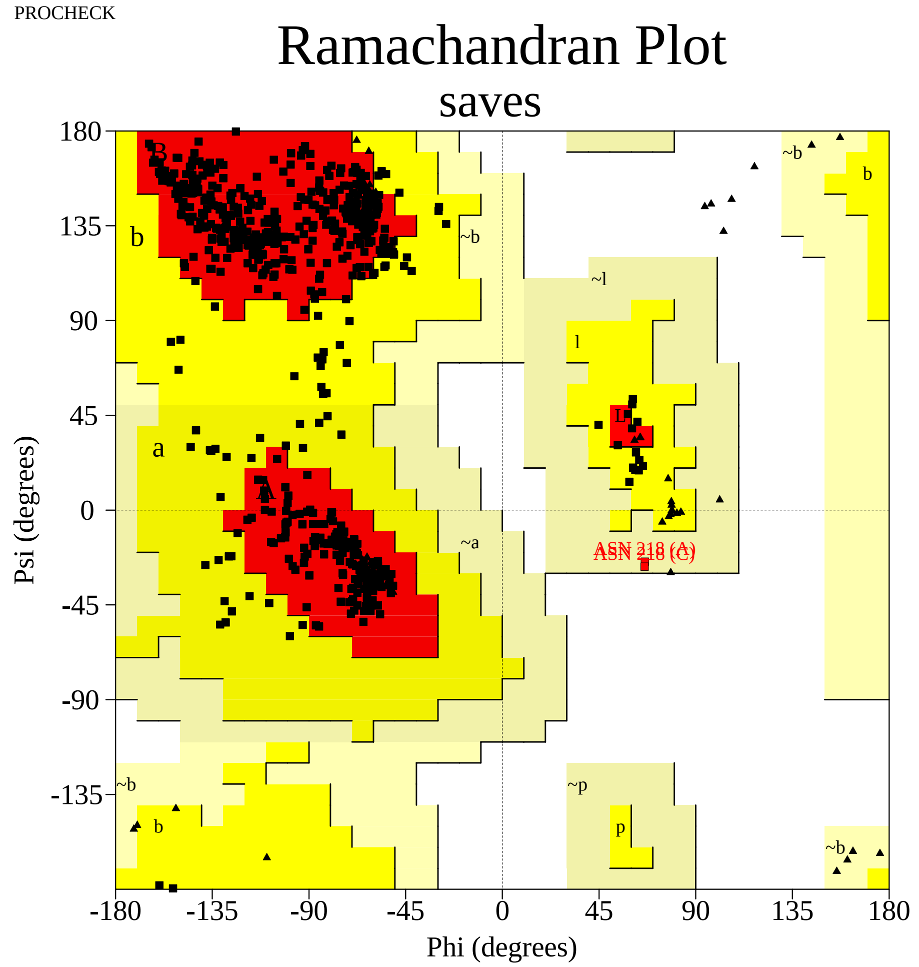


Plot statistics

| | | |
|---|---|---|
| Residues in most favoured regions [A,B,L] | 512 | 88.4% |
| Residues in additional allowed regions [a,b,l,p] | 65 | 11.2% |
| Residues in generously allowed regions [~a,~b,~l,~p] | 2 | 0.3% |
| Residues in disallowed regions | 0 | 0.0% |
|  | ---- | ------ |
| Number of non-glycine and non-proline residues | 579 | 100.0% |
| Number of end-residues (excl. Gly and Pro) | 6 | |
| Number of glycine residues (shown as triangles) | 45 | |
| Number of proline residues | 39 | |
|  | ---- |  |
| Total number of residues | 669 | |

Based on an analysis of 118 structures of resolution of at least 2.0 Angstroms and R-factor no greater than 20%, a good quality model would be expected to have over 90% in the most favoured regions.

saves_01.ps

Supplementary Figure S1 (Q). The PROCHECK Ramachandran plot for the validation of the 3D model of Spike RBD of SARS-CoV-2 C.36.3 variant (EPI_ISL_1237137)

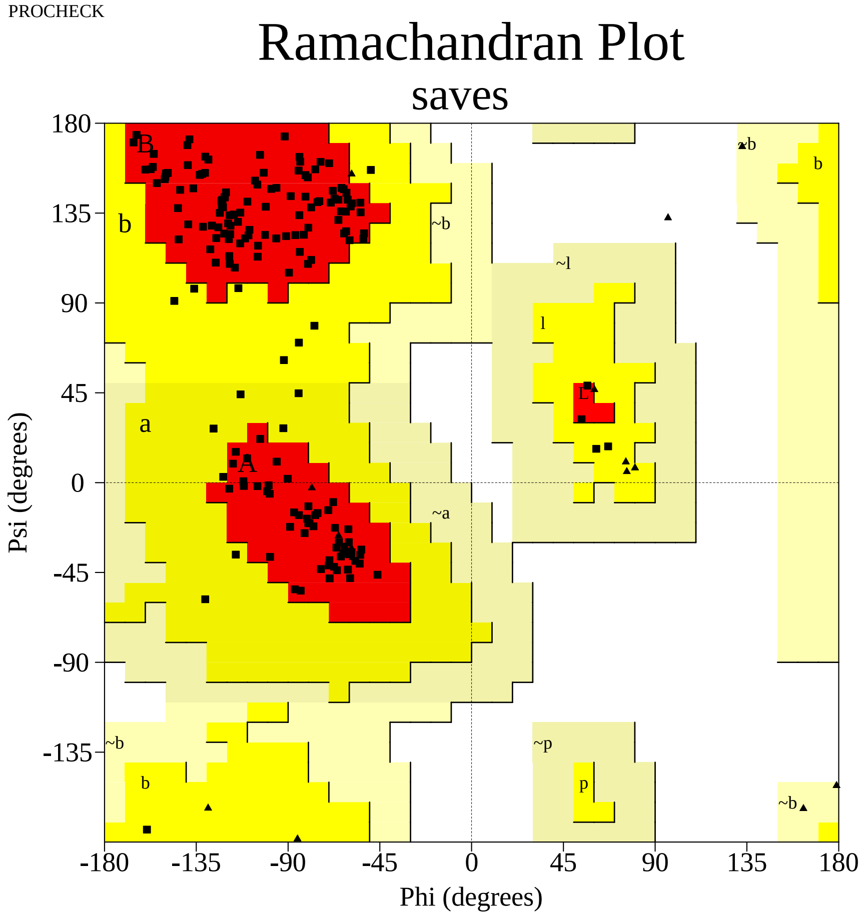


#### Plot statistics

| | | |
|---|---|---|
| Residues in most favoured regions [A,B,L] | 155 | 90.6% |
| Residues in additional allowed regions [a,b,l,p] | 16 | 9.4% |
| Residues in generously allowed regions [~a,~b,~l,~p] | 0 | 0.0% |
| Residues in disallowed regions | 0 | 0.0% |
|  | ---- | ------ |
| Number of non-glycine and non-proline residues | 171 | 100.0% |
| Number of end-residues (excl. Gly and Pro) | 2 | |
| Number of glycine residues (shown as triangles) | 15 | |
| Number of proline residues | 11 | |
|  | ---- |  |
| Total number of residues | 199 | |

Based on an analysis of 118 structures of resolution of at least 2.0 Angstroms
and R-factor no greater than 20%, a good quality model would be expected
to have over 90% in the most favoured regions.

saves_01.ps

Supplementary Figure S1 (R). The PROCHECK Ramachandran plot for the validation of the 3D model of Spike RBD of SARS-CoV-2 AT.1 variant (EPI_ISL_1259283)

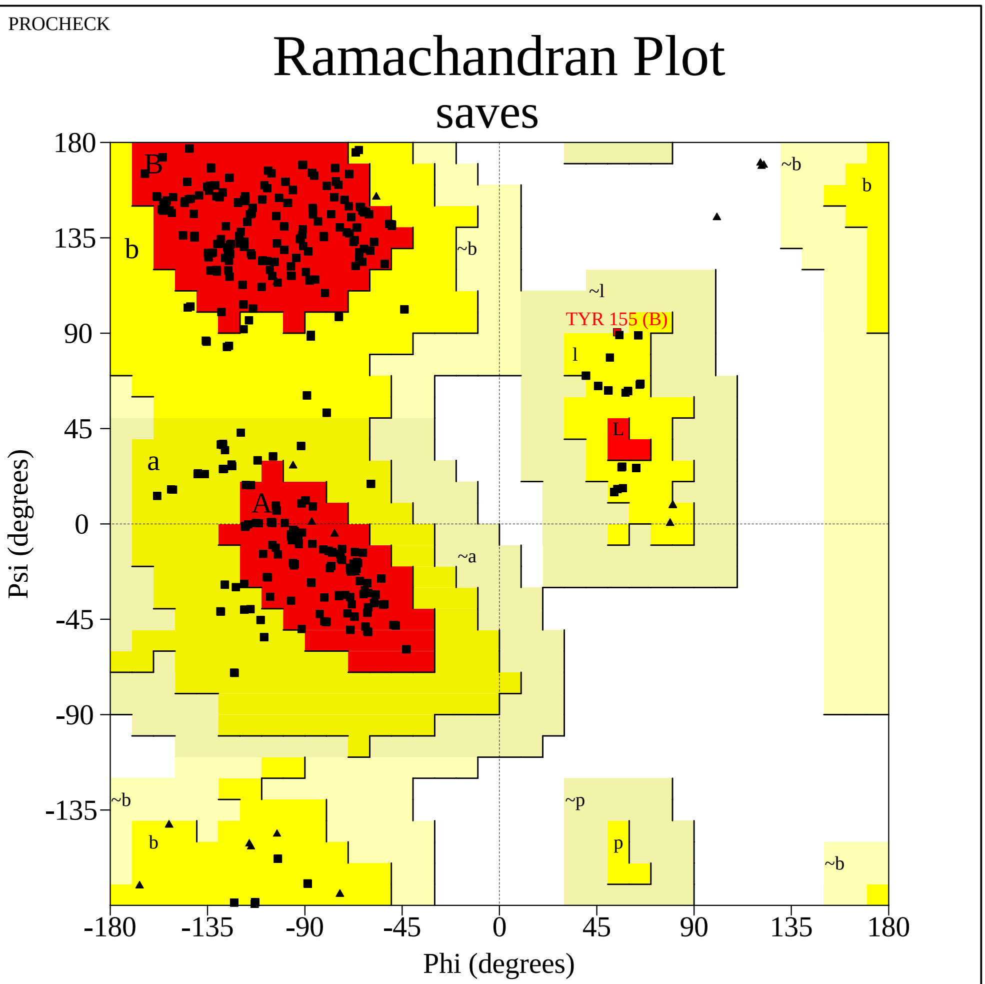


Plot statistics

| | | |
|---|---|---|
| Residues in most favoured regions [A,B,L] | 455 | 78.6% |
| Residues in additional allowed regions [a,b,l,p] | 123 | 21.2% |
| Residues in generously allowed regions [~a,~b,~l,~p] | 1 | 0.2% |
| Residues in disallowed regions | 0 | 0.0% |
|  | ---- | ------ |
| Number of non-glycine and non-proline residues | 579 | 100.0% |
| Number of end-residues (excl. Gly and Pro) | 6 | |
| Number of glycine residues (shown as triangles) | 45 | |
| Number of proline residues | 39 | |
|  | ---- |  |
| Total number of residues | 669 | |

Based on an analysis of 118 structures of resolution of at least 2.0 Angstroms and R-factor no greater than 20%, a good quality model would be expected to have over 90% in the most favoured regions.

saves_01.ps

Supplementary Figure S1 (S). The PROCHECK Ramachandran plot for the validation of the 3D model of Spike RBD of SARS-CoV-2 B.1.617.1(Kappa) variant (EPI_ISL_1357699)

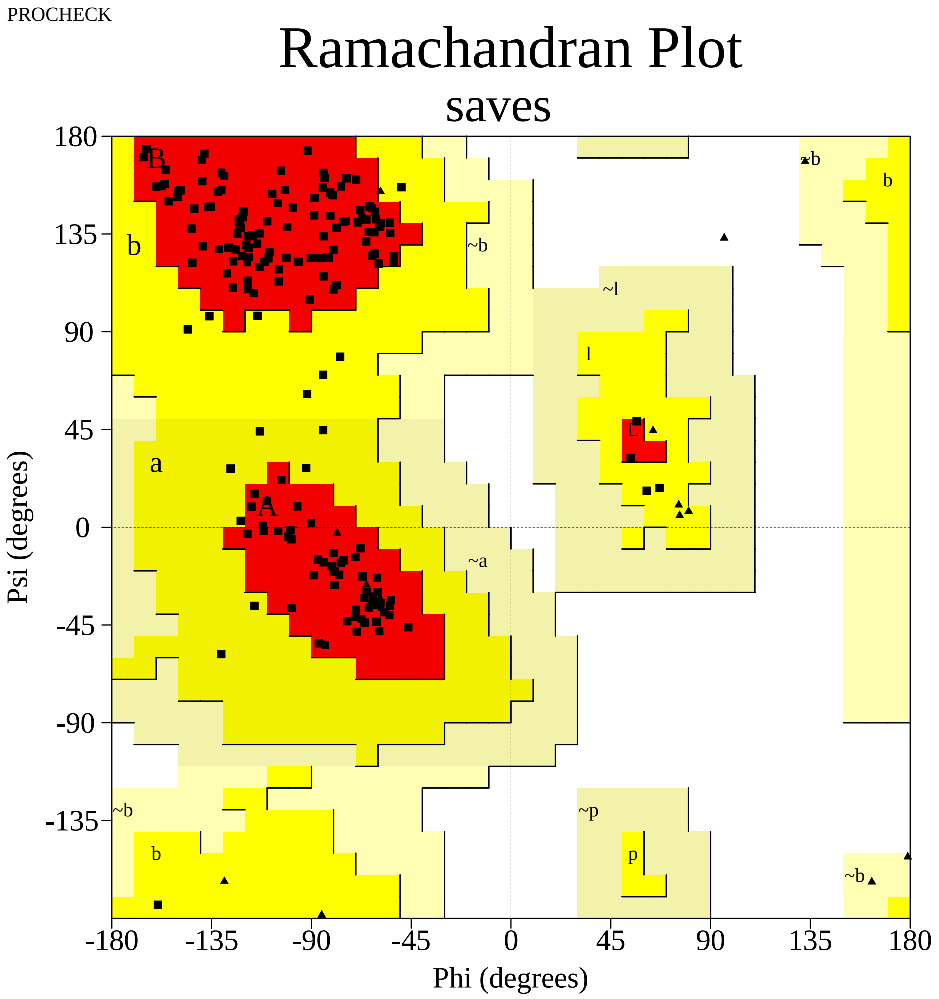


Plot statistics

| | | |
|---|---|---|
| Residues in most favoured regions [A,B,L] | 154 | 90.6% |
| Residues in additional allowed regions [a,b,l,p] | 16 | 9.4% |
| Residues in generously allowed regions [~a,~b,~l,~p] | 0 | 0.0% |
| Residues in disallowed regions | 0 | 0.0% |
|  | ---- | ------ |
| Number of non-glycine and non-proline residues | 170 | 100.0% |
| Number of end-residues (excl. Gly and Pro) | 2 | |
| Number of glycine residues (shown as triangles) | 15 | |
| Number of proline residues | 12 | |
|  | ---- |  |
| Total number of residues | 199 | |

Based on an analysis of 118 structures of resolution of at least 2.0 Angstroms and R-factor no greater than 20%, a good quality model would be expected to have over 90% in the most favoured regions.

saves_01.ps

Supplementary Figure S1 (T). The PROCHECK Ramachandran plot for the validation of the 3D model of Spike RBD of SARS-CoV-2 B.1.1.523 variant (EPI_ISL_1448584)

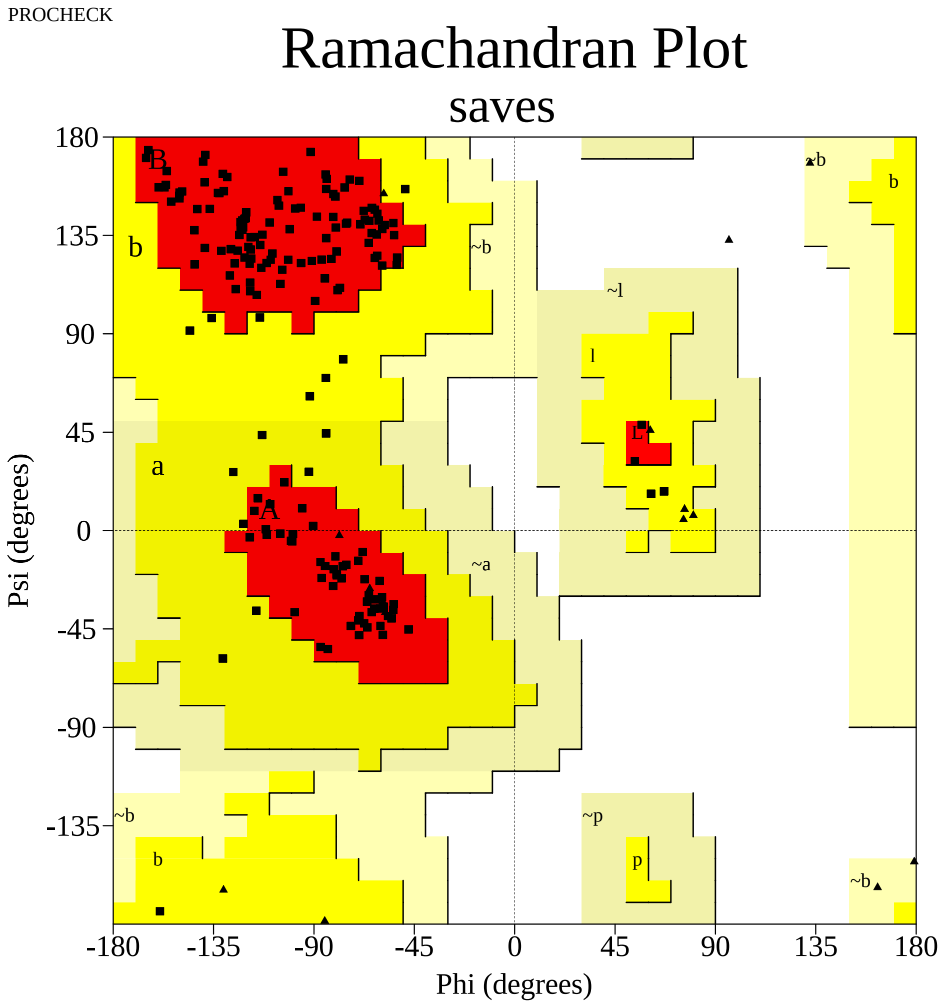


Plot statistics

| | | |
|---|---|---|
| Residues in most favoured regions [A,B,L] | 155 | 90.6% |
| Residues in additional allowed regions [a,b,l,p] | 16 | 9.4% |
| Residues in generously allowed regions [~a,~b,~l,~p] | 0 | 0.0% |
| Residues in disallowed regions | 0 | 0.0% |
|  | ---- | ------ |
| Number of non-glycine and non-proline residues | 171 | 100.0% |
| Number of end-residues (excl. Gly and Pro) | 2 | |
| Number of glycine residues (shown as triangles) | 15 | |
| Number of proline residues | 11 | |
|  | ---- |  |
| Total number of residues | 199 | |

Based on an analysis of 118 structures of resolution of at least 2.0 Angstroms and R-factor no greater than 20%, a good quality model would be expected to have over 90% in the most favoured regions.

saves_01.ps

Supplementary Figure S1 (U). The PROCHECK Ramachandran plot for the validation of the 3D model of Spike RBD of SARS-CoV-2 B.1.620 variant (EPI_ISL_1579527)

PROCHECK

### Ramachandran Plot

#### saves

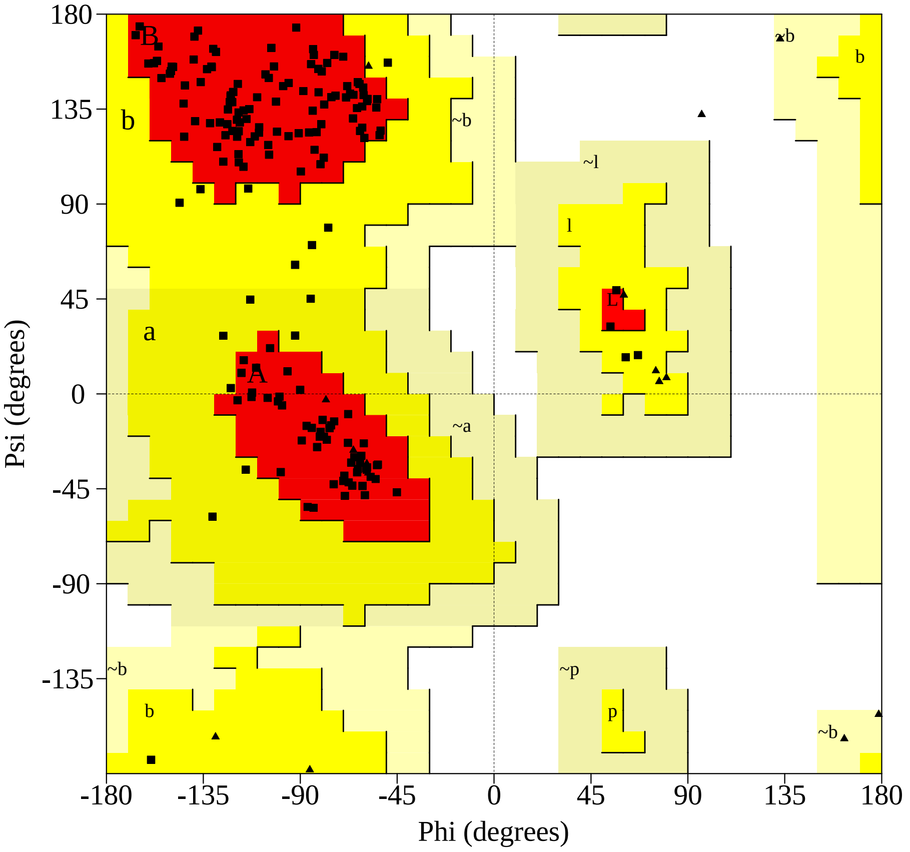


Plot statistics

| | | |
|---|---|---|
| Residues in most favoured regions [A,B,L] | 155 | 90.6% |
| Residues in additional allowed regions [a,b,l,p] | 16 | 9.4% |
| Residues in generously allowed regions [~a,~b,~l,~p] | 0 | 0.0% |
| Residues in disallowed regions | 0 | 0.0% |
|  | ---- | ------ |
| Number of non-glycine and non-proline residues | 171 | 100.0% |
| Number of end-residues (excl. Gly and Pro) | 2 | |
| Number of glycine residues (shown as triangles) | 15 | |
| Number of proline residues | 11 | |
|  | ---- |  |
| Total number of residues | 199 | |

Based on an analysis of 118 structures of resolution of at least 2.0 Angstroms and R-factor no greater than 20%, a good quality model would be expected to have over 90% in the most favoured regions.

saves_01.ps

Supplementary Figure S1 (V). The PROCHECK Ramachandran plot for the validation of the 3D model of Spike RBD of SARS-CoV-2 AV.1 variant (EPI_ISL_1595332)

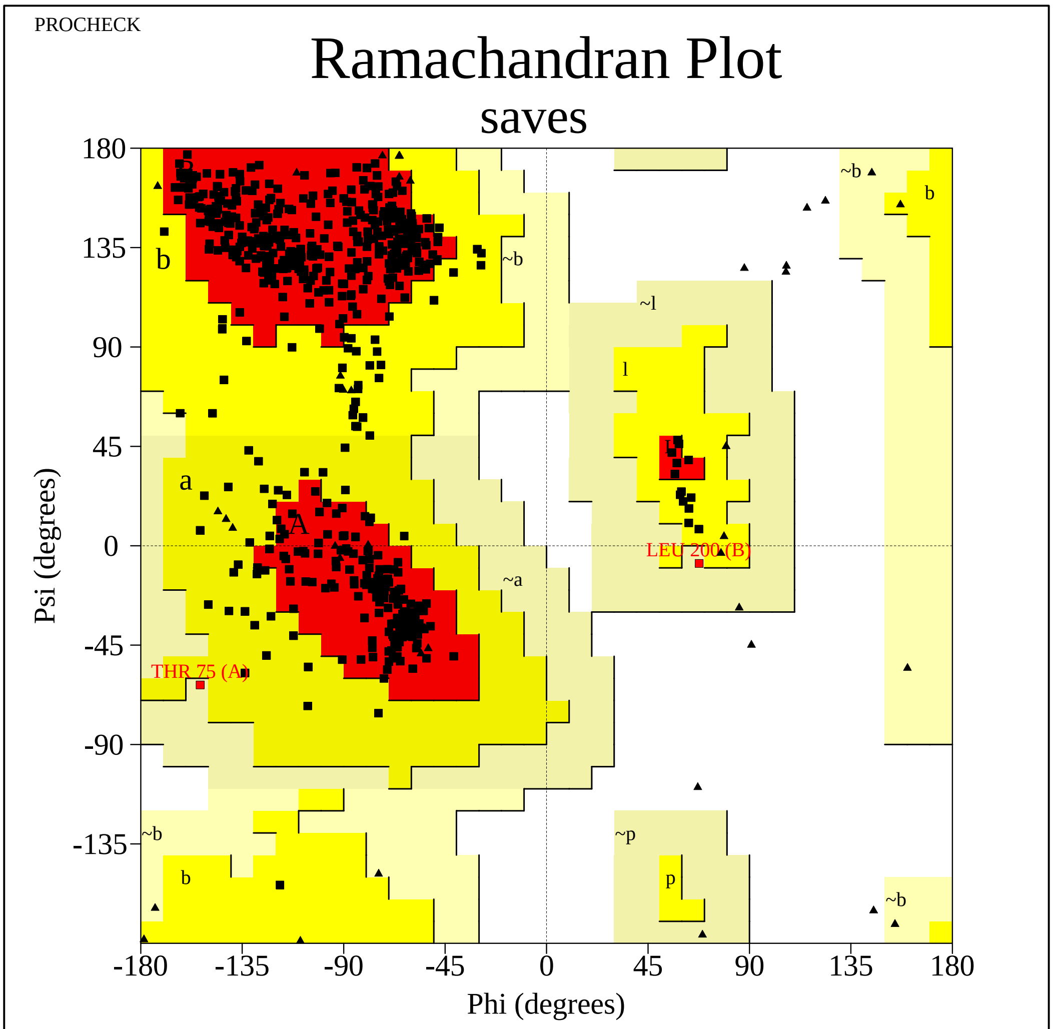


Plot statistics

| | | |
|---|---|---|
| Residues in most favoured regions [A,B,L] | 501 | 86.5% |
| Residues in additional allowed regions [a,b,l,p] | 76 | 13.1% |
| Residues in generously allowed regions [~a,~b,~l,~p] | 2 | 0.3% |
| Residues in disallowed regions | 0 | 0.0% |
|  | ---- | ------ |
| Number of non-glycine and non-proline residues | 579 | 100.0% |
| Number of end-residues (excl. Gly and Pro) | 6 | |
| Number of glycine residues (shown as triangles) | 45 | |
| Number of proline residues | 39 | |
|  | ---- |  |
| Total number of residues | 669 | |

Based on an analysis of 118 structures of resolution of at least 2.0 Angstroms and R-factor no greater than 20%, a good quality model would be expected to have over 90% in the most favoured regions.

saves_01.ps

Supplementary Figure S1 (W). The PROCHECK Ramachandran plot for the validation of the 3D model of Spike RBD of SARS-CoV-2 B.1.630 variant (EPI_ISL_3045385)

PROCHECK

### Ramachandran Plot

#### saves

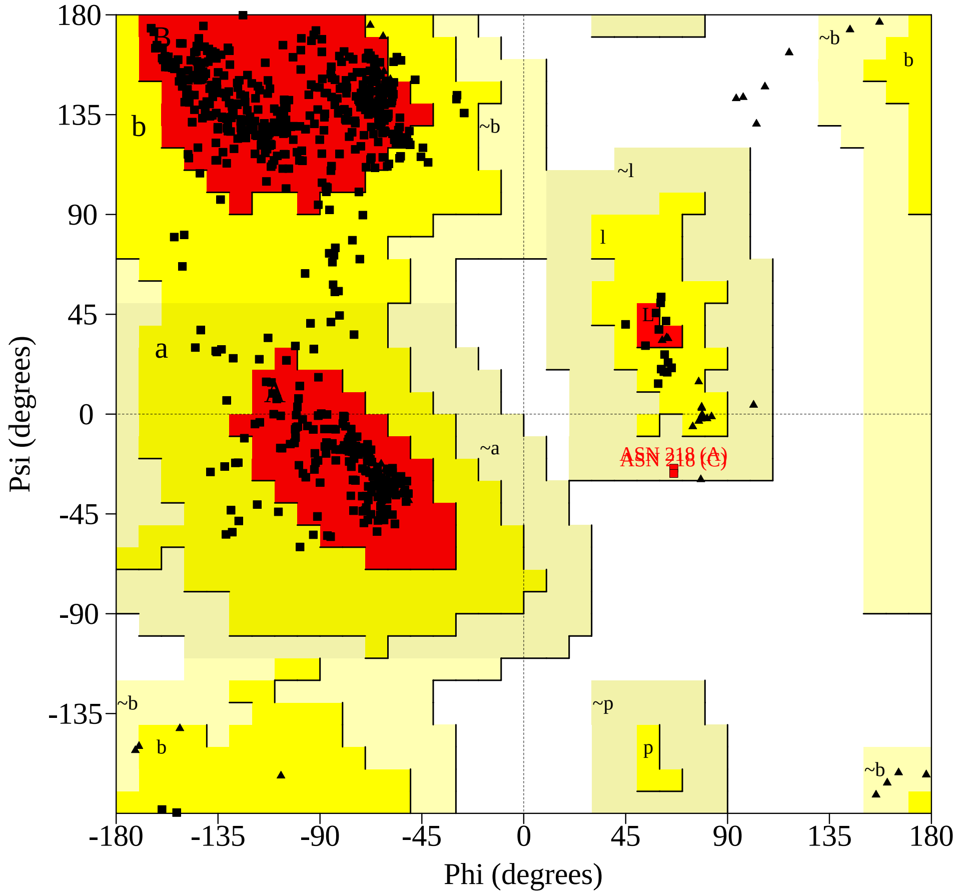


Plot statistics

| | | |
|---|---|---|
| Residues in most favoured regions [A,B,L] | 512 | 88.4% |
| Residues in additional allowed regions [a,b,l,p] | 65 | 11.2% |
| Residues in generously allowed regions [~a,~b,~l,~p] | 2 | 0.3% |
| Residues in disallowed regions | 0 | 0.0% |
|  | ---- | ------ |
| Number of non-glycine and non-proline residues | 579 | 100.0% |
| Number of end-residues (excl. Gly and Pro) | 6 | |
| Number of glycine residues (shown as triangles) | 45 | |
| Number of proline residues | 39 | |
|  | ---- |  |
| Total number of residues | 669 | |

Based on an analysis of 118 structures of resolution of at least 2.0 Angstroms and R-factor no greater than 20%, a good quality model would be expected to have over 90% in the most favoured regions.

saves_01.ps

Supplementary Figure S1 (X). The PROCHECK Ramachandran plot for the validation of the 3D model of Spike RBD of SARS-CoV-2 C.1.2 variant (EPI_ISL_3447714)

PROCHECK

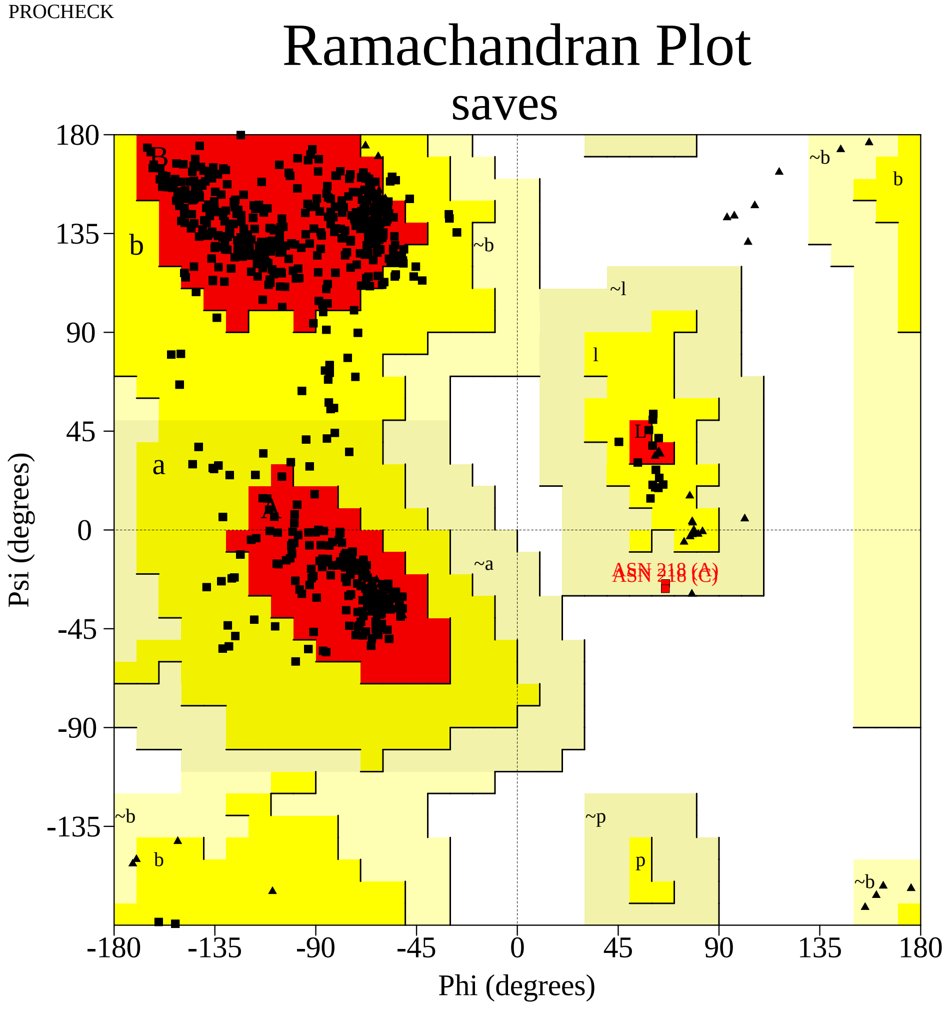


Plot statistics

| | | |
|---|---|---|
| Residues in most favoured regions [A,B,L] | 511 | 88.3% |
| Residues in additional allowed regions [a,b,l,p] | 66 | 11.4% |
| Residues in generously allowed regions [~a,~b,~l,~p] | 2 | 0.3% |
| Residues in disallowed regions | 0 | 0.0% |
|  | ---- | ------ |
| Number of non-glycine and non-proline residues | 579 | 100.0% |
| Number of end-residues (excl. Gly and Pro) | 6 | |
| Number of glycine residues (shown as triangles) | 45 | |
| Number of proline residues | 39 | |
|  | ---- |  |
| Total number of residues | 669 | |

Based on an analysis of 118 structures of resolution of at least 2.0 Angstroms and R-factor no greater than 20%, a good quality model would be expected to have over 90% in the most favoured regions.

saves_01.ps

Supplementary Figure S1 (Y). The PROCHECK Ramachandran plot for the validation of the 3D model of Spike RBD of SARS-CoV-2 B.1.640 variant (EPI_ISL_5592661)

**A**

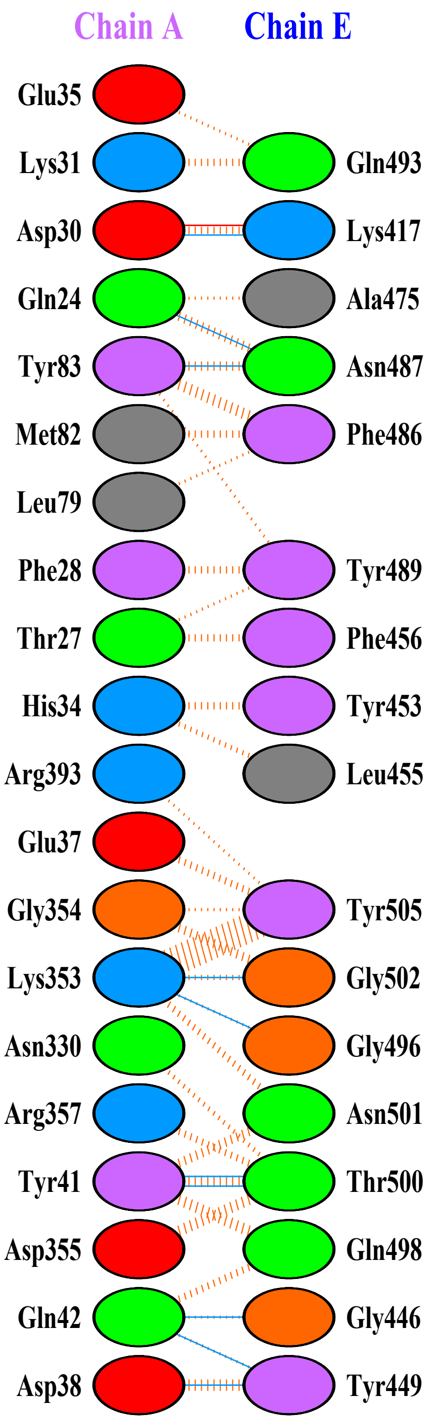


B

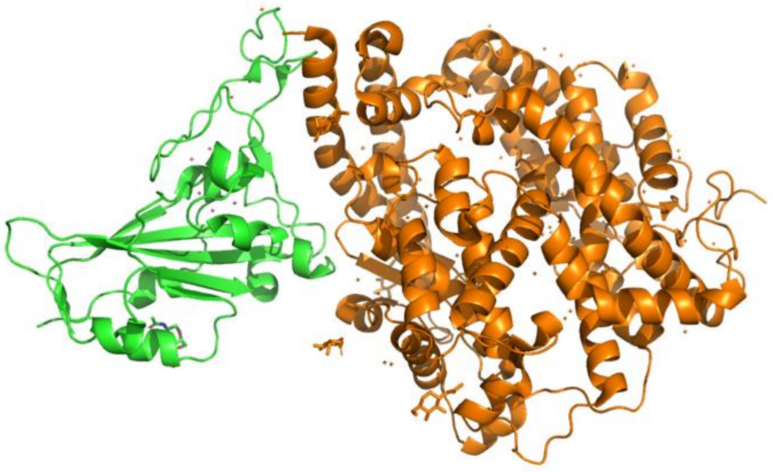

**Supplementary Figure:S2.A: Protein–protein docking representation of hACE2 and RBD of WIV04(EPI_ISL_402124)** . (A) The interaction representation, which includes hydrogen (▬▬), salt bridges (▬▬), and non-bonded (IIIIIII) interactions. Chain A represents hACE2 and chain B represents RBD . Re[illegible] colors: positive (blue): H,K,R; negative (red): D,E; neutral (green): S,T,N,Q; aliphatic (grey): A,V[illegible]M; aromatic (purple): F,Y,W; Pro & Gly (orange): P,G. (B) The binding interface of hACE2:RBD complex.

A

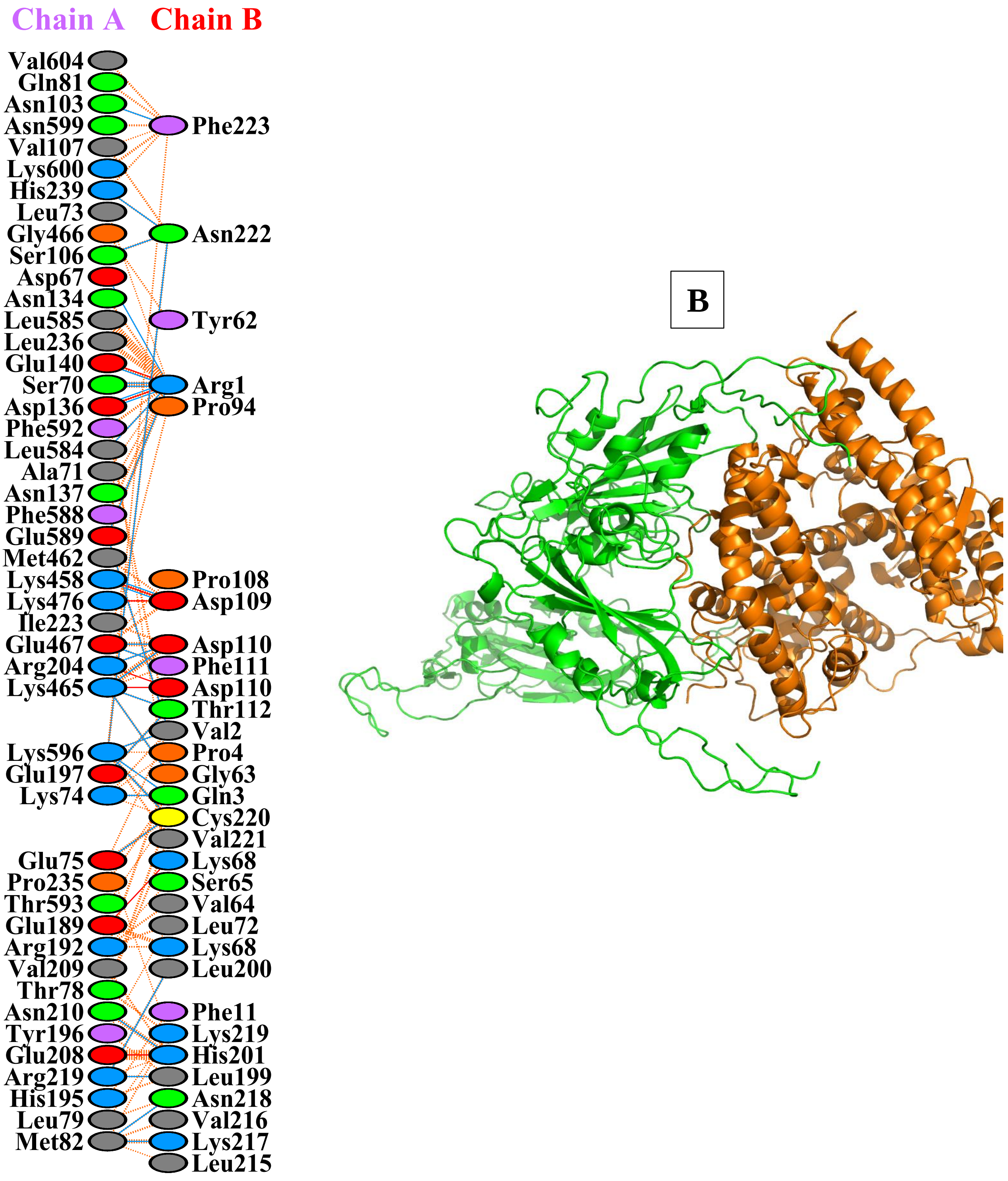


**Supplementary Figure:S2.B: Protein–protein docking representation of hACE2 and RBD of B.1.1.7(Alpha) (EPI_ISL_596982).** (A) The interaction representation, which includes hydrogen (blue line), salt bridges (red line), and non-bonded (orange dashes) interactions. Chain A represents hACE2 and chain B represents RBD . Residue colors: positive (blue): H,K,R; negative (red): D,E; neutral (green): S,T,N,Q; aliphatic (grey): A,V,L,I,M; aromatic (purple): F,Y,W; Pro & Gly (orange): P,[illegible] A The binding interface of hACE2:RBD complex.

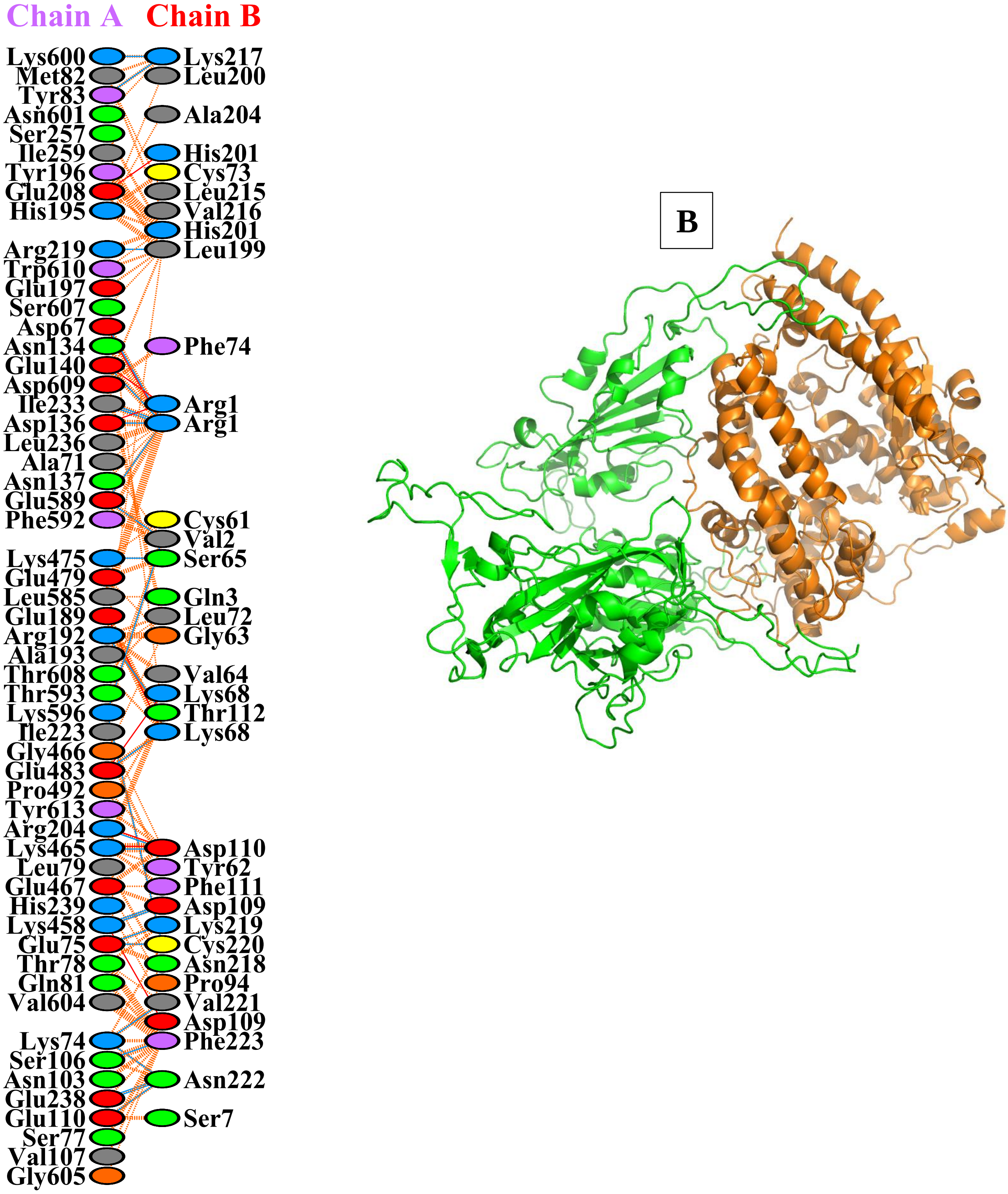


**Supplementary Figure:S2.C: Protein–protein docking representation of hACE2 and RBD of B.1.351(Beta) (EPI_ISL_660629).** (A) The interaction representation, which includes hydrogen ( ), salt bridges ( ), and non-bonded ( ) interactions. Chain A represents hACE2 and chain B represents RBD . Residue colors: positive (blue): H,K,R; negative (re [illegible] E; neutral (green): S,T,N,Q; aliphatic (grey): A,V,L,I,M; aromatic (purple): F,Y,W; Pro & Gly (orange [illegible]. (B) The binding interface of hACE2:RBD complex.

A

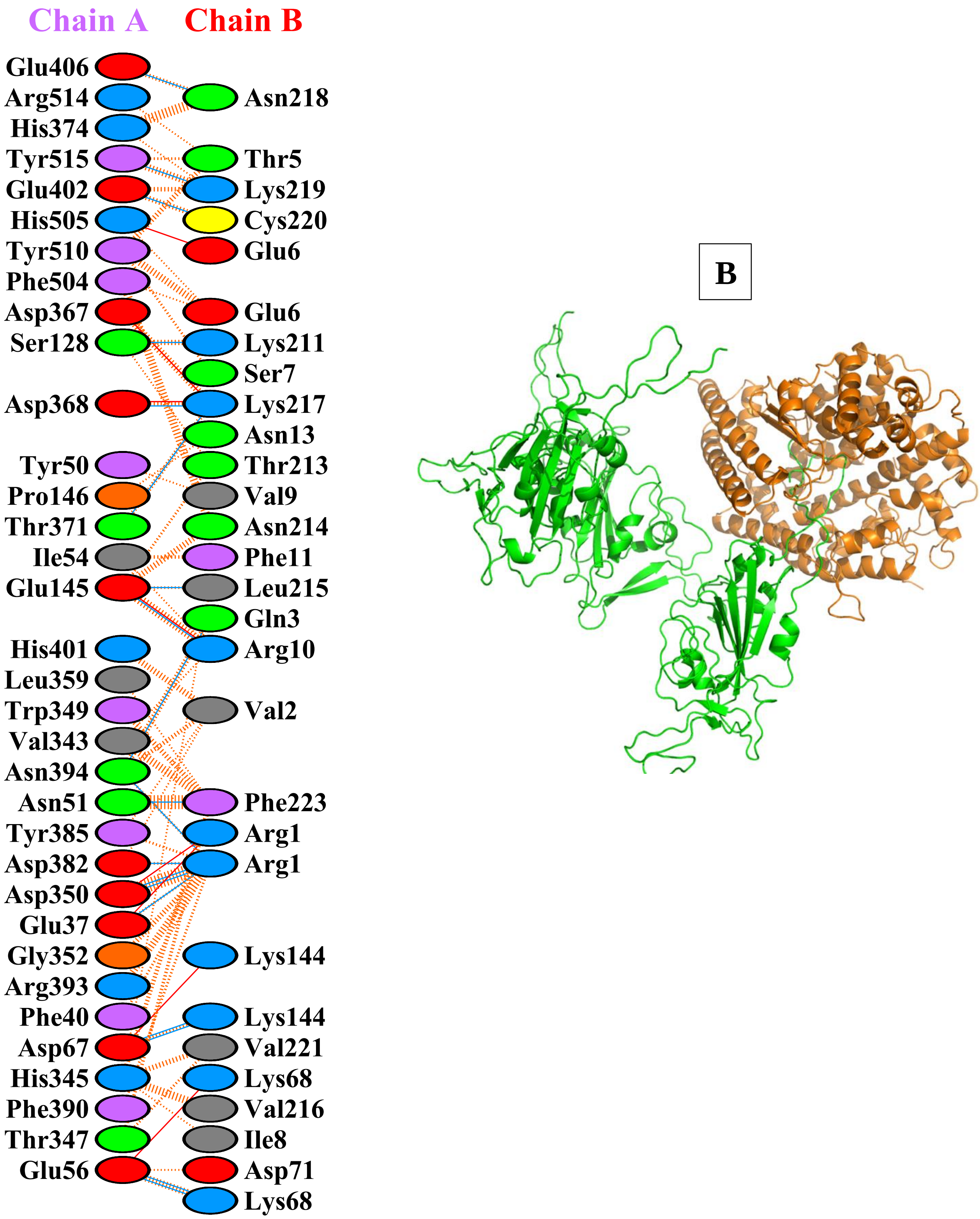


**Supplementary Figure:S2.D: Protein–protein docking representation of hACE2 and RBD of B.1.1.28.1 (Gamma) (EPI_ISL_811149).** (A) The interaction representation, which includes hydrogen ( ), salt bridges ( ), and non-bonded ( ) interactions. Chain A represents hACE2 and chain B represents RBD . Residue colors: positive (blue): H,K,R; negative (red): D,E; neutral (green): S,T,N,Q; aliphatic (grey): A,V,L,I,M; aromatic (purple): F,Y,W; Pro & Gly (orange): P,G. (B) The binding interface of hACE2:RBD complex.

A

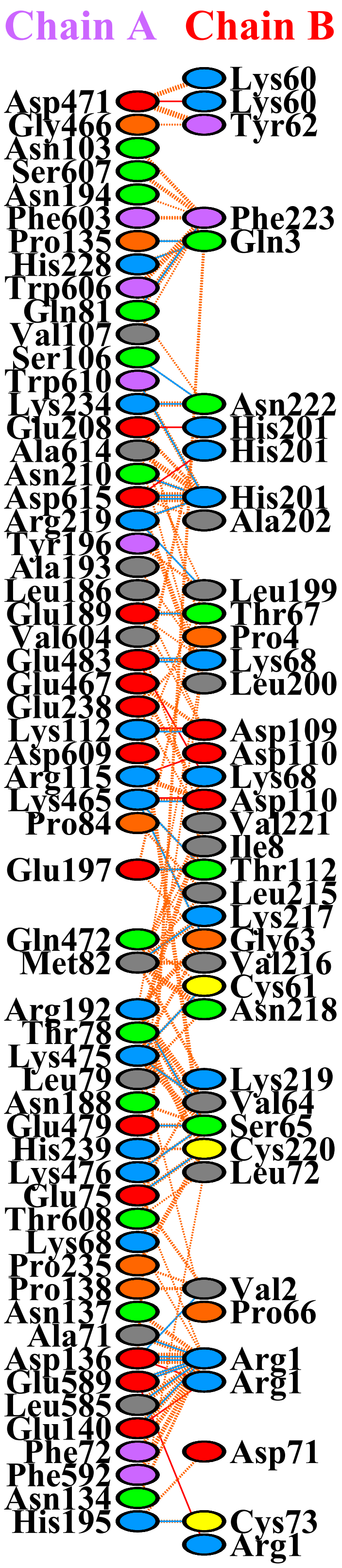


B

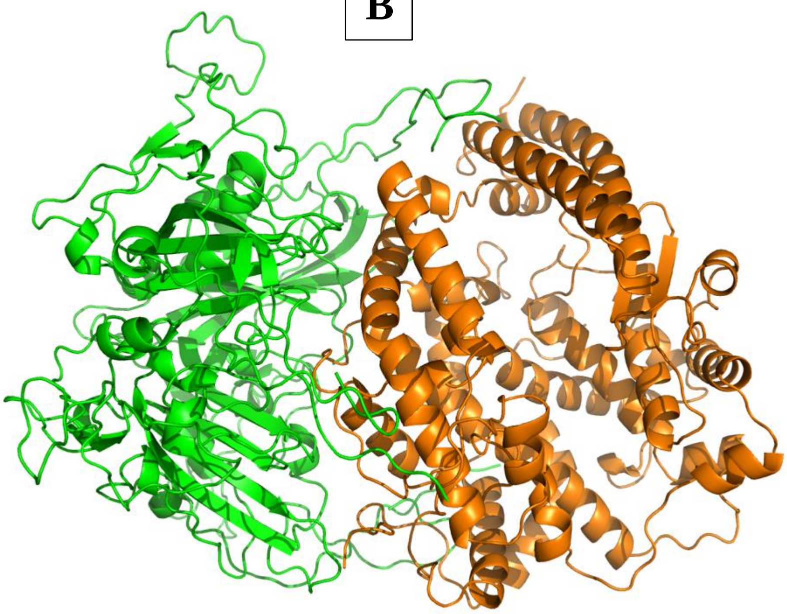

**Supplementary Figure:S2.E: Protein–protein docking representation of hACE2 and RBD of B.1.617(Delta) (EPI_ISL_1360328)**. (A) The interaction representation, which includes hydrogen ( ), salt bridges ( ), and non-bonded ( ) interactions. Chain A represents hACE2 and chain B represents RBD . Residue colors: positive (blue): H,K,R; negative (red): D,E; neutral (green): S,T,N,Q; aliphatic (grey): A,V,L,I,M; aromatic (purple): F,Y,W; Pro & Gly (orange): P,G. (B) The binding interface of hACE2:RBD complex.

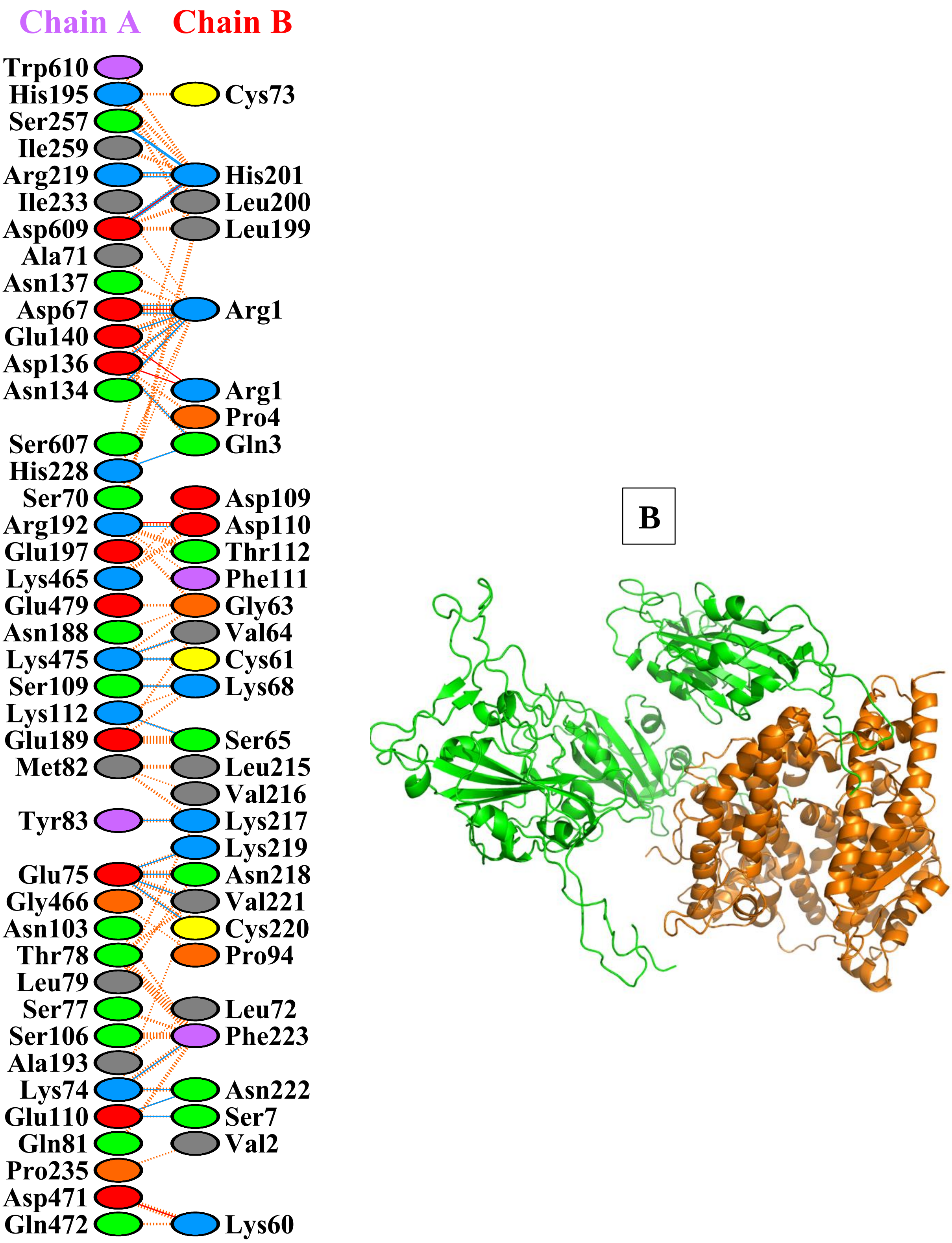


**Supplementary Figure:S2.F: Protein–protein docking representation of hACE2 and RBD of B.1.1.529 (Omicron) (EPI_ISL_6640917 ).** (A) The interaction representation, which includes hydrogen ( ), salt bridges ( ), and non-bonded ( A interactions. Chain A represents hACE2 and chain B represents RBD . Residue colors: positive (blue): H,K,R; negative (red): D,E; neutral (green): S,T,N,Q; aliphatic (grey): A,V,L,I,M; aromatic (purple): F,Y,W; Pro & Gly (orange): P,G. (B) The binding interface of hACE2:RBD complex.

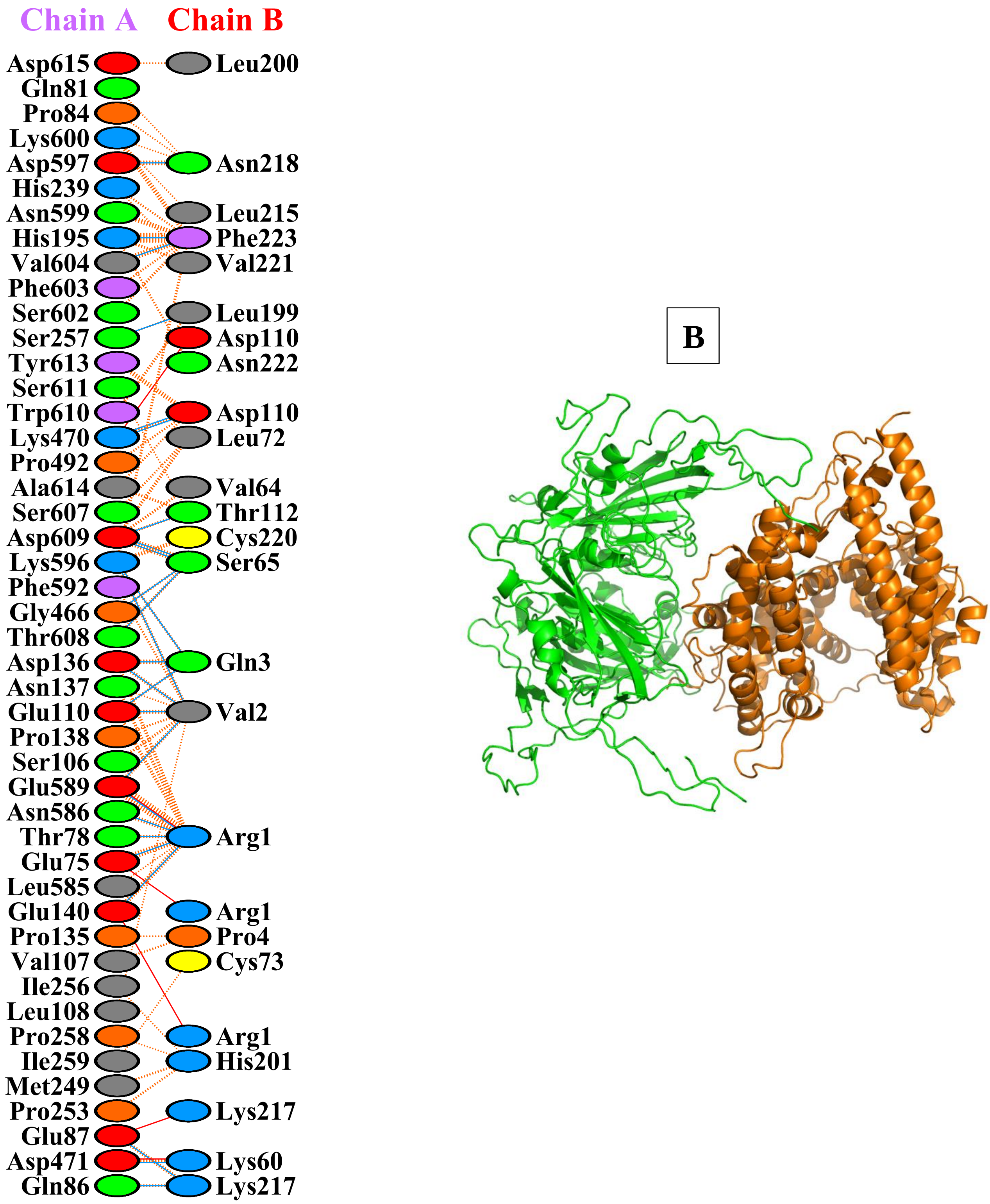


**Supplementary Figure:S2.G: Protein–protein docking representation of hACE2 and RBD of C.37(Lambda) (EPI_ISL_1111334).** (A) The interaction representation, which includes hydrogen ( ), salt bridges ( ), and non-bonded ( ) interactions. Chain A represents hACE2 and chain B represents RBD . Residue colors: positive (blue): H,K,R; negative ( **A** D,E; neutral (green): S,T,N,Q; aliphatic (grey): A,V,L,I,M; aromatic (purple): F,Y,W; Pro & Gly (orange): P,G. (B) The binding interface of hACE2:RBD complex.

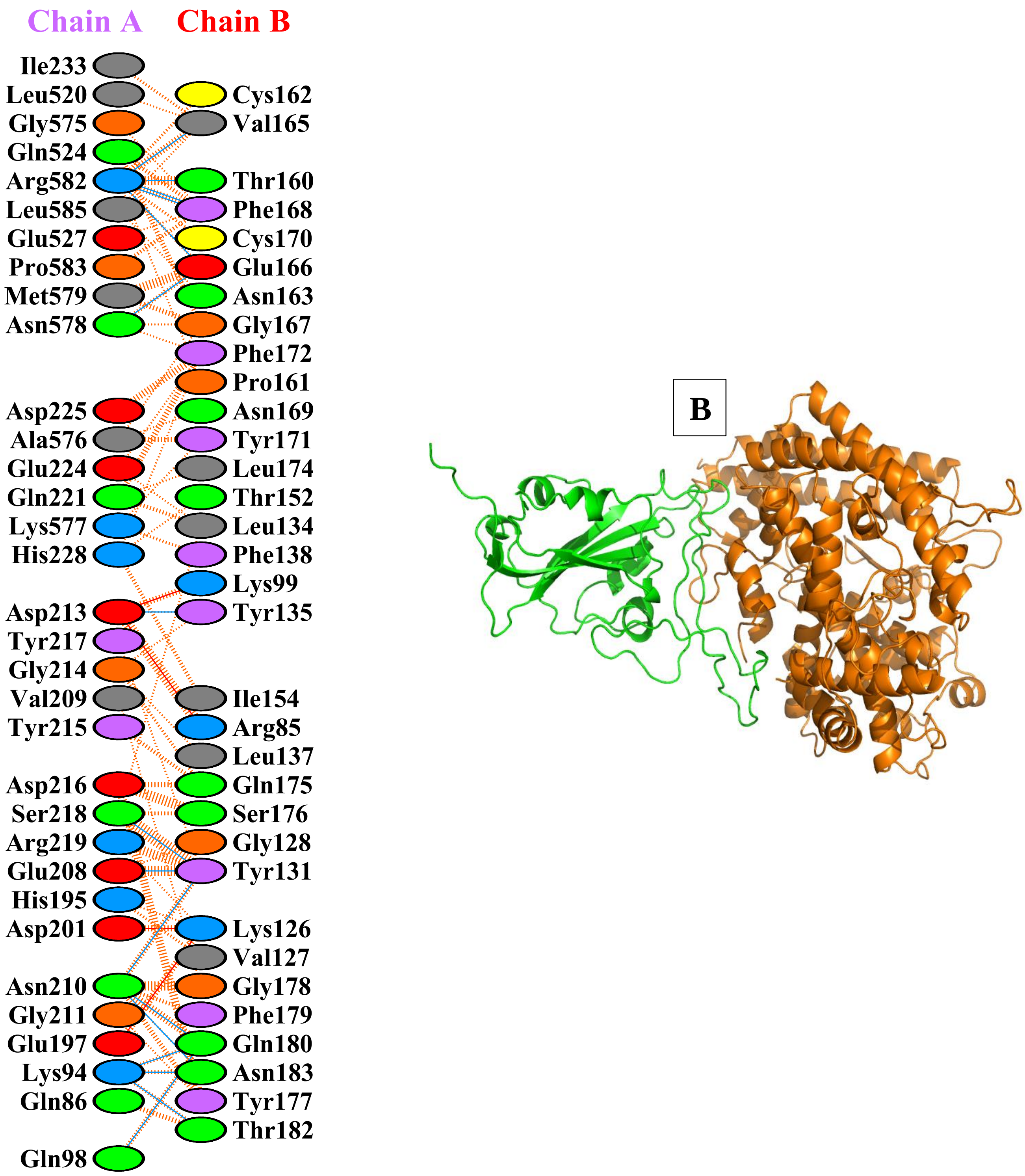


**Supplementary Figure:S2.H: Protein–protein docking representation of hACE2 and RBD of B.1.621(Mu) (EPI_ISL_3369952).** (A) The interaction representation, which includes hydrogen (▬), salt bridges (▬), and non-bonded (IIIII [illegible]ractions. Chain A represents hACE2 and chain B represents RBD . Residue colors: positive (blue): H,K,R[illegible]tive (red): D,E; neutral (green): S,T,N,Q; aliphatic (grey): A,V,L,I,M; aromatic (purple): F,Y,W; Pro & Gly (orange): P,G. (B) The binding interface of hACE2:RBD complex.

A

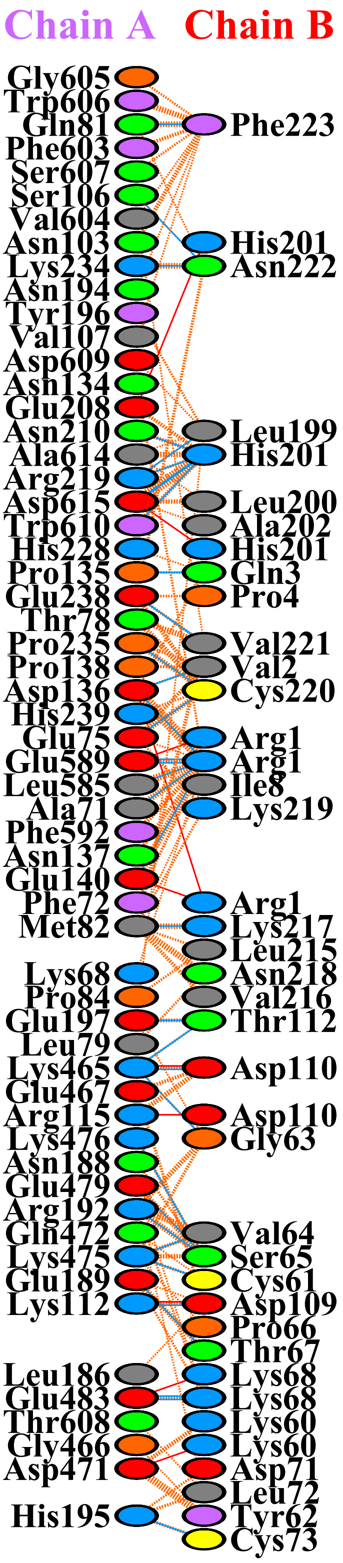


B

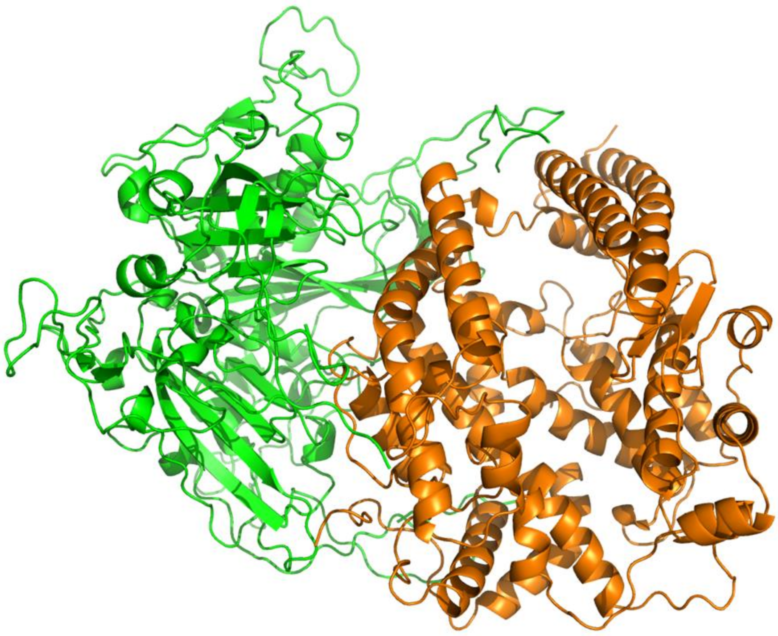

A

**Supplementary Figure:S2.I: Protein–protein docking representation of hACE2 and RBD of B.1.427/B.1.429 (Epsilon) (EPI_ISL_648527**). (A) The interaction representation, which includes hydrogen ( ), salt bridges ( ), and non-bonded [illegible] ) interactions. Chain A represents hACE2 and chain B represents RBD . Residue colors: positive (blue): H,K,R; negative (red): D,E; neutral (green): S,T,N,Q; aliphatic (grey): A,V,L,I,M; aromatic (purple): F,Y,W; Pro & Gly (orange): P,G. (B) The binding interface of hACE2:RBD complex.

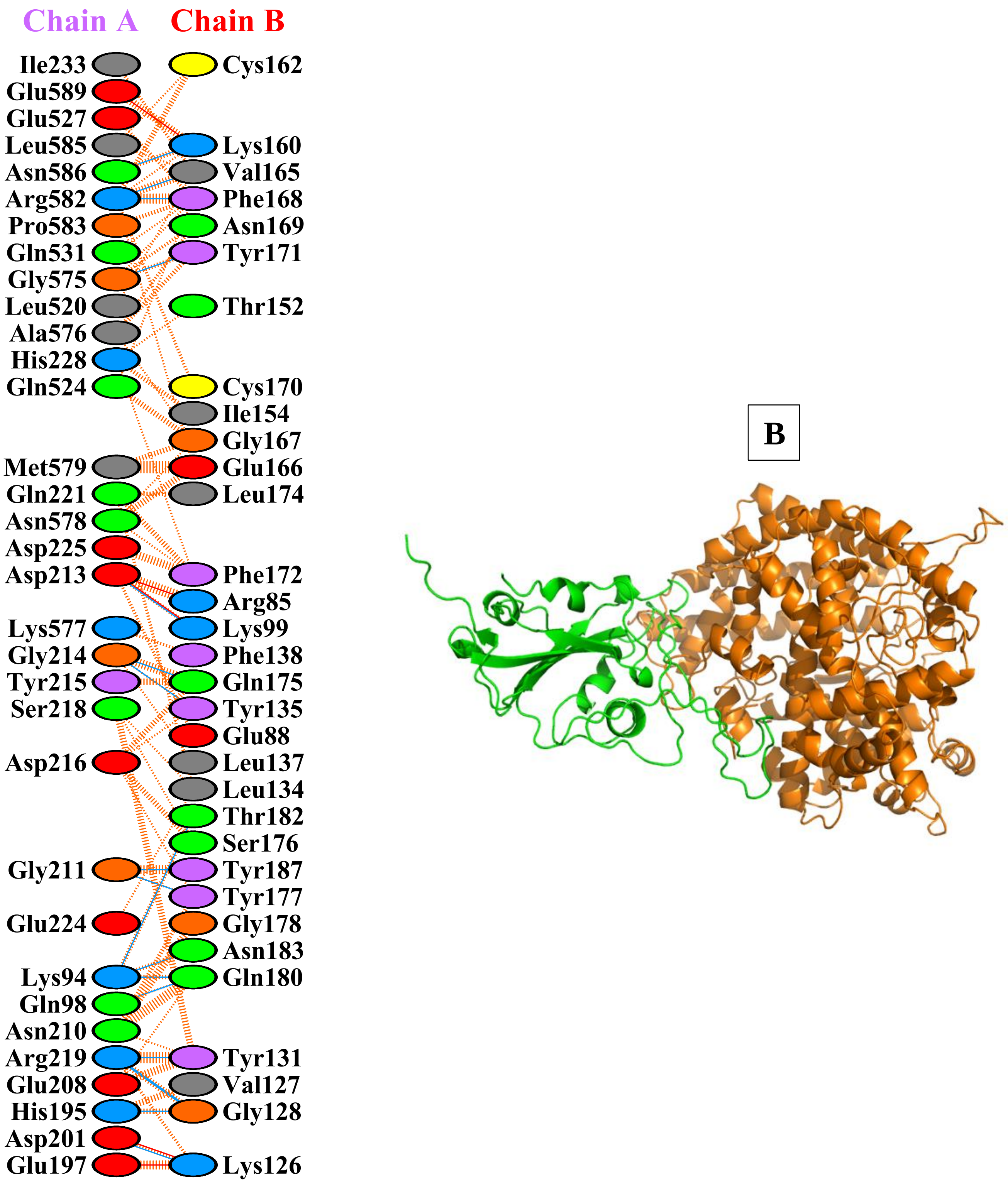


**Supplementary Figure:S2.J: Protein–protein docking representation of hACE2 and RBD of B.1.1.519 (EPI_ISL_721617).** (A) The interaction representation, which includes hydrogen ( ), salt bridges ( ), and non-bonded ( ) interactions. Chain A represents hACE2 and chain B represents RBD . Residue colors: positive (blue): H,K,R; negative ( A D,E; neutral (green): S,T,N,Q; aliphatic (grey): A,V,L,I,M; aromatic (purple): F,Y,W; Pro & Gly (orange): P,G. (B) The binding interface of hACE2:RBD complex.

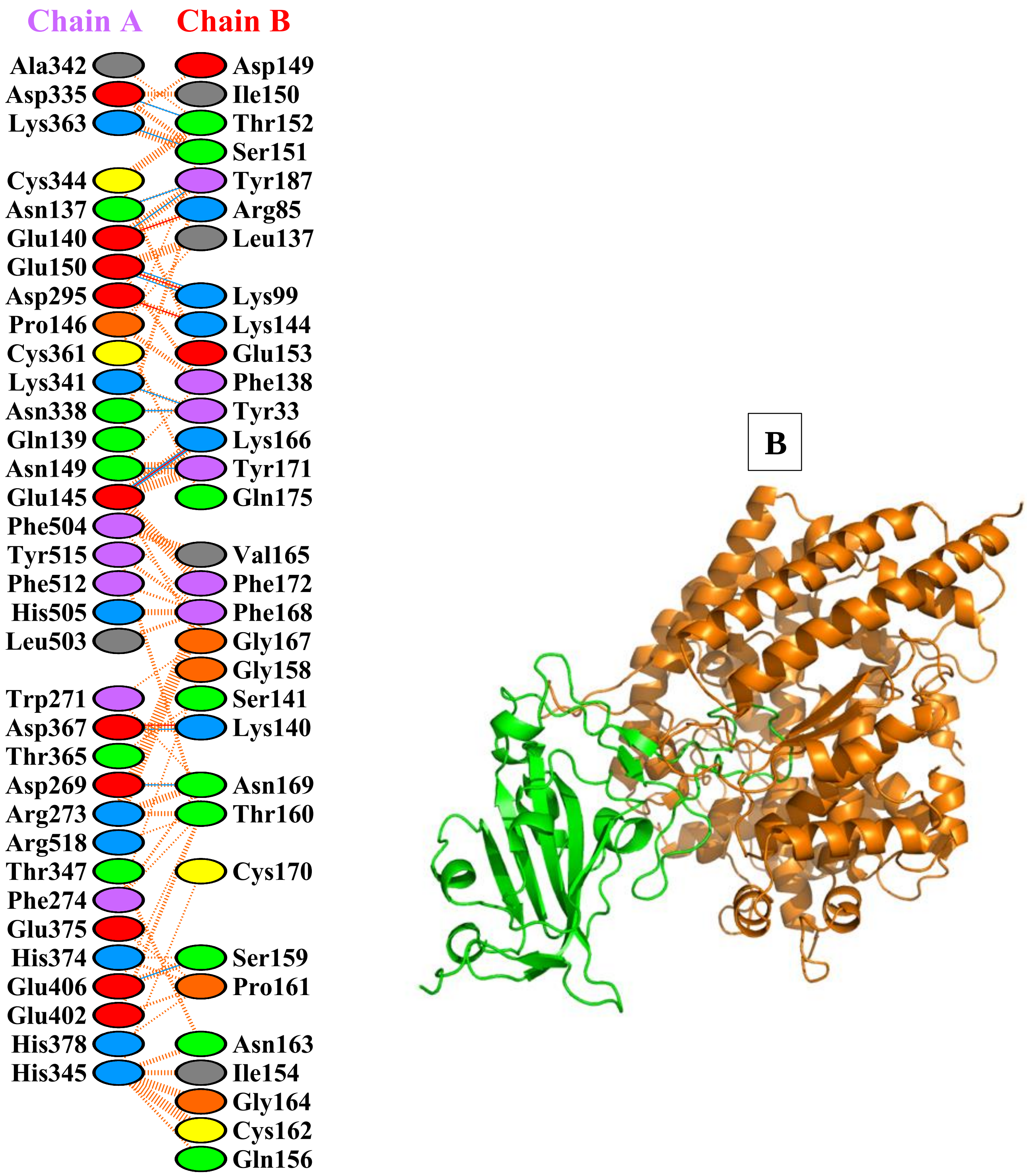


**Supplementary Figure:S2.K: Protein–protein docking representation of hACE2 and RBD of R.1 (EPI_ISL_736897).** (A) The interaction representation, which includes hydrogen (▬▬), salt bridges (▬▬), and non-bonded (IIIIII) inter[illegible]s. Chain A represents hACE2 and chain B represents RBD . Residue colors: positive (blue): H,K,R; negative (red): D,E; neutral (green): S,T,N,Q; aliphatic (grey): A,V,L,I,M; aromatic (purple): F,Y,W; Pro & Gly (orange): P,G. (B) The binding interface of hACE2:RBD complex.

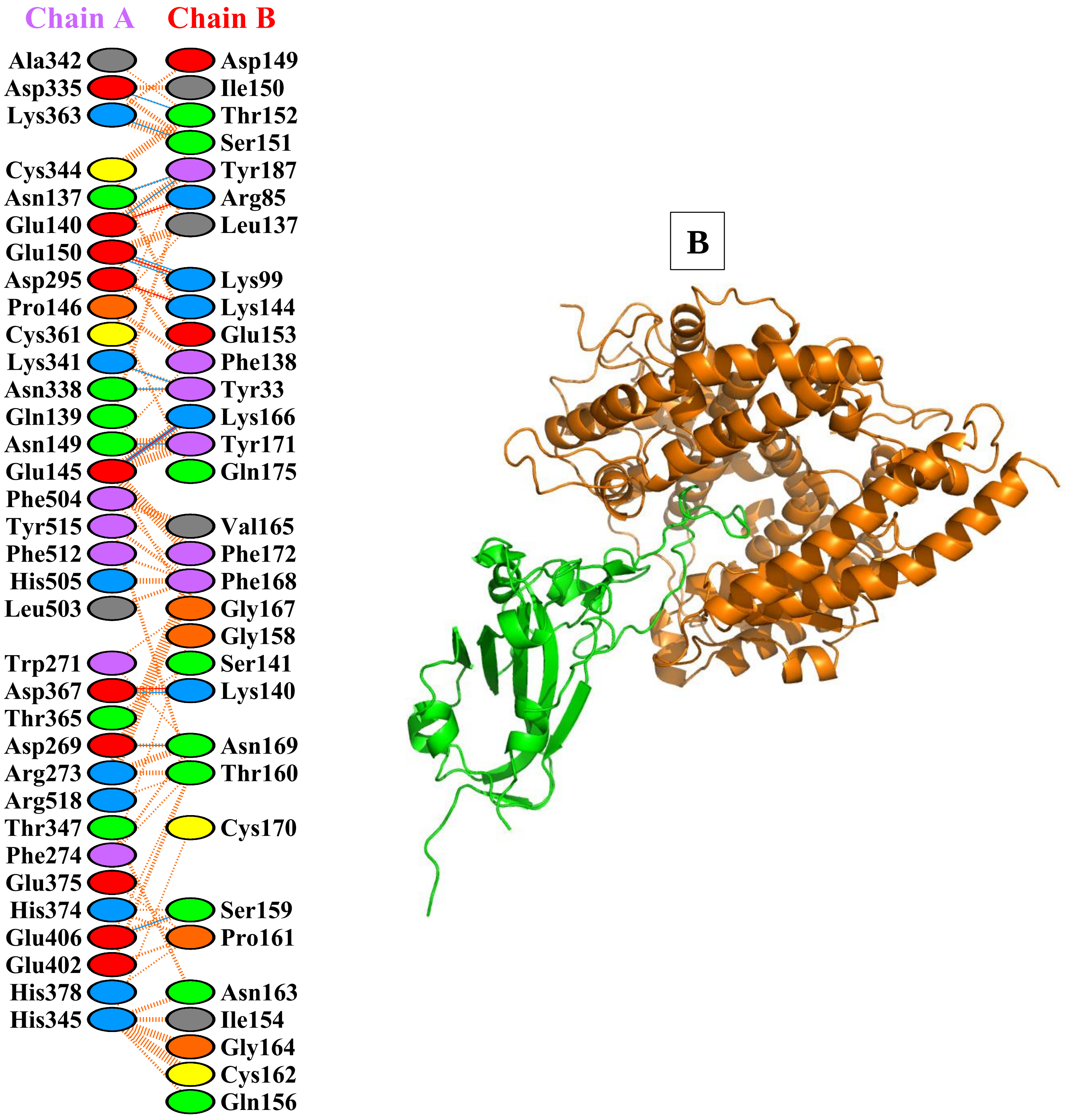


**Supplementary Figure:S2.L: Protein–protein docking representation of hACE2 and RBD of B.1.525(Eta) (EPI_ISL_760883).** (A) The interaction representation, which includes hydrogen ( ), salt bridges ( ), and non-bonded ( ) intera[illegible] Chain A represents hACE2 and chain B represents RBD . Residue colors: positive (blue): H,K,R; negative (re[illegible] A ,E; neutral (green): S,T,N,Q; aliphatic (grey): A,V,L,I,M; aromatic (purple): F,Y,W; Pro & Gly (orange): P,G. (B) The binding interface of hACE2:RBD complex.

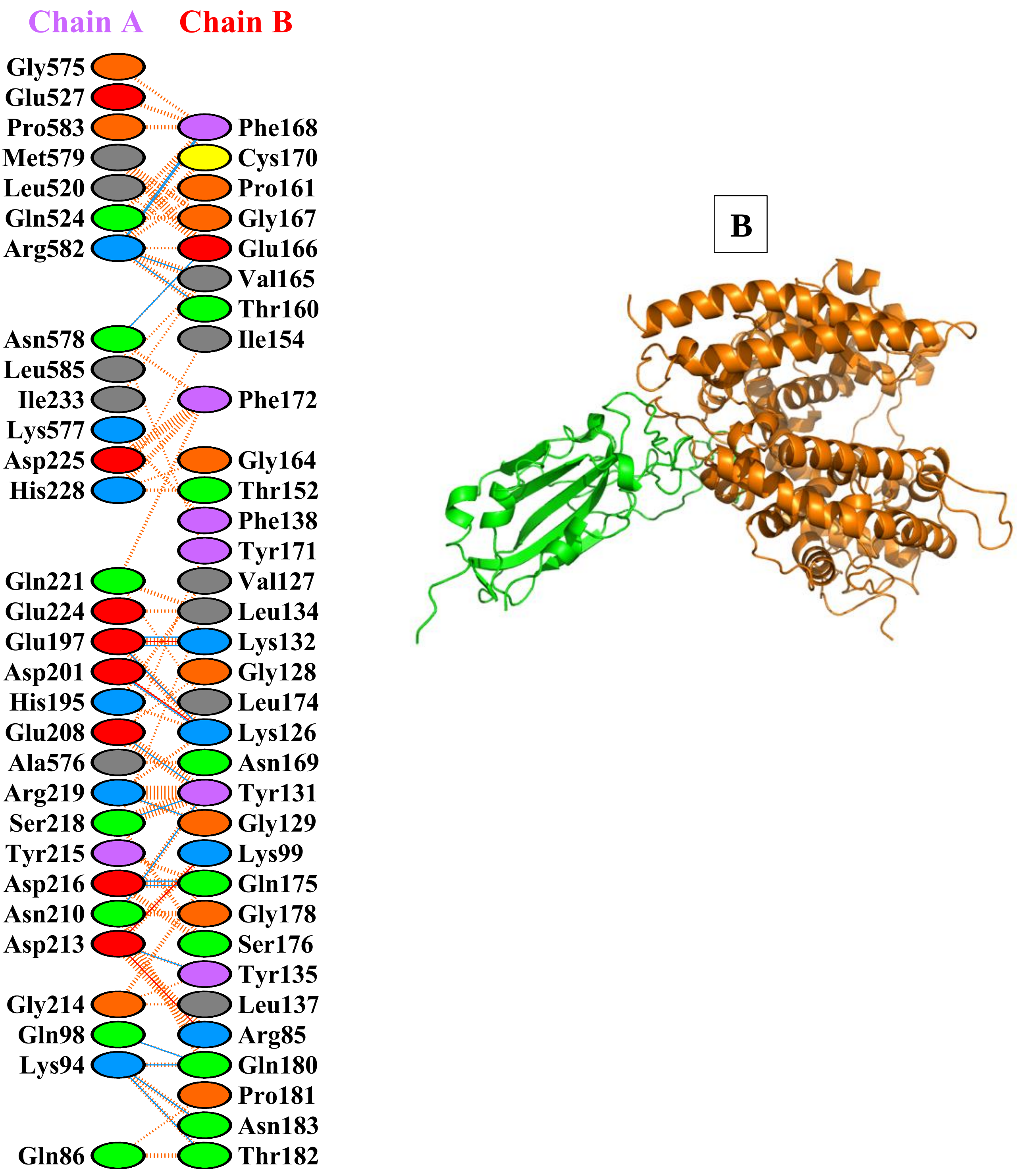


**Supplementary Figure:S2.M: Protein–protein docking representation of hACE2 and RBD of B.1.214.2 (EPI_ISL_760951).** (A) The interaction representation, which includes hydrogen ( ), salt bridges ( ), and non-bonded (IIIIII) interact[illegible] Chain A represents hACE2 and chain B represents RBD . Residue colors: positive (blue): H,K,R; negative (rec[illegible]; neutral (green): S,T,N,Q; aliphatic (grey): A,V,L,I,M; aromatic (purple): F,Y,W; Pro & Gly (orange): P,G. (B) The binding interface of hACE2:RBD complex.

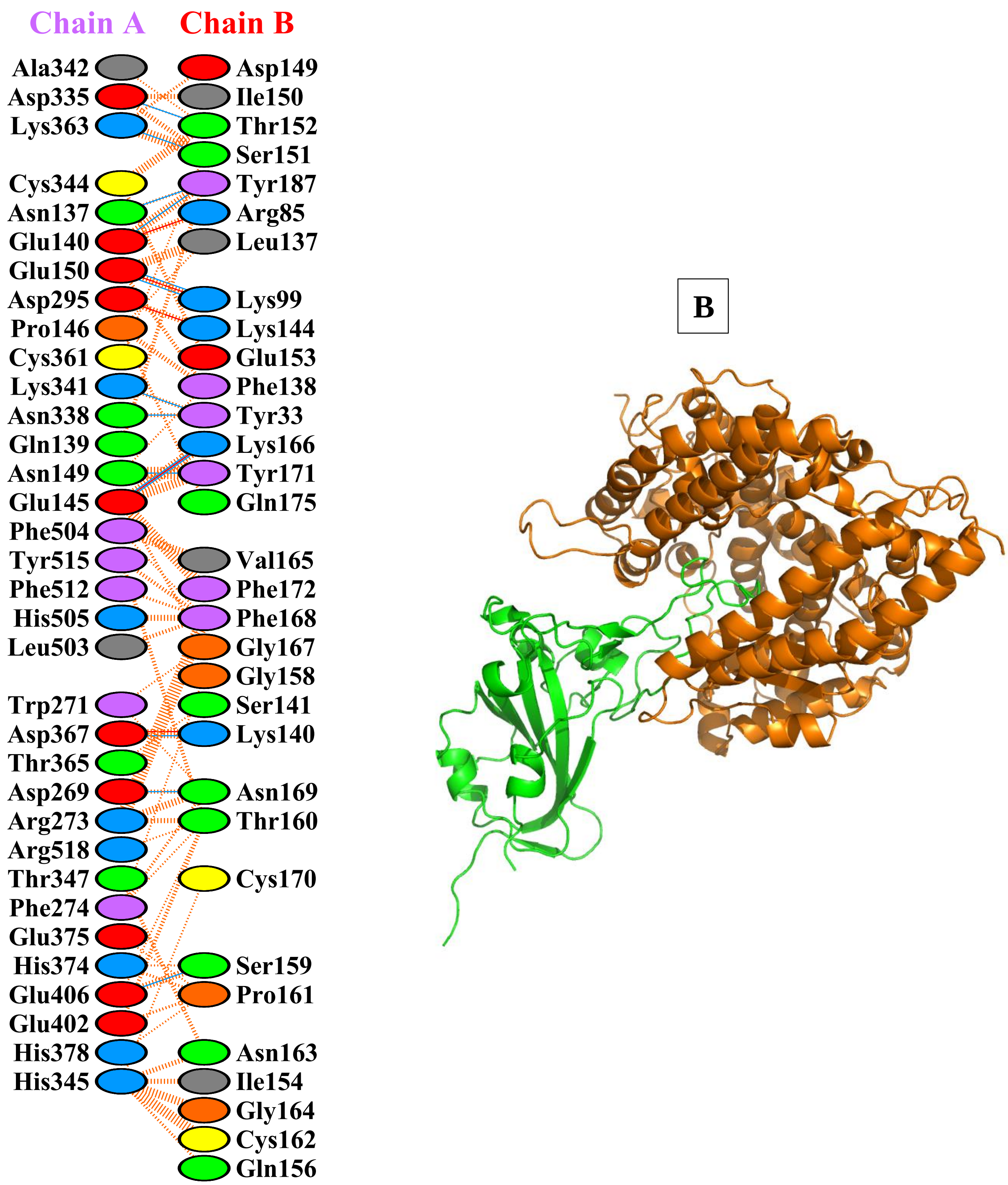


**Supplementary Figure:S2.N: Protein–protein docking representation of hACE2 and RBD of B.1.526(Lota) (EPI_ISL_765494).** (A) The interaction representation, which includes hydrogen ( ), salt bridges ( ), and non-bonded ( ) intera[illegible] **A** [illegible]s. Chain A represents hACE2 and chain B represents RBD . Residue colors: positive (blue): H,K,R; negative (red): D,E; neutral (green): S,T,N,Q; aliphatic (grey): A,V,L,I,M; aromatic (purple): F,Y,W; Pro & Gly (orange): P,G. (B) The binding interface of hACE2:RBD complex.

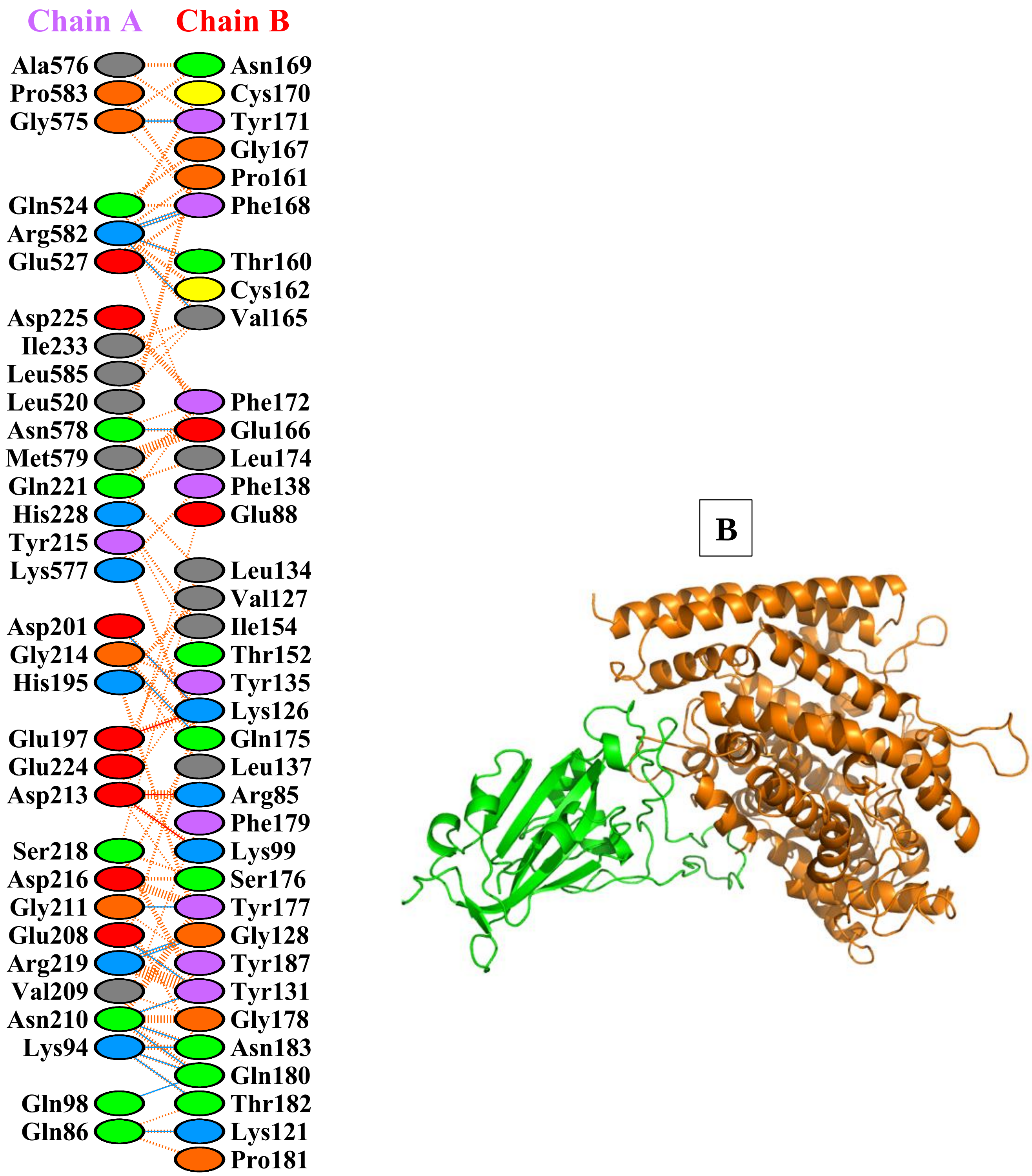


**Supplementary Figure:S2.O: Protein–protein docking representation of hACE2 and RBD of B.1.466.2 (EPI_ISL_877419).** (A) The interaction representation, which includes hydrogen ( ), salt bridges ( ), and non-bonded ( A interactions. Chain A represents hACE2 and chain B represents RBD . Residue colors: positive (blue): H,K,R; negative (red): D,E; neutral (green): S,T,N,Q; aliphatic (grey): A,V,L,I,M; aromatic (purple): F,Y,W; Pro & Gly (orange): P,G. (B) The binding interface of hACE2:RBD complex.

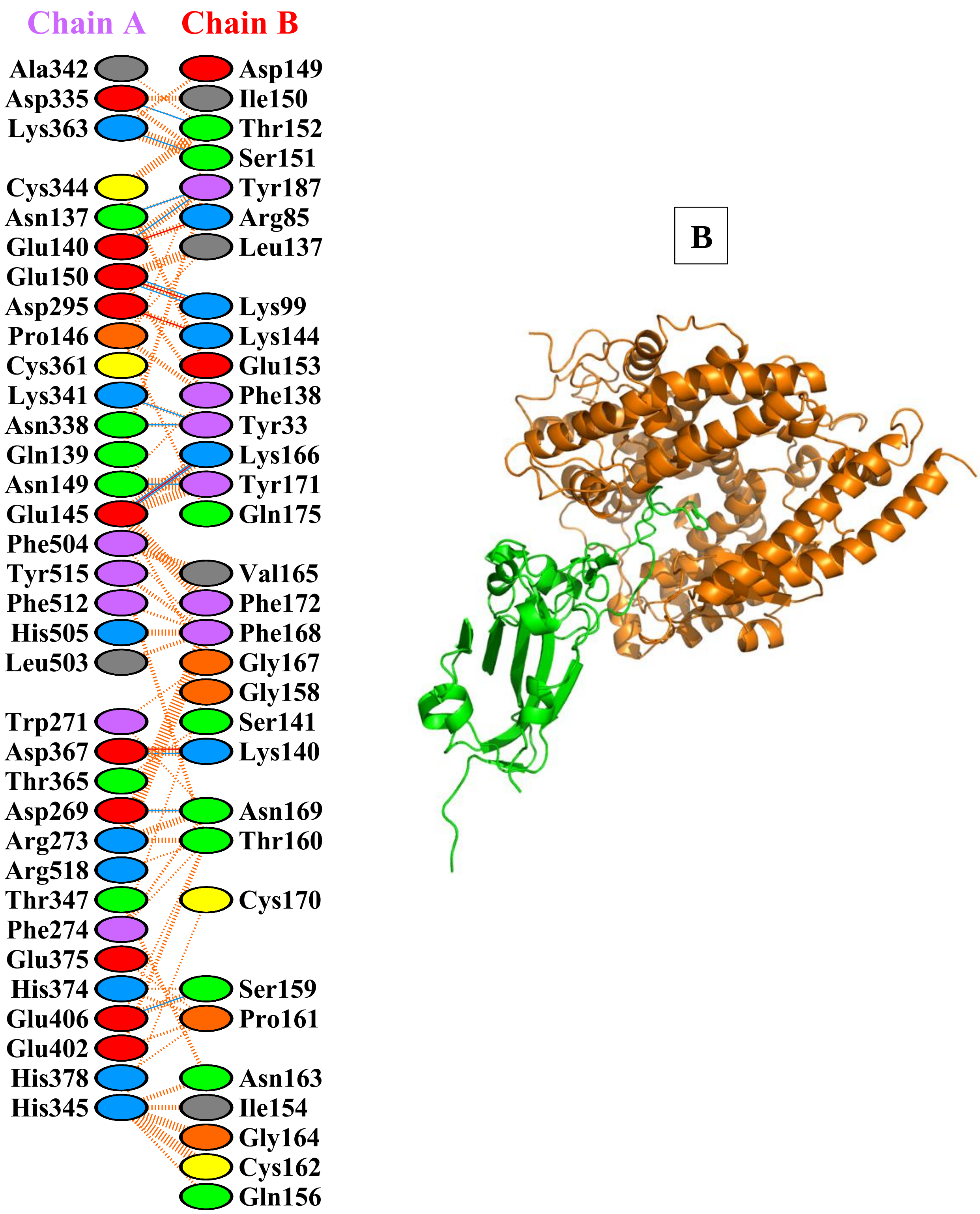


**Supplementary Figure:S2.P: Protein–protein docking representation of hACE2 and RBD of B.1.1.318 (EPI_ISL_937654)**. (A) The interaction representation, which includes hydrogen (▬▬), salt bridges (▬▬), and non-bonded (IIIIIII) interacti[illegible] A [illegible]hain A represents hACE2 and chain B represents RBD . Residue colors: positive (blue): H,K,R; negative (red): D,E; neutral (green): S,T,N,Q; aliphatic (grey): A,V,L,I,M; aromatic (purple): F,Y,W; Pro & Gly (orange): P,G. (B) The binding interface of hACE2:RBD complex.

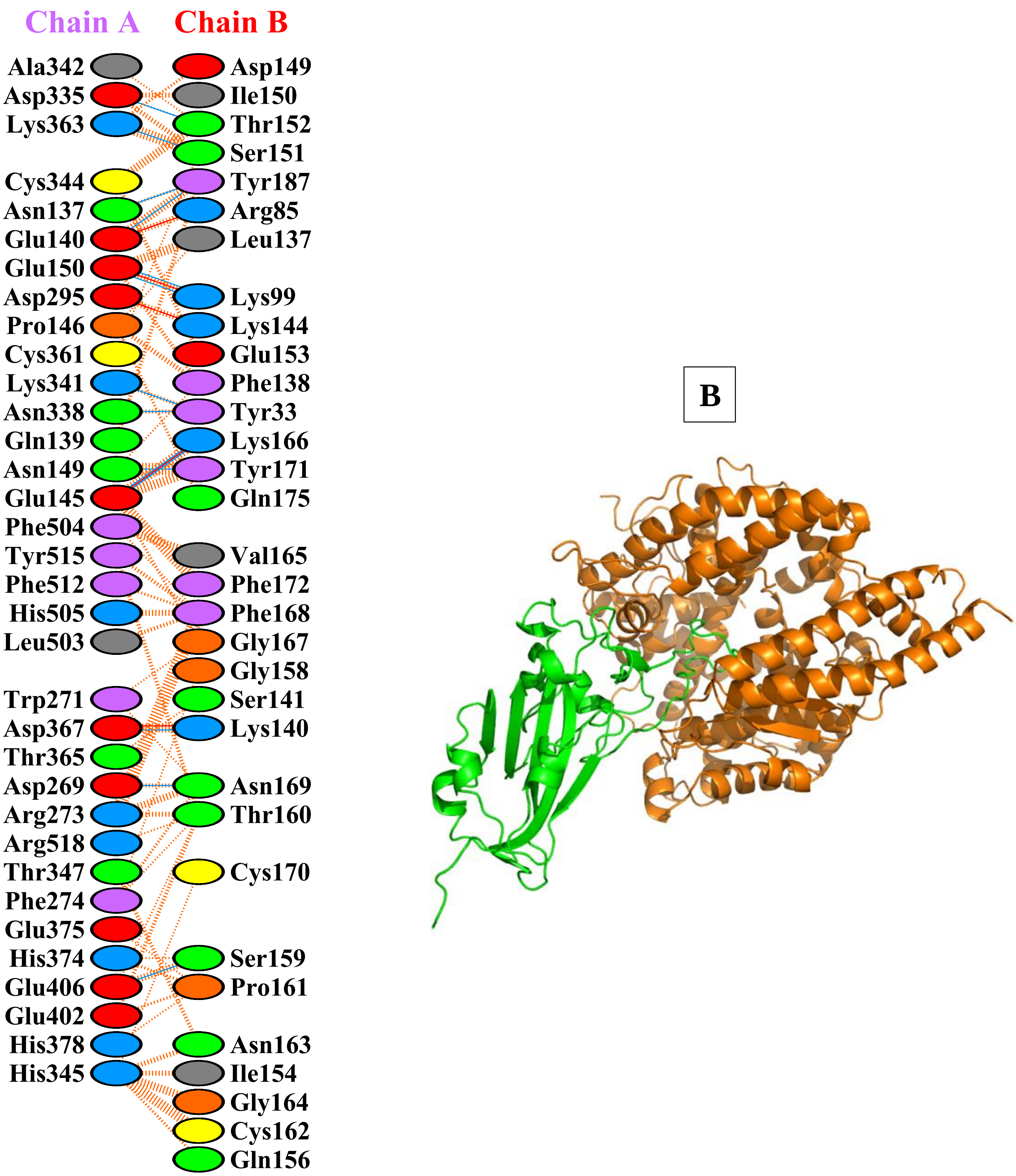


**Supplementary Figure:S2.Q: Protein–protein docking representation of hACE2 and RBD of B.1.619 (EPI_ISL_1150929 ).** (A [illegible] interaction representation, which includes hydrogen (—), salt bridges (—), and non-bonded (IIIIII) [illegible]tions. Chain A represents hACE2 and chain B represents RBD . Residue colors: positive (blue): H,K,R; negative (red): D,E; neutral (green): S,T,N,Q; aliphatic (grey): A,V,L,I,M; aromatic (purple): F,Y,W; Pro & Gly (orange): P,G. (B) The binding interface of hACE2:RBD complex.

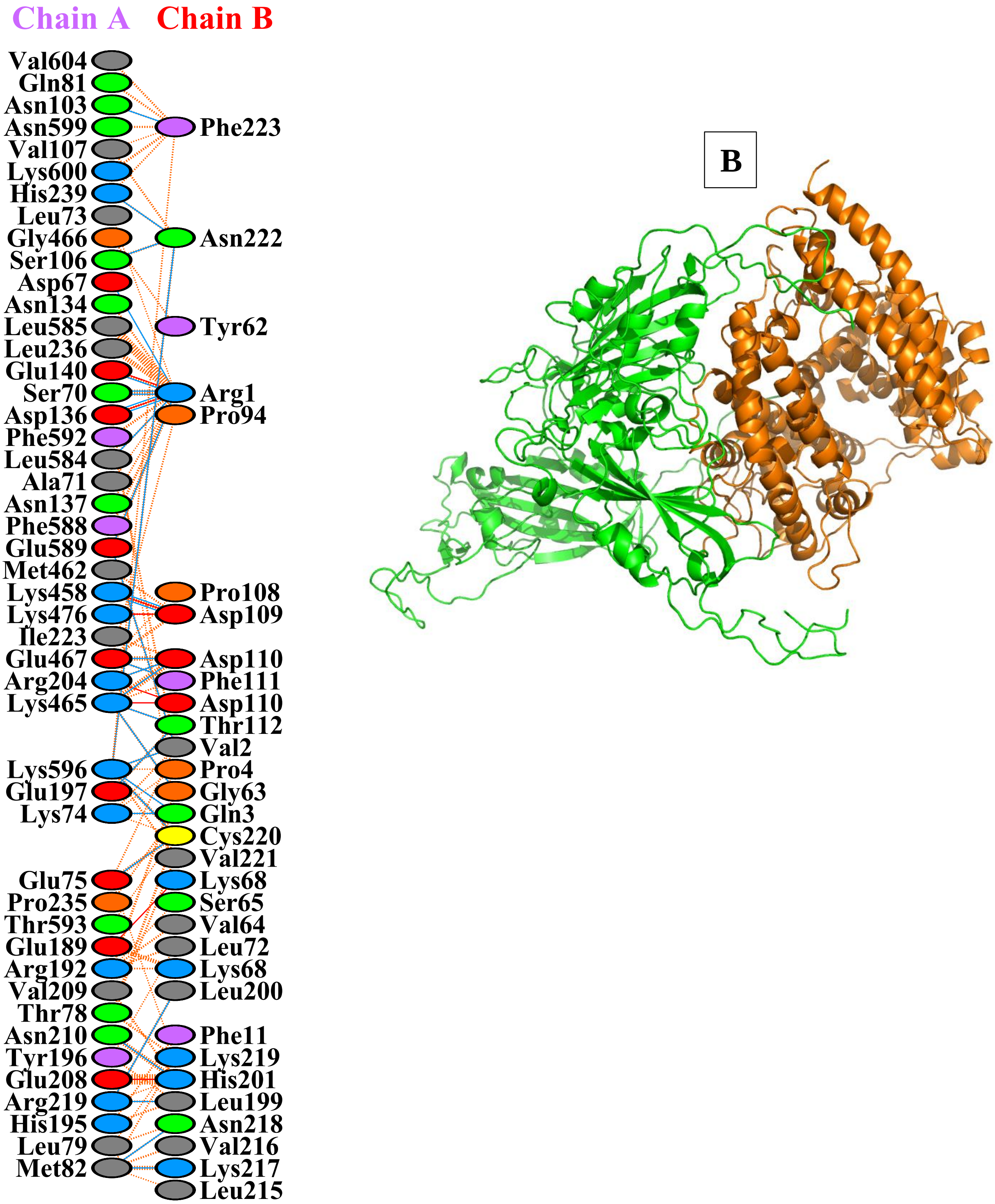


**Supplementary Figure:S2.R: Protein–protein docking representation of hACE2 and RBD of C.36.3 (EPI_ISL_1237137).** (A) The interaction representation, which includes hydrogen ( ), salt bridges ( ), and non-bonded ( ) inte[illegible]ns. Chain A represents hACE2 and chain B represents RBD . Residue colors: positive (blue): H,K,R; negative (red): D,E; neutral (green): S,T,N,Q; aliphatic (grey): A,V,L,I,M; aromatic (purple): F,Y,W; Pro & Gly (orange): P,G. (B) The binding interface of hACE2:RBD complex.

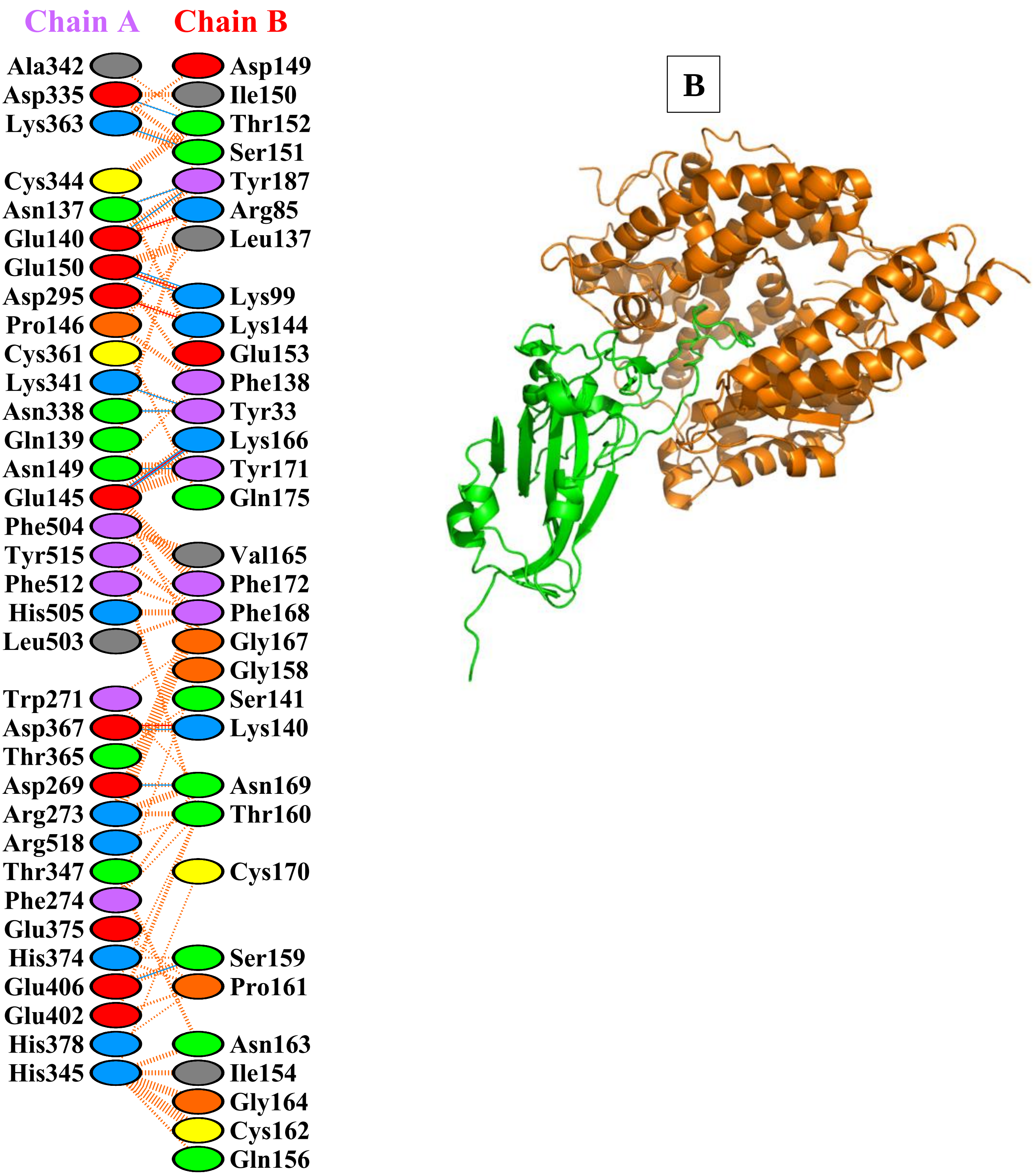


**Supplementary Figure:S2.S: Protein–protein docking representation of hACE2 and RBD of AT.1 (EPI_ISL_1259283) .** (A) The interaction representation, which includes hydrogen ( ), salt bridges ( ), and non-bonded ( [illegible] A [illegible]teractions. Chain A represents hACE2 and chain B represents RBD . Residue colors: positive (blue): H,K,[illegible]gative (red): D,E; neutral (green): S,T,N,Q; aliphatic (grey): A,V,L,I,M; aromatic (purple): F,Y,W; Pro & Gly (orange): P,G. (B) The binding interface of hACE2:RBD complex.

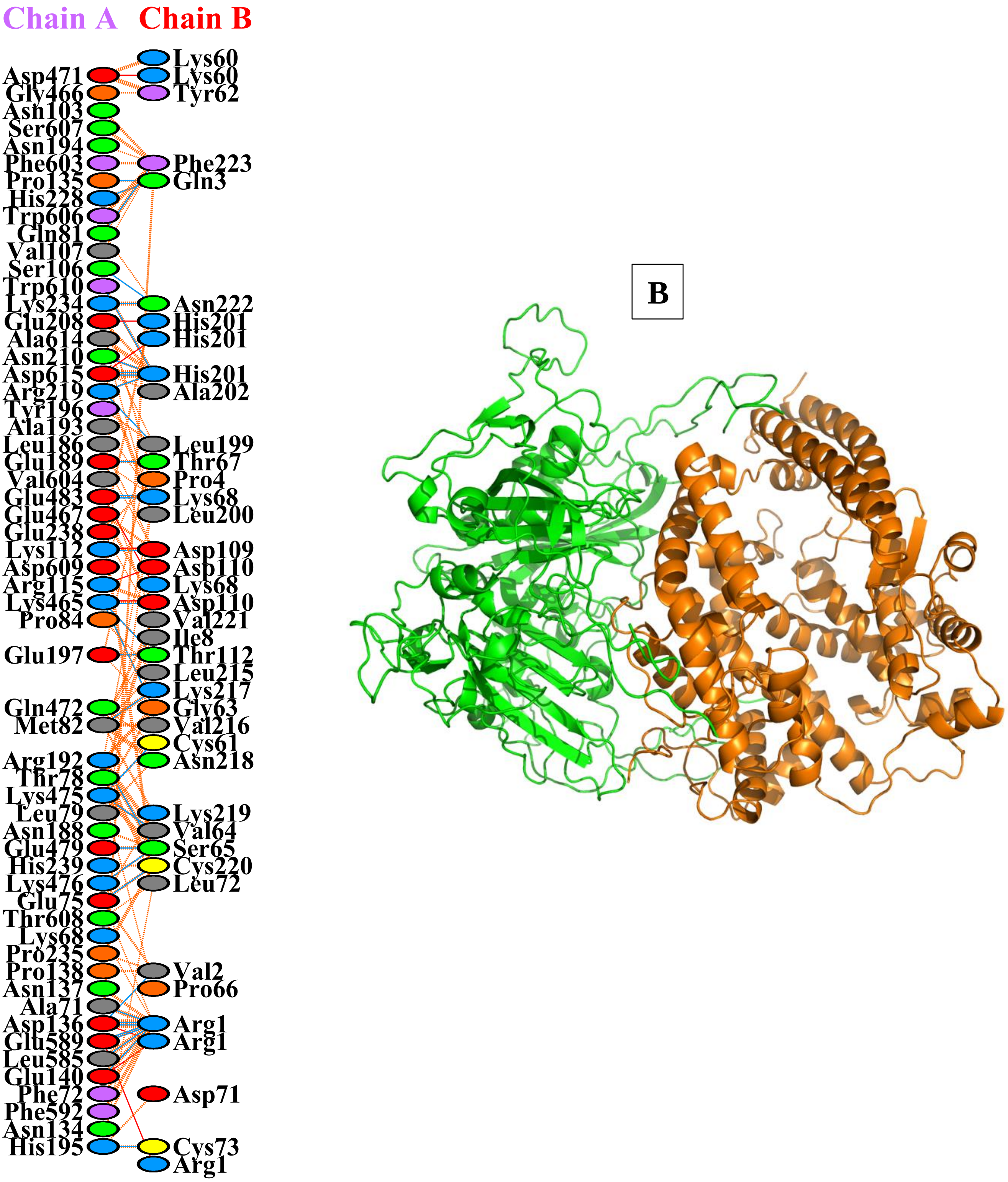


**Supplementary Figure:S2.T: Protein–protein docking representation of hACE2 and RBD of B.1.617.1(Kappa) (EPI_ISL_1357699)**. ([illegible] A [illegible]e interaction representation, which includes hydrogen (blue line), salt bridges (red line), and non-bonded (orange dashes) inte[illegible]ns. Chain A represents hACE2 and chain B represents RBD . Residue colors: positive (blue): H,K,R; negative (red): D,E; neutral (green): S,T,N,Q; aliphatic (grey): A,V,L,I,M; aromatic (purple): F,Y,W; Pro & Gly (orange): P,G. (B) The binding interface of hACE2:RBD complex.

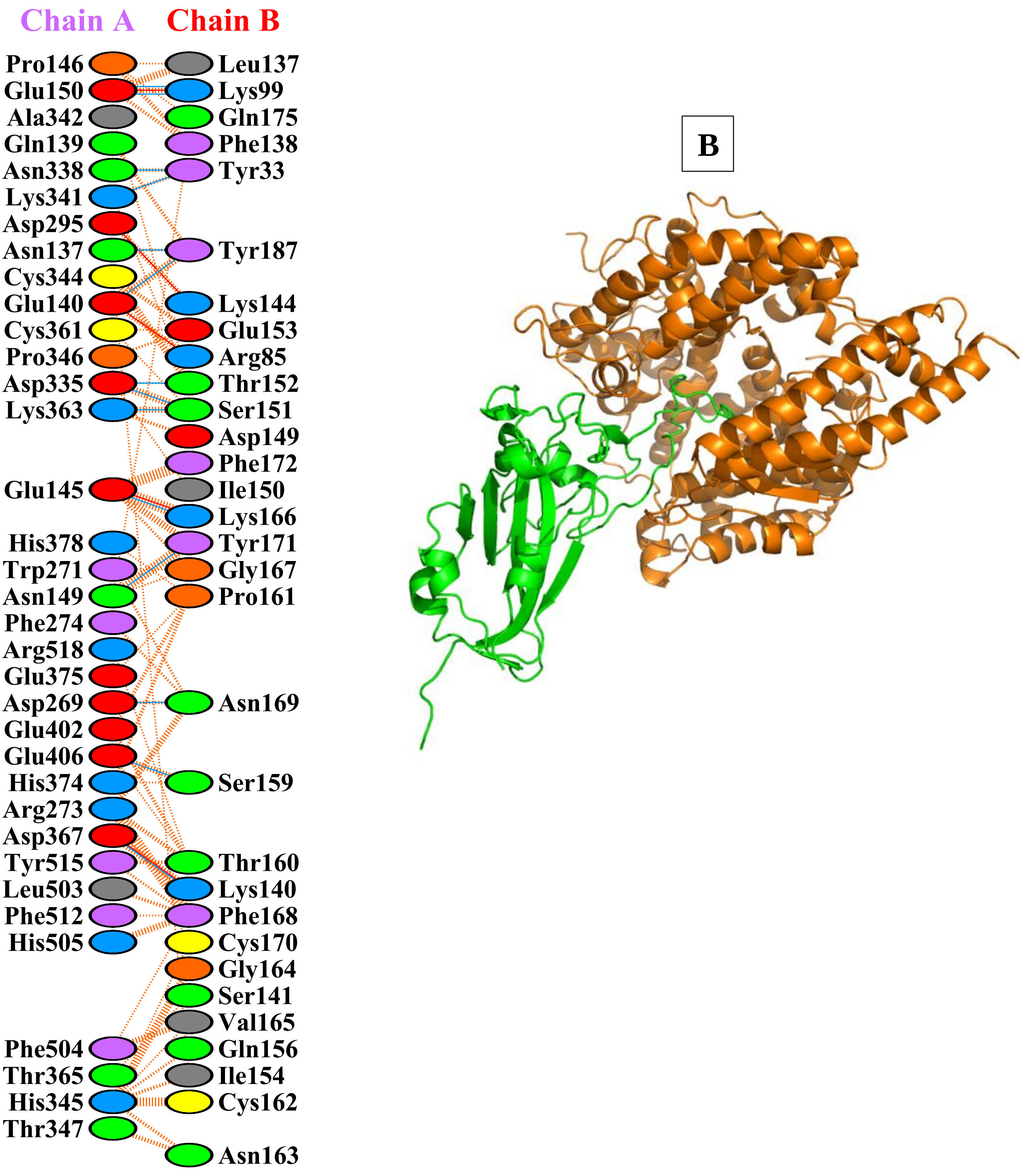


**Supplementary Figure:S2.U: Protein–protein docking representation of hACE2 and RBD of B.1.1.523 (EPI_ISL_1448584)** . (A) The interaction representation, which includes hydrogen (▬▬), salt bridges (▬▬), and non-bonded (IIIIIII) inte[illegible]ns. Chain A represents hACE2 and chain B represents RBD . Residue colors: positive (blue): H,K,R; negative (red): D,E; neutral (green): S,T,N,Q; aliphatic (grey): A,V,L,I,M; aromatic (purple): F,Y,W; Pro & Gly (orange): P,G. (B) The binding interface of hACE2:RBD complex.

A

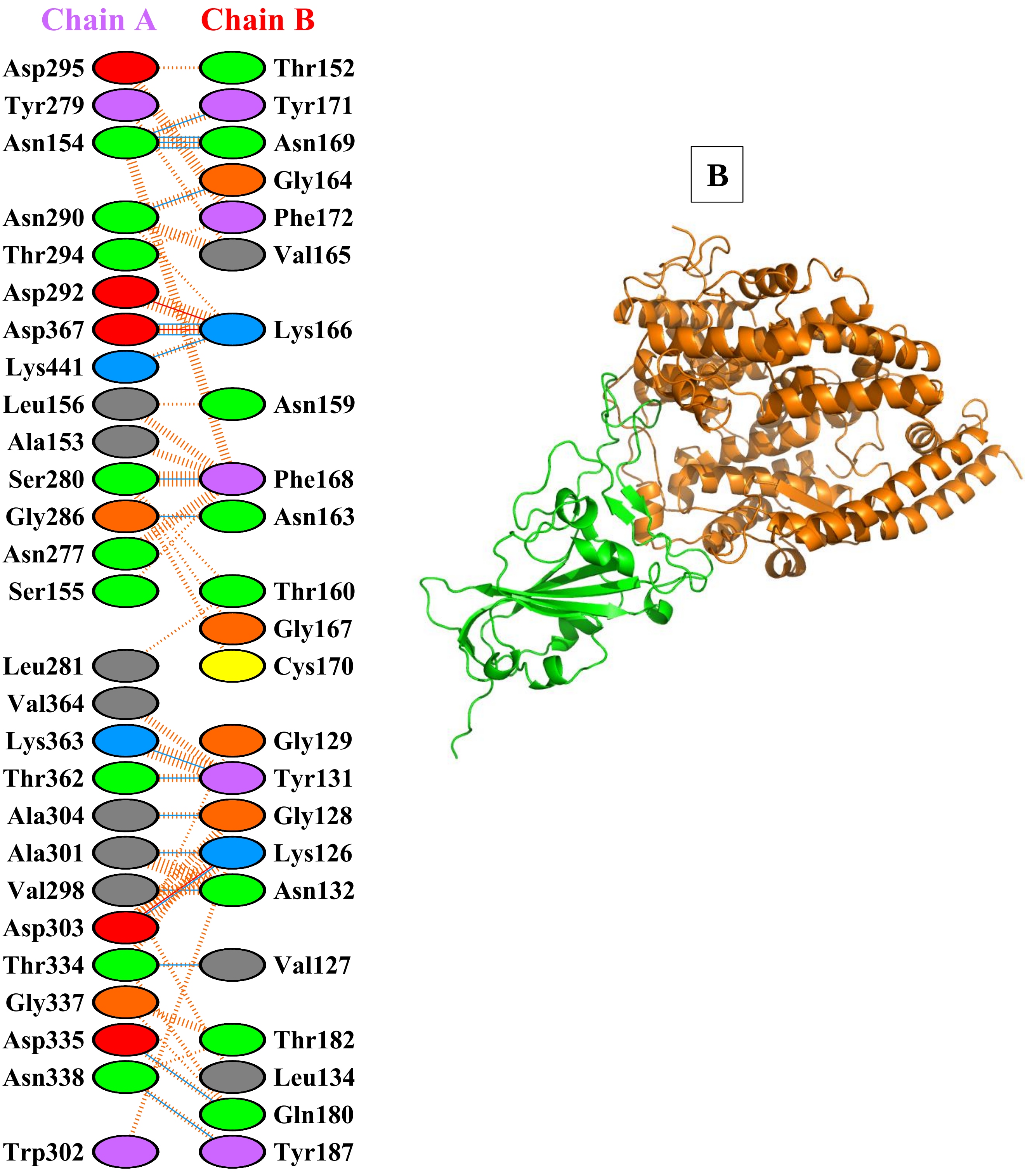


**Supplementary Figure:S2.V: Protein–protein docking representation of hACE2 and RBD of B.1.620 (EPI_ISL_1579527)**. (A) The interaction representation, which includes hydrogen ( ), salt bridges ( ), and non-bonded ( ) interactions. Chain A represents hACE2 and chain B represents RBD . Residue colors: positive (blue): H,K,R; ne[illegible] **A** (red): D,E; neutral (green): S,T,N,Q; aliphatic (grey): A,V,L,I,M; aromatic (purple): F,Y,W; Pro & Gl[illegible]nge): P,G. (B) The binding interface of hACE2:RBD complex.

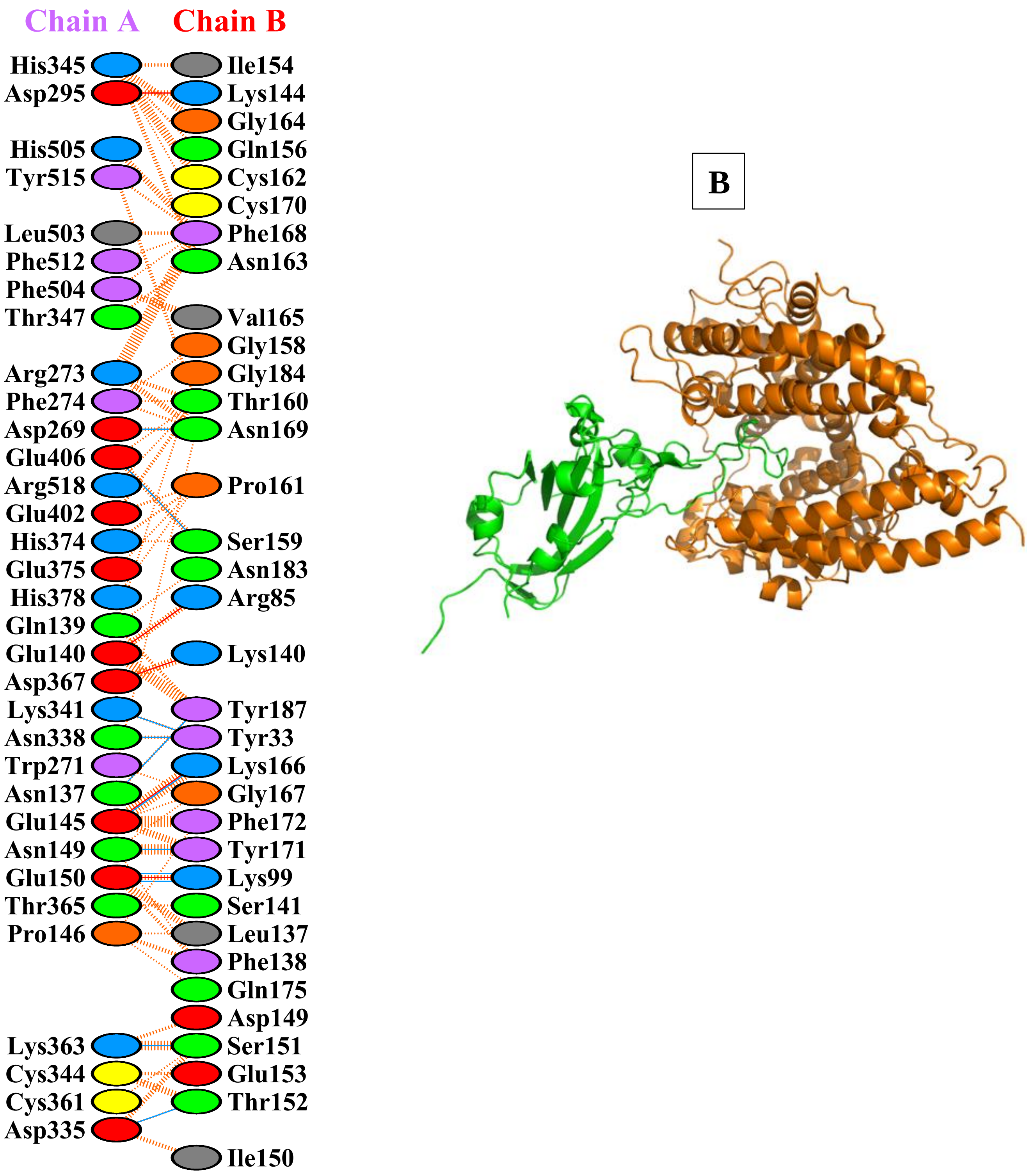


**Supplementary Figure:S2.W: Protein–protein docking representation of hACE2 and RBD of AV.1 (EPI_ISL_1595332).** (A) The interaction representation, which includes hydrogen ( ), salt bridges ( ), and non-bonded ( ) [illegible]teractions. Chain A represents hACE2 and chain B represents RBD . Residue colors: positive (blue): H,K,[illegible]gative (red): D,E; neutral (green): S,T,N,Q; aliphatic (grey): A,V,L,I,M; aromatic (purple): F,Y,W; Pro & Gly (orange): P,G. (B) The binding interface of hACE2:RBD complex.

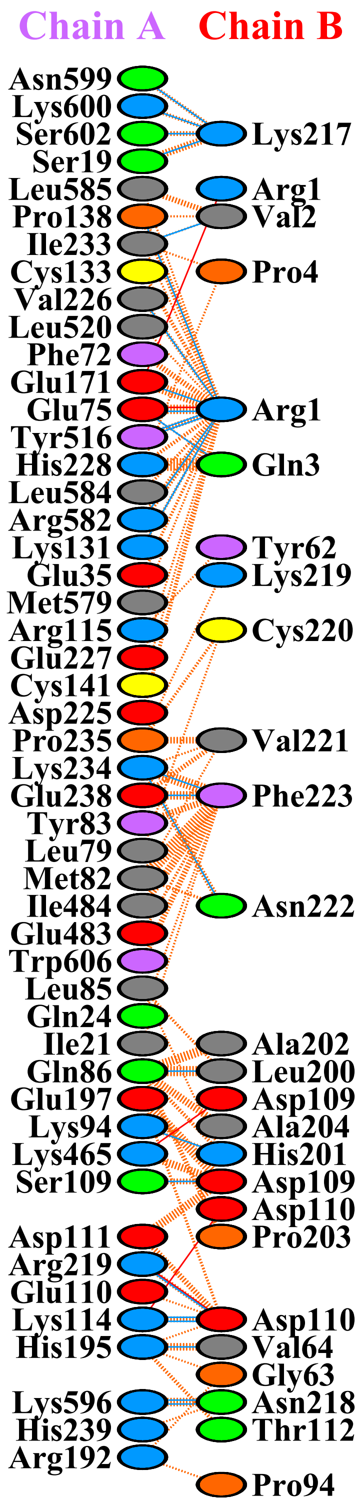


B

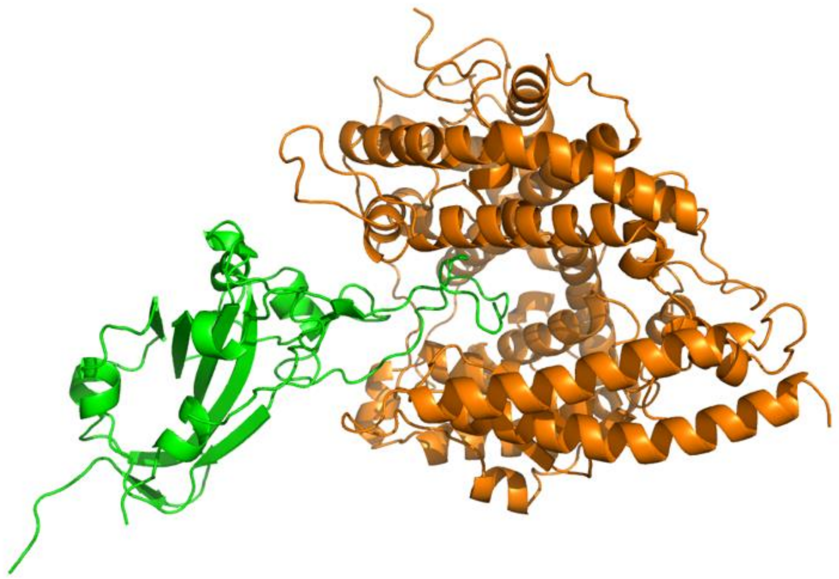

**Supplementary Figure:S2.X: Protein–protein docking representation of hACE2 and RBD of B.1.630 (EPI_ISL_3045385).** (A) The interaction representation, which includes hydrogen ( ), salt bridges ( ), and non-bonde[illegible] A ) interactions. Chain A represents hACE2 and chain B represents RBD . Residue colors: positive (blue): H,K,R; negative (red): D,E; neutral (green): S,T,N,Q; aliphatic (grey): A,V,L,I,M; aromatic (purple): F,Y,W; Pro & Gly (orange): P,G. (B) The binding interface of hACE2:RBD complex.

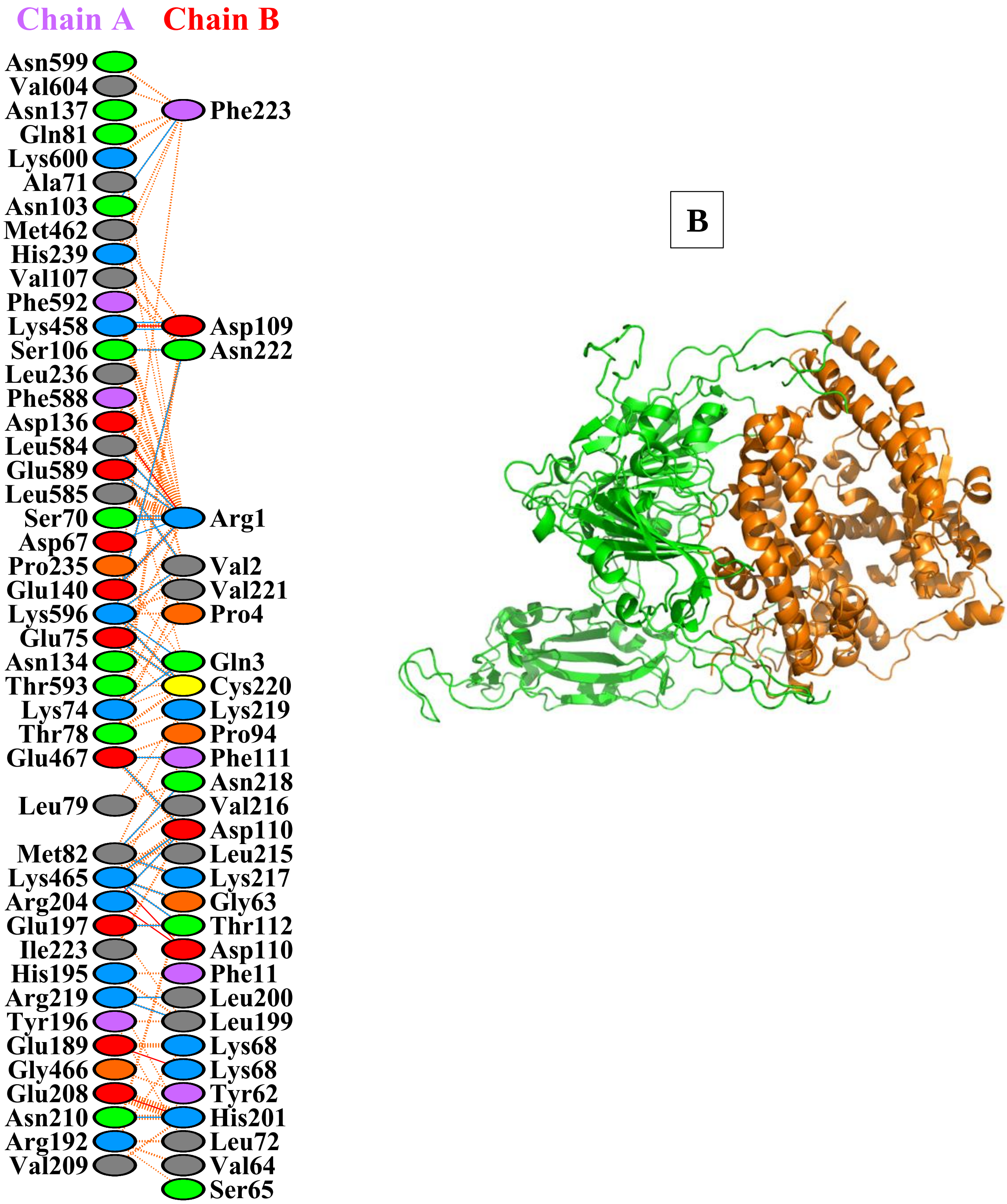


**Supplementary Figure:S2.Y: Protein–protein docking representation of hACE2 and RBD of C.1.2 (EPI_ISL_3447714) .** (A) The interaction representation, which includes hydrogen (▬▬), salt bridges (▬▬), and non-bonded (||||||) interactions. Chain A represents hACE2 and chain B represents RBD . Residue colors: positive (blue): H,K,R; negative (red): D,E; neutral (green): S,T,N,Q; aliphatic (grey): A,V,L,I,M; aromatic (purple): F,Y,W; Pro & Gly (orange): P,G. (B) The binding interface of hACE2:RBD complex.

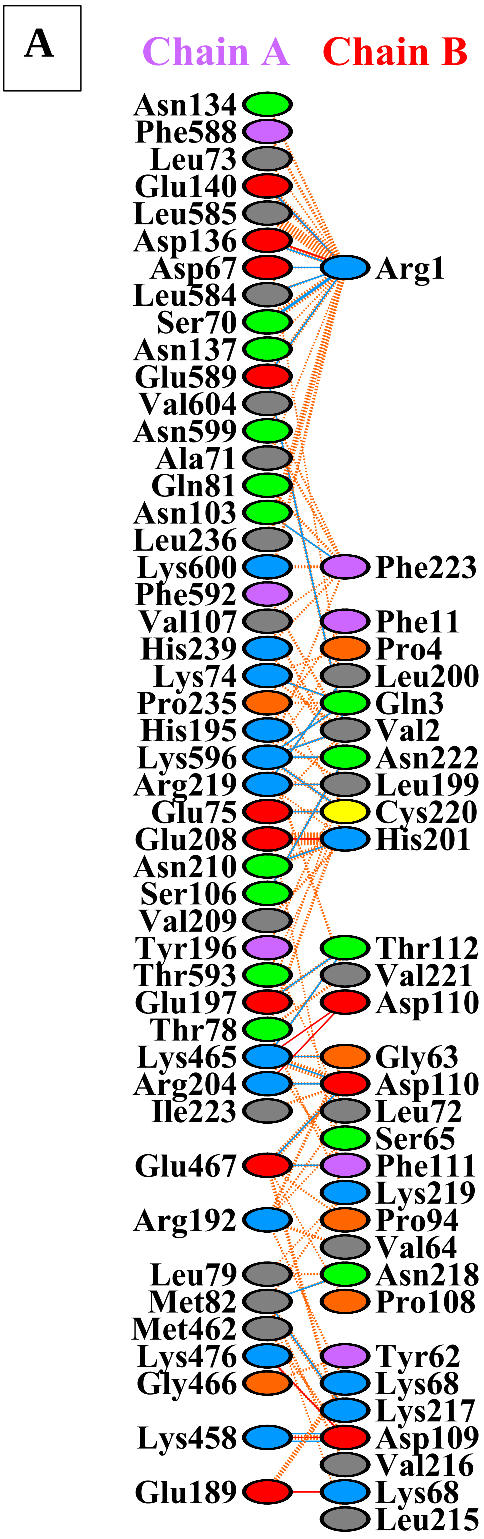


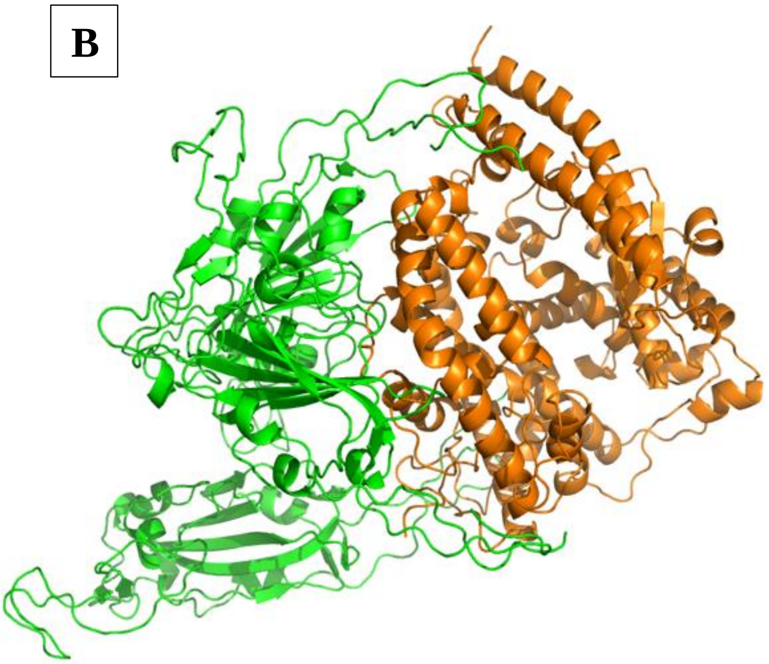


**Supplementary Figure S2.Z: Protein–protein docking representation of hACE2 and RBD of B.1.640 (EPI_ISL_5592661).** (A) The interaction representation, which includes hydrogen (blue line), salt bridges (red line), and non-bonded (orange dashed line) interactions. Chain A represents hACE2 and chain B represents RBD. Residue colors: positive (blue): H,K,R; negative (red): D,E; neutral (green): S,T,N,Q; aliphatic (grey): A,V,L,I,M; aromatic (purple): F,Y,W; Pro & Gly (orange): P,G. (B) The binding interface of hACE2:RBD complex.
